# The search for Lake Superior ciscoes as described by W. Koelz in 1929 yields a surprising discovery: Shortnose Cisco (*Coregonus reighardi*), a presumed extinct species

**DOI:** 10.1101/2025.04.20.649663

**Authors:** Owen T. Gorman, Amanda S. Ackiss, Thomas C. Pratt, Joshua F. Lyons, Renee E. Renauer-Bova, J. Vinnie Siegel, Ann J. Ropp, Yu-Chun Kao

**Affiliations:** U.S. Geological Survey, Great Lakes Science Center, Lake Superior Biological Station, WI, USA; U.S. Geological Survey, Great Lakes Science Center, Ann Arbor, MI, USA; Fisheries and Oceans Canada, Sault Ste. Marie, ON, Canada; U.S. Fish and Wildlife Service, La Crosse Fish and Wildlife Conservation Office, Onalaska, WI, USA

## Abstract

The Laurentian Great Lakes (LGL) formed following the retreat of the Wisconsin glaciation 15-9 ka and provided a vast volume of freshwater habitat for coregonines, which included a diverse endemic complex of ciscoes. However, anthropogenic impacts in the 19^th^ and 20^th^ centuries resulted in declines and losses of ciscoes, leaving only Lake Superior with a relatively intact complex that still dominates its prey fish community. In the 1920s, W. Koelz described six putative cisco species from Lake Superior: *Coregonus artedi*, *C. hoyi*, *C. kiyi*, *C. zenithicus, C. reighardi*, and *C. nigripinnis*. Of these, *artedi*, *hoyi* and *kiyi* remain extant and abundant in Lake Superior, *zenithicus* is considered extirpated throughout the LGL except in Lake Superior, and *reighardi* and *nigripinnis* are considered extinct and had been synonymized as *zenithicus* in Lake Superior. Using an integration of morphological and genetic approaches, we present results reaffirming the presence of *artedi*, *hoyi*, and *kiyi,* and new evidence for the presence of *reighardi*, a species thought to be extinct. Our analysis did not elucidate the presence of *zenithicus* and *nigripinnis*, leaving their status unresolved. The principal morphological characters that discriminate *artedi, hoyi, kiyi, and reighardi* are related to habitat and trophic (ecological) specialization and foster niche segregation: eye size, gill raker length, and gill raker spacing, the same as those that foster niche segregation in subarctic European whitefishes. The discovery of *reighardi* is expected to stimulate new interest in the development of conservation and restoration plans for rare and imperiled cisco species in the LGL and other freshwater lakes throughout the Northern Hemisphere, and our integrated morphological and genetic approach can benefit these efforts by providing insights for the evolution, diversity, and ecology of ciscoes.

## INTRODUCTION

The Earth is in the midst of its sixth mass extinction event, with human impacts over the past two centuries accelerating major ecosystem disruptions and rapid faunal declines (Ceballos et al. 2020; Cowie et al. 2022) and led to the adoption of the “Anthropocene” to describe the current geologic era^1^ (Crutzen 2002; Steffen et al. 2011; Malhi 2017). Although freshwater fishes represent one-fourth of all vertebrate species despite freshwater occupying < 1% of the Earth’s surface (Miranda and Miqueleiz 2021), they are subject to the highest extinction rates among vertebrates. This disturbing trend is exemplified by North American freshwater fishes, which are experiencing an extinction rate conservatively estimated to be 877 times greater than the background extinction rate of one extinction every 3 million years (Burkhead 2012). Fishes in the Laurentian Great Lakes (LGL) serve as a prime example of loss of biological diversity and ecosystem function. The precursors to the modern LGL formed following the retreat of the Wisconsin glaciation 15-9 ka (Dalton et al. 2020, Breckenridge 2007) and provided a vast volume of freshwater habitat that was colonized by ancestral ciscoes from glacial refugia (Bailey and Smith 1981; Mandrak and Crossman 1992; Turgeon and Bernatchez 2001a, b, 2003; Eshenroder and Jacobson 2020; Eshenroder et al. 2024). Prior to the 20^th^ century, the LGL contained a diverse endemic complex of ciscoes that dominated their fish communities (Koelz 1929, Eshenroder et al. 2016), but the anthropogenic impacts of European settlement in the 19^th^ and 20^th^ centuries resulted in widespread declines in diversity of ciscoes (Smith 1968, 1972). Currently, Lake Superior is the only LGL with a relatively intact complex of ciscoes, with *Coregonus artedi (ART)*, *C. hoyi (HOY)*, and *C. kiyi (KIY)* dominating the prey fish community (Bronte et al. 2003, Zimmerman and Krueger 2009, Gorman and Hoff 2009, Gorman 2012, Goldsworthy et al. 2024). The importance of ciscoes in the Lake Superior food web and their substantial contribution to ecosystem function (Kitchell et al. 2000, Gamble et al. 2011a, b, Matthias et al. 2021) has led to interest in restoring diminished and extirpated populations in other LGL: Huron, Erie, and Ontario (Weidel et al. 2022, Bunnell et al. 2023).

In the 1920s, Koelz (1929) described six cisco morphotypes as species from Lake Superior: *ART*, *HOY*, *KIY*, *C. zenithicus (ZEN), C. reighardi dymondi (REI)*, and *C. nigripinnis cyanopterus (NIG)^2^*. By the late 20^th^ century, *REI*, and *NIG* were considered extinct in the LGL, and *ZEN* is considered extirpated in the LGL except in Lake Superior (Todd 2003, Webb and Todd 1995, Eshenroder et al. 2016). Distinction of these three species was confounded when Lake Superior *REI* and *NIG* were synonymized with *ZEN* (Todd and Smith 1980, Robbins et al. 1991). An examination of the status of the synonymized taxon *ZEN* in Lake Superior in the 2000s showed that their abundance was greatly reduced compared to the 1920s (Gorman and Todd 2007, Gorman 2012, Pratt 2012), but consistent identification of these fish has been uncertain over time (Eshenroder et al. 2016). The challenge of confidently identifying ciscoes stems from high phenotypic variation in coregonines (ciscoes and whitefishes) that confounds species identification, and genetic tools in the past have been insufficient in resolving differences among species. This conundrum, termed the “coregonid problem” (Svärdson 1949) led to treating all ciscoes as a species complex and a single taxon, *C. artedi* (*sensu lato*) (McPhail and Lindsey 1970, Turgeon and Bernatchez 2003, Turgeon et al. 2016). New studies have shown *ART, HOY*, and *KIY* to be genetically distinct with the application of new genomic tools (e.g., Restriction-site Associated DNA sequencing (RADseq), transcriptomics, and whole genome sequencing (WGS); Ackiss et al. 2020, Lachance et al. 2021, Bernal et al. 2022, Backenstose et al. 2024), and Bernal et al. (2022) showed concordance in distinguishing these species with genomic, morphological, and stable isotope approaches.

Building on this body of work, our goal was to reexamine the contemporary diversity of ciscoes in Lake Superior as described by Koelz (1929) with an integrated application of morphometrics and genomics. Our genomic results provided evidence for four genetic clusters that aligned with four morphological species: *ART, HOY, KIY,* and, most notably, *REI*, a species thought to be extinct. We also identified the most informative morphometric characters for differentiating the four genetic clusters to aid in identification of species and provide insights as to their function in relation to ecological specialization and niche segregation. Based on a presentation of our recent discovery of *REI* in Lake Superior at the Joint Aquatic Sciences Meeting in Grand Rapids, Michigan in 2022, the American Fisheries Society-American Society of Ichthyologists and Herpetologists Names Committee decided to elevate the status of *REI* from extinct to extant and remove its synonymy with *ZEN* (Page et al. 2023). Our findings parallel the well-documented adaptive radiation of whitefish species in European post-glacial lakes (e.g., Præbel et al. 2013, Häkli et al. 2018, De-Kayne et al. 2022) and those studies provide a baseline and insights for understanding the post-glacial radiation of ciscoes in the LGL. Finally, our findings will increase understanding of the diversity, ecology, and evolution of ciscoes in the LGL; motivate additional genomic, morphological, and ecological investigations; and support ongoing conservation and restoration efforts.

## METHODS

To address our principal goal of reexamining the contemporary diversity of ciscoes in Lake Superior with an integrated application of morphology and genetics, our Methods address the following sequence of objectives:

1. Sample contemporary diversity of ciscoes in Lake Superior.
2. Identify the six ciscoes (*ART, HOY, KIY, ZEN, REI, NIG*) in Lake Superior as described by Koelz (1929) using observed and manually measured morphological characters (morphological species ID).
3. Genotype the morphological species and evaluate agreement between morphological species ID and clusters of cisco genotypes to assign species identities for genotypes (genetic species ID).
4. Create a database of size-corrected measures of morphometric characters from digital images of ciscoes with morphological species ID.
5. Identify the most informative morphometric characters that maximally discriminate genetic species.

### Morphological diversity of Lake Superior ciscoes

During 2003-2021, 2853 ciscoes were collected broadly across Lake Superior using bottom and midwater trawls and bottom-set small-mesh gillnets at 183 locations with depths ranging 20-200 m. Fish were frozen and stored at the U.S. Geological Survey Lake Superior Biological Station for later identification (ID) and morphological workup. Because of the high degree of morphological similarity and phenotypic variation among LGL ciscoes (Eshenroder et al. 2016), discrimination of species was not possible with a dichotomous key based on a few discrete diagnostic characters. Instead, we used a panel of definitive morphological characters based on those used by Koelz (1929) and Eshenroder et al. (2016) to ID the six Lake Superior ciscoes *(ART, HOY, KIY, ZEN, NIG, REI)* potentially present in Lake Superior (Table 1). Our ID of *NIG* was based on Koelz’s (1929) description for the Lake Michigan form because his description for Lake Superior *NIG* was very similar to that of *ZEN* and previous studies showed that Lake Superior *NIG* and *ZEN* could not be distinguished morphologically (Todd and Smith 1980) nor isotopically (Schmidt et al. 2009). We limited our analysis to adult fish (*n=*1883) to avoid confounding the effects of ontogeny on morphology and because species descriptions by Koelz (1929) and Eshenroder et al. (2016) are based on adult fish.

**Table 1.**
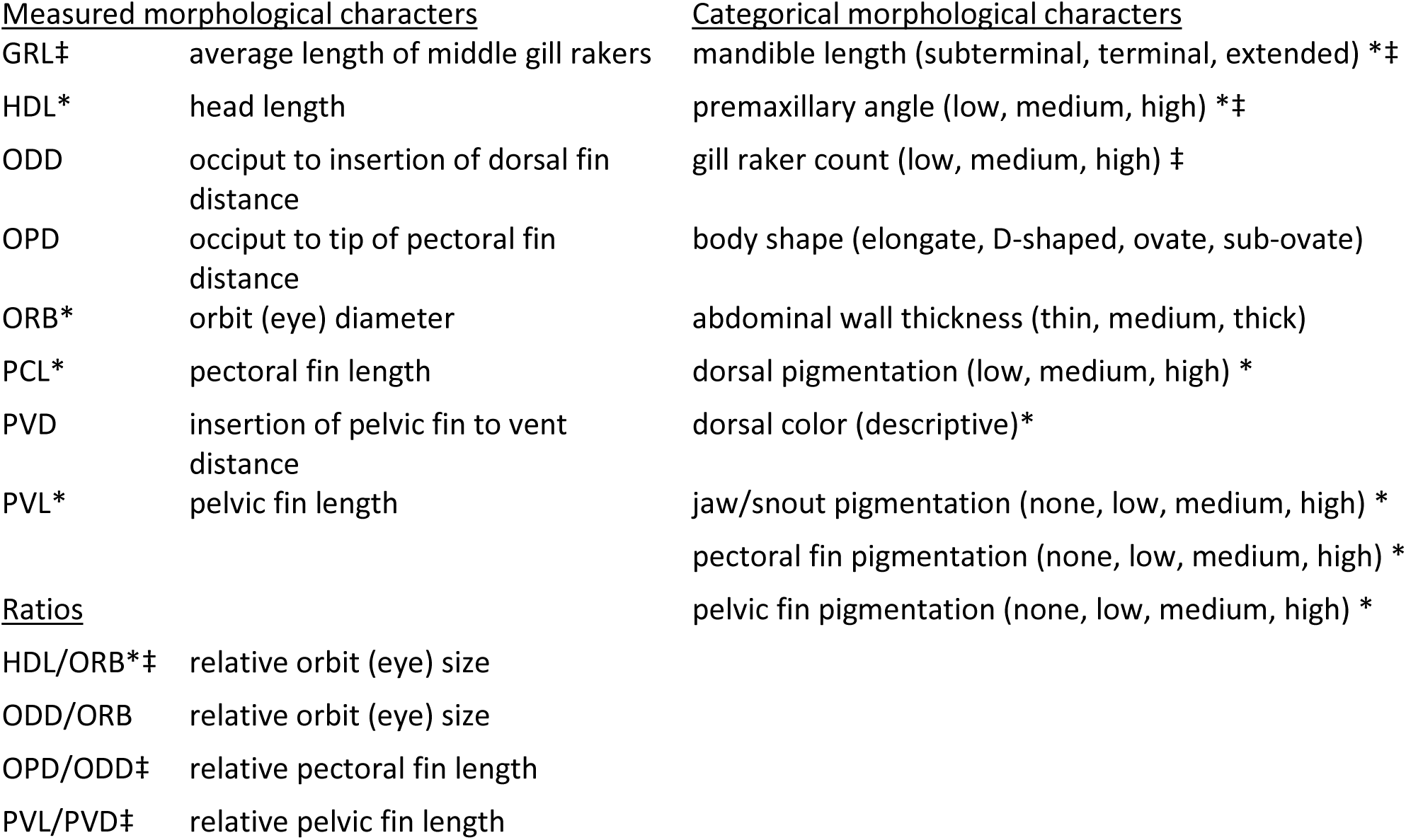
Morphological characters measured by hand, ratios of morphological characters, and categorical morphological characters assessed visually to identify ciscoes in a laboratory setting using the Cisco Score Card (Suppl. 3). Measured characters, ratios, and categorical morphological characters based on Koelz (1929) are indicated with “*”. OPD/ODD and PVL/PVD provide alternative indices of paired fin lengths compared to those used by Koelz (1929). Standard, fork, and total lengths (STL, FL, TL) were measured for all fish. Seven primary morphological characters used for ID of fish were double weighted and indicated with “‡”. All lengths and distances are in mm. A complete listing and definitions of morphological measures and counts is provided in Suppl. 2.

| <u>Measured morphological characters</u> |  | <u>Categorical morphological characters</u> |
| --- | --- | --- |
| GRL‡ | average length of middle gill rakers | mandible length (subterminal, terminal, extended) *‡ |
| HDL* | head length | premaxillary angle (low, medium, high) *‡ |
| ODD | occiput to insertion of dorsal fin distance | gill raker count (low, medium, high) ‡ |
| OPD | occiput to tip of pectoral fin distance | body shape (elongate, D-shaped, ovate, sub-ovate) |
| ORB* | orbit (eye) diameter | abdominal wall thickness (thin, medium, thick) |
| PCL* | pectoral fin length | dorsal pigmentation (low, medium, high) * |
| PVD | insertion of pelvic fin to vent distance | dorsal color (descriptive)* |
| PVL* | pelvic fin length | jaw/snout pigmentation (none, low, medium, high) * |
|  |  | pectoral fin pigmentation (none, low, medium, high) * |
|  |  | pelvic fin pigmentation (none, low, medium, high) * |
| <u>Ratios</u> |  |  |
| HDL/ORB*‡ | relative orbit (eye) size |  |
| ODD/ORB | relative orbit (eye) size |  |
| OPD/ODD‡ | relative pectoral fin length |  |
| PVL/PVD‡ | relative pelvic fin length |  |

To facilitate ID of ciscoes, a tabular scoring matrix was developed to assess the match of the character states of seven primary and five secondary definitive morphological characters for the six species (Table 1; Suppl. 3, Cisco Score Card). ID of each fish was conducted independently by two persons. For each fish, the identifier compared the expressed state of each morphological character with the definitive state for each of the six species and the relative match was scored as none, partial, or complete (0.0, 0.5, 1.0). Scores for the seven primary morphological characters were double weighted, secondary characters were single weighted, and scores were summed to create a composite score for each of the six species. IDs were decided by comparison of composite scores among the six cisco species: a composite score close to the maximum possible (19) indicates high conformation with one of the cisco species. In cases where multiple near ties occurred and summed scores were well below maximum, no identification was possible. The degree of match of morphological character states with one of the six cisco species and differentiation from other species determined the conformation score and level of confidence in the ID. Fish with conformation scores <6.0 were recorded as unidentified ciscoes (*n=*287), fish with scores ≥6.0 to <7.0 were given provisional IDs (*n=*244), and fish with scores ≥7.0 (*n=*1352) were considered putative IDs. Of fish with putative IDs, 39 had a maximum score of 10.0, representing a perfect conformation with one of the six cisco species. A final ID and conformation score were decided by consensus by the two identifiers. We emphasize that the conformation score is intended to reflect the degree of confidence in the ID, not an absolute ID.

Following manual measurements and consensus ID of fish, each fish with a conformation score ≥6.0 (provisional ID) was carefully positioned on a platform and photographed with a Canon® EOS-80D camera equipped with a Canon® EF-S 60 mm macro lens mounted to a copy stand to provide low-distortion, high-resolution digital images. After photographing each fish, gill arches were resected, mounted, and photographed in a similar manner. Fish were then dissected to determine sex and reproductive condition, stomachs removed for diet analysis, otoliths and scales harvested for aging analysis, and muscle plugs taken for genetic and stable isotope analysis. From digital images of each of the adult fish and associated gill raker arch, an extended set of 38 morphological characters (Suppl. 2) were digitally measured using SigmaScan Pro 5.0® software to provide an extensive morphometric dataset for each fish.

### Genetic diversity of Lake Superior ciscoes

Of the 1352 adult ciscoes with putative morphological IDs, 682 were selected for genetic analysis to optimize spatial coverage and maximal inclusion of rare ciscoes (*NIG, REI, ZEN*) and were genotyped with a GT-seq panel (Campbell et al. 2015) designed for species identification in Great Lakes ciscoes (Weidel et al. 2022). DNA extraction from frozen muscle tissue was performed with DNeasy Blood & Tissue kits (Qiagen Inc.) following manufacturer protocols, and GT-seq library preparation followed the protocols described in (Weidel et al. 2022). Libraries were sequenced on an Illumina MiSeq using paired-end (2 x 150 bp) v2 chemistry.

Sequence data for the 682 fish were processed and filtered in R (R Core Team 2021). GT-seq panel loci (single nucleotide polymorphisms, SNPs) were identified from demultiplexed, raw sequences using GTscore (v.1.3; github.com/gjmckinney/GTscore), and putatively contaminated samples were identified and removed (heterozygosity >0.25, contamination score >0.5). Iterative filtering (*sensu* O’Leary et al. 2018) removed badly genotyped loci (method = “loc”, threshold = 0.30) and individuals with high rates of missingness (method = “ind”, threshold = 0.5). This was followed by more stringent filters that removed loci with a genotyping rate less than 90% (method = “loc”, threshold = 0.9) and any individuals missing genotypes for more than 10% of remaining loci (method = “ind”, threshold = 0.9), and resulted in a subset of 602 genotypes. The locations where the 602 genotyped ciscoes were collected in Lake Superior are shown in Fig. 1.

**Figure 1.**
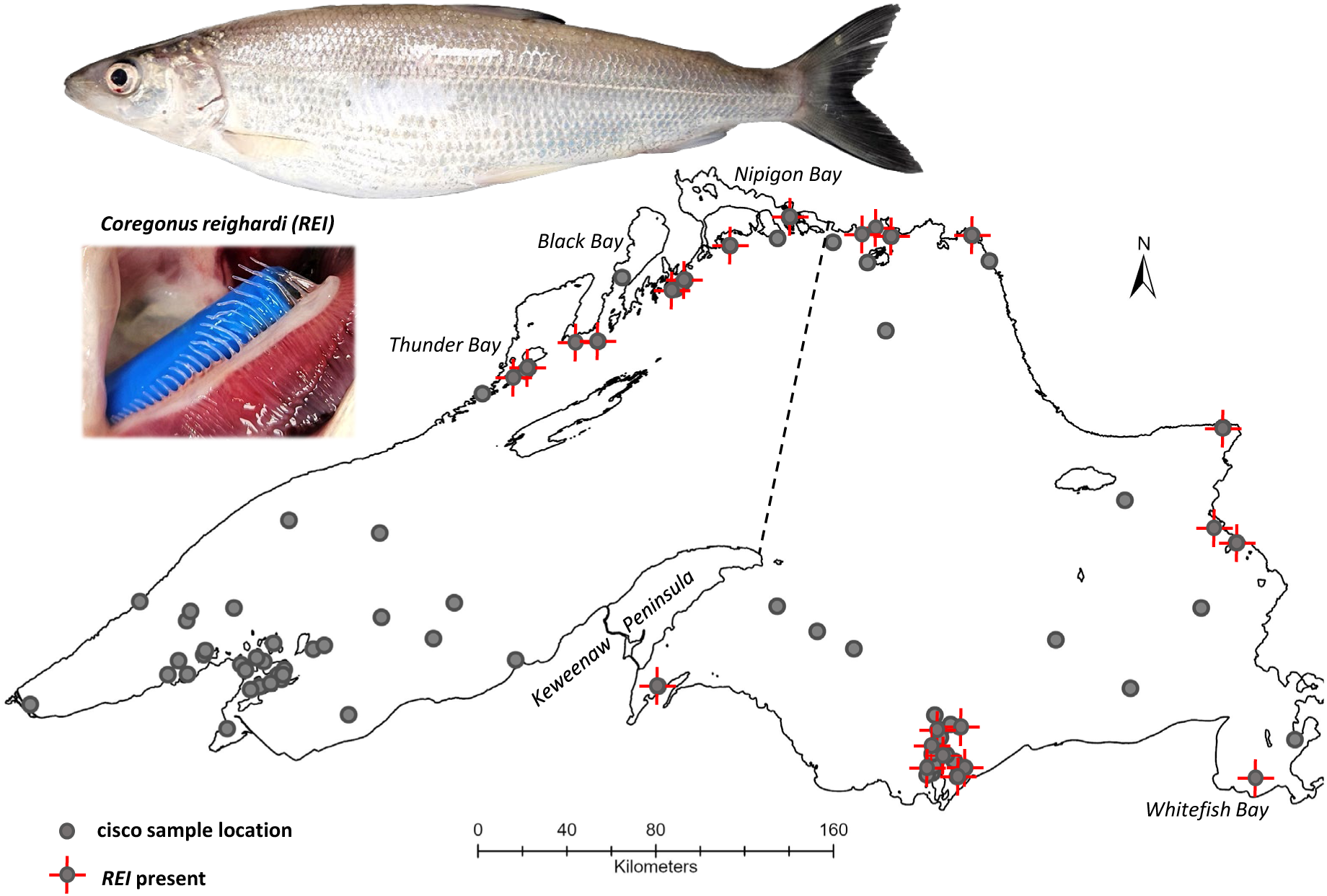
Lake Superior collection locations of genotyped ciscoes used in this study. Shown are 60 locations for 602 adult fish collected between 2006-2021 that conformed to Koelz’s (1929) descriptions as *Coregonus artedi (ART), C. hoyi (HOY), C. kiyi (KIY), C. zenithicus (ZEN), C. reighardi (REI),* and *C. nigripinnis (NIG).* Dashed line demarks the boundary of the eastern-most distribution of *REI* as described by Koelz (1929). Example of a typical adult *REI* (USGS specimen #2408273; 306 mm total length) is pictured and accompanied by a photo of the subject fish’s distinctively short, widely spaced gill rakers. A partially occluded 6.35 diameter mm dowel behind gill arch serves as reference for size.

Genetic diversity and differentiation in Lake Superior ciscoes were examined with two approaches. First, principal components analysis (PCA) was performed to examine species-associated allele frequency differences in the R package *adegenet* (Jombart 2008). Missing data were imputed for PCA using means. Second, Bayesian clustering was performed with STRUCTURE (Pritchard et al. 2000; Falush et al. 2003). Because genotyped individuals were composed of individuals morphologically identified as six different species (*ART*, *HOY*, *KIY*, *ZEN*, *REI*, *NIG*), STRUCTURE was run for inferred clusters (*K*) 1–6 with five replicates each. Runs consisted of a burn-in period of 50,000 MCMC iterations followed by an additional 100,000 steps under the assumption of population admixture and correlated allele frequencies without *a priori* assignment (USEPOPINFO = 0). STRUCTURE results were processed and visualized with the R package *pophelper* (Francis 2017). The optimal number of clusters (*K*) was evaluated with log likelihood [L(*K*)] and Evanno (Δ*K*; Evanno et al. 2005) methods, and plots of all *K*s were examined for additional biologically meaningful clusters beyond the optimal *K* as recommended for this type of analysis (Janes et al. 2017). STRUCTURE Q-scores for the best *K* were used to determine cluster membership using a threshold of Q ≥0.70 following similar cutoffs used for ciscoes (Ackiss et al. 2020, Lachance et al. 2021). Individuals that met this threshold were considered to have reliable genotypes and those that did not were designated as putative hybrids.

### Morphometrical analysis of genetic species

Following removal of 103 putative hybrids and 28 fish with missing morphometric data from the 602 genotyped adult ciscoes, 471 individuals were available for morphometric analysis that had both morphological and genetic IDs. Statistical analysis relied on morphometric measurements of 38 body and gill raker characters taken from digital images (described in Suppl. 2). Prior to statistical analysis, morphometric characters for each fish *i* were size corrected using Size Free Morphometric Ratios (*SFMR*) as per Eshenroder et al. (2021) using the equation

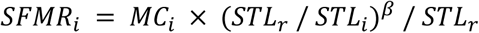

where *MC_i_* is the morphometric character of fish *i* in a genetic species, *STL_i_* is the standard length of fish *i*, *STL*_r_ is the geometric mean of the standard length of the species, and β is the allometric exponent that establishes a proportional relationship between *MC* to *STL*^β^. For each morphometric character, the parameter β was estimated as the regression coefficient of log(*STL*) in the model

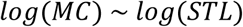

for each genetic species. This indicates that each genetic species has its own estimate of β for each morphometric character. We did not attempt to estimate a common β for any morphological character across genetic species, as their genetic distinctions make it unreasonable to assume identical allometric growth patterns. While this method is effective at removing size effects from our analysis, its use of group-based correlations compresses within-group variation, exaggerating divergence between groups (genetic species). Thus, a duplicate analysis was conducted using a simple arithmetic ratio of the morphometric character to STL as a contrast. While simple ratios have been discounted as effective size adjustment tools (Albrecht et al. 1993), this measure adds certainty to the inter-species divergence examined in this study.

Statistical analysis of morphometric data excluded putative hybrids and individuals with missing data. We used two approaches for PCA of morphometric data. First, we used Quadratic Discriminant Analysis (QDA; Venables and Ripley, 2002) to select the most informative characters among an extended set of 38 digital morphometric characters (Suppl. 2) for distinguishing cisco species based on genetic identity. To select the most informative morphometric characters to maximize discrimination of genetic species, we used a stepwise, cross-validation algorithm to select the most informative QDA model (Weihs et al. 2005). We calculated Variable Inflation Factor (VIF) and Pearson correlation coefficient (r) for characters in the final model to evaluate multicollinearity. In the PCA, groups based on genetic identity were visualized in morphological space by plotting 95% concentration ellipses on the plane of the first two principal components (PC1 and PC2; Kassambara 2016). Second, we used cluster analysis (k-means++; Smith 2020) to identify intrinsic morphological groups in our final PCA model and plotted 95% ellipses over PC1 and PC2. The optimal number of clusters was determined using average silhouette methodology. These two approaches allowed us to compare visual clustering of species by morphometrics vs. genetics.

## RESULTS

### Genetic and morphological diversity of Lake Superior ciscoes

After iterative filtering, our dataset consisted of 602 genotyped adult ciscoes at 993 SNPs. Individual genotype missingness ranged from 0-0.10 (average 0.01). Ciscoes in our filtered dataset were putatively identified with morphology as six species: *ART*, *n*=71; *HOY*, *n*=203; *KIY n*=163; *ZEN*, *n*=112; *REI*, *n*=30; *NIG*, *n*=23 (Table 2). PCA indicated that these ciscoes grouped into four primary genetic clusters (Fig. 2a, b). Three clusters aligned with individuals that had been morphologically identified as *ART*, *HOY*, and *KIY* following previously detected discrimination of these species (Ackiss et al 2020, Lachance et al. 2021, Bernal et al. 2022). The fourth cluster was comprised of all but one the individuals morphologically identified as *REI* and a small subset of individuals morphologically identified as *ZEN*. STRUCTURE results were consistent with PCA (Fig. 2c; Suppl. 1, Fig. 1). Log-likelihood analysis suggested that the optimal number of clusters (*K*) was 4, while the Evanno analysis suggested that optimal *K* = 2. Q-score plots indicated that *K* = 2 discriminates *ART*/*HOY* clusters from *KIY*/*REI* clusters, *K* = 3 further discriminates the *ART* cluster from the *HOY* cluster, and *K* = 4 discriminates all four of these species from each other (Q-scores for K = 4 are reported in Suppl. 1, Table 1). *K* ≥ 5 forced inferred clustering within one of the four primary clusters and did not reveal additional biologically meaningful clustering. The consensus morphological species that aligned with specific genetic clusters were designated the genetic species for that cluster. Fish associated with a cluster but whose morphological species ID did not match the genetic species ID of that cluster were judged mis-identified morphologically and assigned the genetic ID for that cluster. In both PCA and STRUCTURE analyses, individuals identified as *ZEN* and *NIG* were distributed across clusters or intermediately among primary clusters (Fig. 2a, b). Finally, fish with Q-scores <0.70 were not associated with any genetic clusters and designated putative hybrids (*n*=103) following thresholds applied in Ackiss et al. (2020) and Lachance et al. (2021).

**Figure 2.**
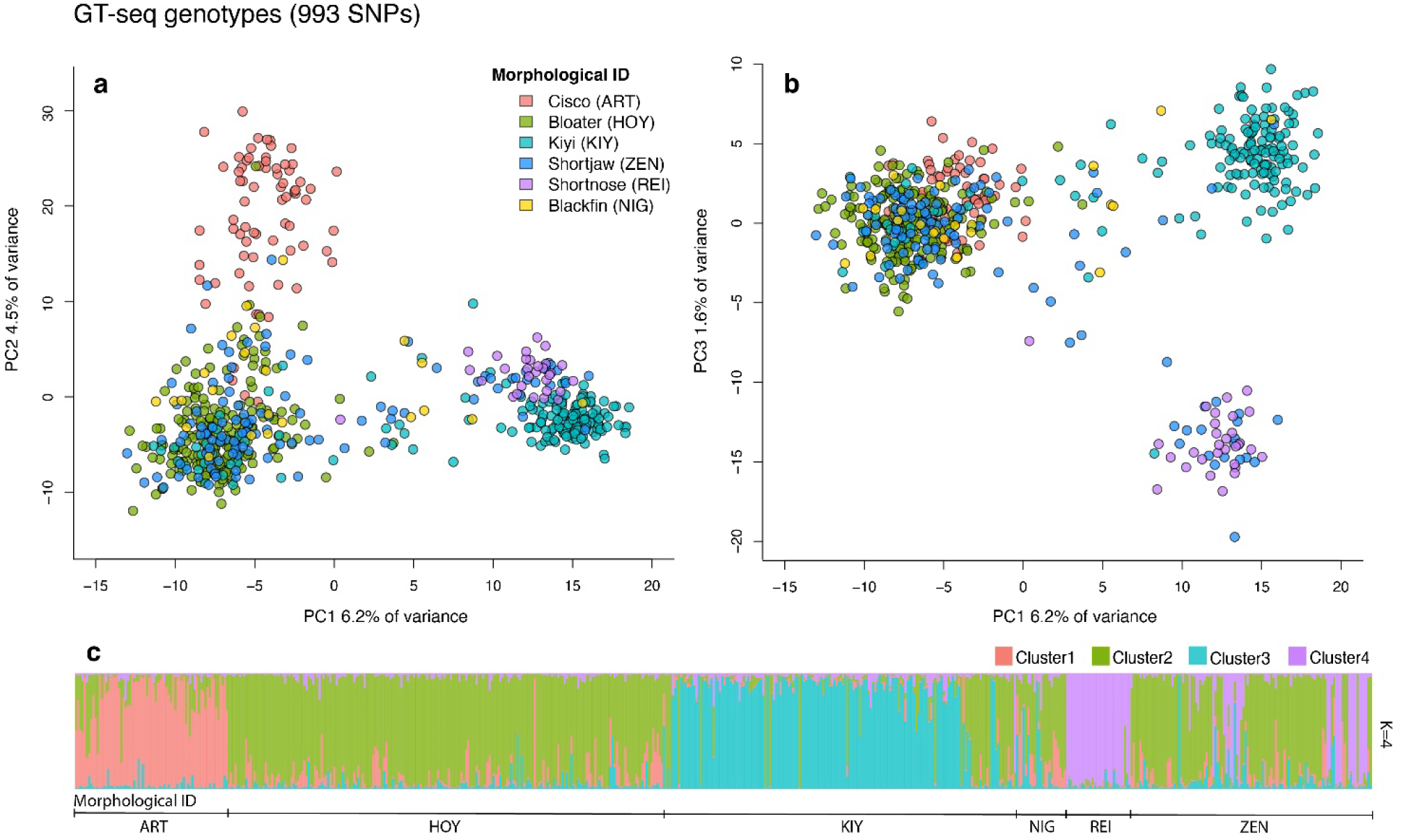
Principal components analysis and STRUCTURE plot based on 993 SNPs for 602 genotypes showing support for 4 genetic clusters in the Lake Superior ciscoes we genotyped. The genetic PCA scores are color-coded by species identified morphologically and serve to visualize the concordance between genetic clusters and species identification. The first two principal components (PC1 and PC2) discriminate *ART*, *HOY* and *KIY*/*REI* clusters (a). The third principal component (PC3) discriminates *KIY* from *REI* (b). Q-scores in the STRUCTURE plot (c) show that the 4 genetic clusters correspond with morphological IDs of ciscoes as follows: 1 = *ART*, 2 = *HOY*, 3 = *KIY*, 4 = *REI,* and NIG and ZEN do not correspond with individual genetic clusters. STRUCTURE plots of Q-scores for 2 to 6 genetic clusters are provided in Suppl. 1, Fig. 1 to show that 4 is the optimal number of genetic clusters supported by the data. Q-scores for K = 4 are reported in Suppl. 1, Table 1.

**Table 2.** Comparison of morphological and genetic identification of ciscoes that were genotyped. Of 602 ciscoes genotyped, 103 were putative hybrids, yielding 499 fish with reliable genotypes. Only four species were recognized by genetics (*ART*, *HOY*, *KIY*, *REI*) while two additional species (*ZEN*, *NIG*) were identified with morphology. Overall match of morphological species *ART*, *HOY*, *KIY*, and *REI* with non-hybrid genetic counterparts was 93.1%. Differences in agreement of species identification by morphology (italicized values, n = 403) vs. genetics (underlined values, n = 375) were not significant (*p* = 0.676, Chi-square Test of Independence = 1.529; 3 *df*). The distribution of putative hybrids among the six morphological IDs was significantly different than the expected, 17.1% overall (*p* < 0.001, Chi-Square Goodness of Fit Test = 34.926; 5 *df*).

| Genetic ID | Morph ID |  |  |  | *ZEN* | *NIG* | Genetic Total |
| --- | --- | --- | --- | --- | --- | --- | --- |
|  | *ART* | *HOY* | *KIY* | *REI* |  |  |  |
| *ART* | <u>50</u> | 1 |  |  |  |  | 51 |
| *HOY* | 1 | <u>175</u> | 25 |  | 64 | 9 | 274 |
| *KIY* |  |  | <u>121</u> |  | 2 | 2 | 125 |
| *REI* |  |  | 1 | <u>29</u> | 19 |  | 49 |
| putative hybrid | 20 | 27 | 16 | 1 | 27 | 12 | 103 |
| Morphologic Total | 71 | 203 | 163 | 30 | 112 | 23 | 602 |
| Total, less hybrids | 51 | 176 | 147 | 29 | 85 | 11 | 499 |
| % ID match | 98.0% | 99.4% | 82.3% | 100.0% | — | — | — |
| % hybrid | 28.2% | 13.3% | 9.8% | 3.3% | 24.1% | 52.2% | 17.1% |

After filtering putative hybrids and reassignment of morphologically mis-identified ciscoes, the remaining 499 ciscoes aligned with four species: *ART*, *n*=51; *HOY*, *n*=274; *KIY n*=125; *REI*, *n*=49 (Table 2). Morphological IDs of the four species (*n*=403) showed high agreement with matching genetic IDs (82.3-100.0%, 93.1% overall) and was not significantly different from being equivalent (Table 2). However, the morphological IDs of *ZEN* and *NIG* (*n*=96) did not correspond with matched genetic clusters and their IDs were dispersed among genetic clusters of *HOY*, *KIY*, and *REI*. The frequency of putative hybrids varied significantly from expected values among the morphological species, with *ART*, *ZEN*, and *NIG* having the highest hybrid proportions (28.2%, 24.1%, 52.2%, respectively) and *HOY*, *KIY*, and *REI* having the lowest proportions (13.3%, 9.8%, 3.3%, respectively; Table 2).

### Resolution of genetic species with morphometrics

Following removal of 103 putative hybrids and 28 fish with missing morphometric data from the 602 genotyped adult ciscoes, 471 individuals were available for morphometric analysis. The most supported model from the QDA of morphometric data using Genetic ID as the grouping variable included the following morphometric characters, in order of importance: orbital length (ORB), mean length of the three gill rakers nearest the gill arch inflection point (GRML), and gill raker spacing (GRS). This model was robust and, when trained using 70% of the individuals, species assignment to the remaining 30% was 98.7% accurate. VIF values among predictors were low, with a maximum value of 2.19 for ORB, well below a moderate value of 5. Pearson correlation coefficients were also low except for ORB ∼ GRS (r = 0.733). This value can indicate multicollinearity; however, these characters are not morphometrically correlated (i.e., both are not on the gill arch and have different functionalities) and were not excluded from the model.

The three morphometric characters (ORB, GRML, GRS) explained 91.6% of the variance in PC scores (Fig. 3) and 97.5% of the fish were correctly assigned to their respective genetic species clusters (*ART*, *HOY*, *KIY*, *REI*; Fig. 4b). Dimension 1 of the PCA plot segregated species clusters by ORB and GRS and dimension 2 segregated clusters by GRML (Fig. 3). *ART* is distinguished by the smallest ORB and GRS, while *KIY* has the largest ORB and GRS. *HOY* has the longest GRML, but ORB and GRS are intermediate, and *REI* differs from *HOY* by having the smallest GRML (Fig. 4b). Clustering analysis of the morphological PC scores (k-means++) showed that intrinsic clustering of morphological data matched clustering by genetic species 97.6-100.0%, and four clusters were found to be optimal via the average silhouette method (Fig. 4c). PCA plots allowed comparison of clustering by genetic ID and morphology. Clusters of genetic species ID (Fig. 4b) aligned closely with k-means++ clusters of morphometric data: clusters 1 and 4 matched 100% with *REI* and *ART*, respectively; cluster 2 matched 96.7% with *KIY*; and cluster 3 matched 98.2% with *HOY* (Fig. 4d). As a contrast, clustering of the genetic species in a PCA based on simple arithmetic ratios of ORB, GRML and GRS to STL resolved just 42.5% of the variance in PC scores (Fig. 4a). In this PCA plot, *ART* is fully segregated from the other species by having the smallest ORB and GRS; *HOY* and *REI* overlap extensively by sharing intermediate ORB, GRML and GRS; and *KIY* is partially segregated from the others by larger ORB and GRS.

**Figure 3.**
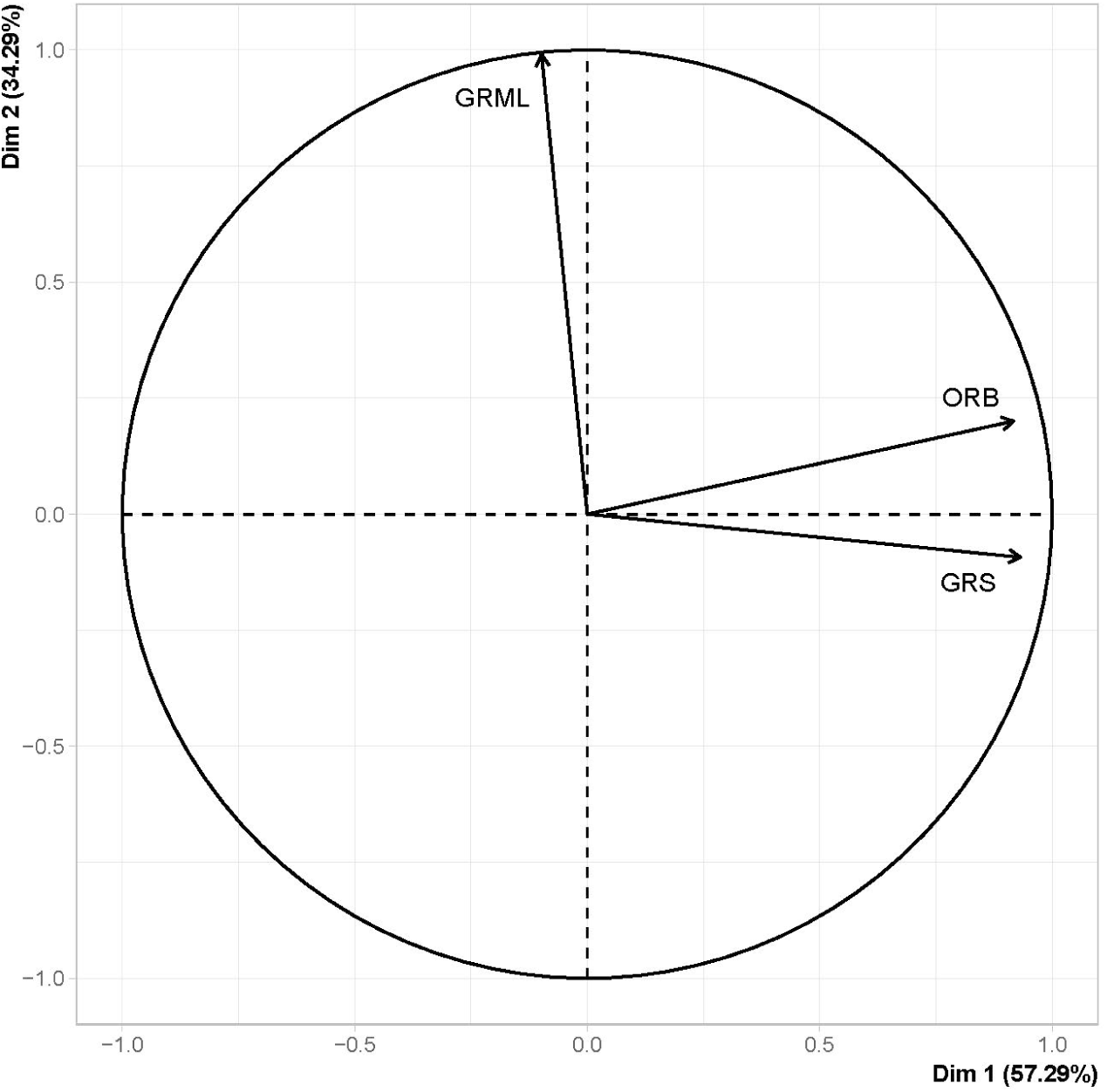
Relative loadings of the three most informative variables (morphological characters) for Principal Component Analysis (PCA) of size-corrected morphometric data. PC1 (Dim 1) is defined by ORB and GRS and maximizes separation of *ART* from other ciscoes by having the smallest relative eye size and narrowest gill raker spacing. PC1 secondarily distinguishes *KIY* from *BLO* and *REI* by having the largest relative eye size and widest gill raker spacing. PC2 (Dim 2) is defined largely by GRLM and distinguishes *REI* from the other ciscoes by having the shortest gill rakers. These three morphological characters explain 91.6% of the variance in PC scores in two axes.

**Figure 4.**
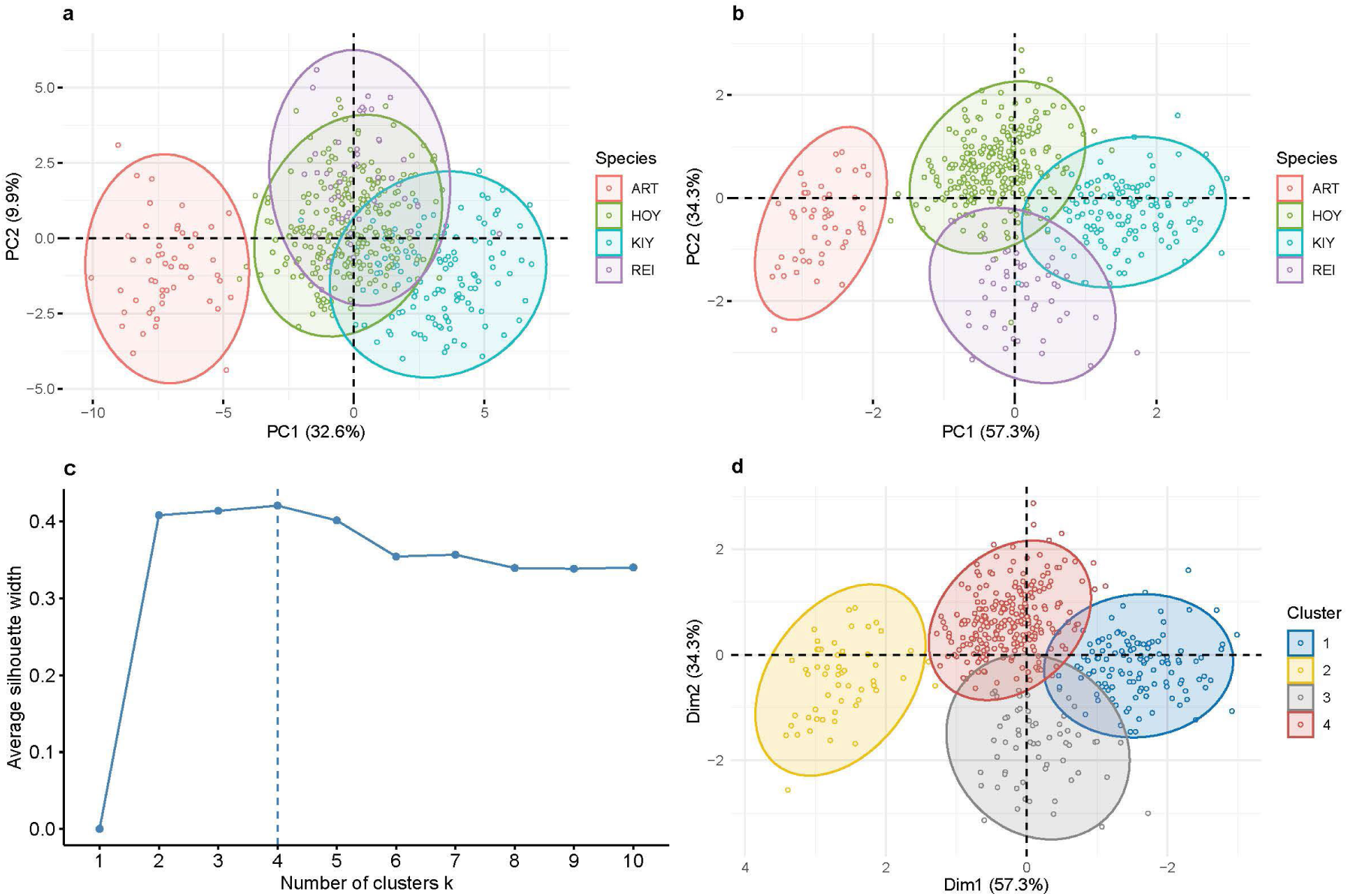
Principal components analysis (PCA) of 471 adult ciscoes with genetic ID using the three most discriminatory morphological characters: orbital length (ORB), maximum length of the three gill rakers nearest the gill arch inflection point (GRLM), and gill arch length/total gill rakers (gill raker spacing, GRS). Shown are results of PCA of morphometric variation for four species identified by genetics with simple size correction (a), allometric size correction (Eshenroder et al. 2021) (b), optimal number of clusters (4) based on average silhouette width is indicated in (c), and k-means++ clustering of morphometric PCA scores without regard to species ID (d). 95% concentration ellipses are colored by genetic identification (a, b), and by cluster number (d). Excluded from the analysis were putative hybrids and fish with missing morphological data (n=131).

## DISCUSSION

### Overview of key findings

The primary goal of our study was to sample the contemporary morphological and genetic diversity of ciscoes in Lake Superior and explore evidence for the presence of the six ciscoes described by Koelz (1929) in the 1920s. However, our PCA and cluster analysis of the genotyped fish yielded only four genetic clusters that aligned with four of six morphological species: *ART, HOY, KIY, REI*. Three of these species, *ART, HOY,* and *KIY,* have repeatably been shown to be both morphologically and genetically distinct (Ackiss et al. 2020; Bernal et al. 2022). Notably, this is the first genetic study to detect a fourth genetic cluster in Lake Superior; one comprised of morphologically identified *REI*, which was thought to be extinct in the LGL (Webb and Todd 1995) and never existed in Lake Superior (Todd and Smith 1980, Todd 2003, Eshenroder et al. 2016). The high level of agreement of morphological IDs of *ART, HOY, KIY, REI* with genetic IDs (93.1% overall; Table 1) demonstrates that there is a concordant morphological and genetic basis for distinguishing these species, a finding reinforced by distinct clustering of species using QDA to select the most informative morphometric characters to discriminate genetic species (97.5% overall; Fig. 4b). Moreover, the high level of match between our morphological and genetic IDs indicates that our methodology of morphological ID based on Koelz’s (1929) descriptions yields reliable results for the four species. Thus, we demonstrate that *ART, HOY, KIY,* and *REI* can be reliably identified in the field or laboratory based on a panel of diagnostic morphological characters (Suppl. 3, Cisco Score Card).

Two morphological species, *ZEN* and *NIG*, did not form distinct genetic clusters and those with reliable genotypes were identified as belonging to one of three genetic species, *HOY, KIY, REI.* Focusing on *ZEN* as an example, of 85 fish morphologically ID as *ZEN*, 64 (75.3%) matched genetic *HOY* and 19 (22.4%) matched genetic *REI*, suggesting that the morphology defined for *ZEN* overlaps with that of *HOY* and *REI*, or the observed morphologies of *HOY* and *REI* are more variable than indicated in Koelz’s (1929) descriptions. The morphology of *HOY* also appears to overlap with *KIY*, as 25 of 147 (17%) of fish ID morphologically as *KIY* were ID as genetic *HOY*. All total, 99 of 274 (36%) of fish genetically ID as *HOY* did not match morphological ID as *HOY*, but for fish that were morphologically ID as *HOY*, the agreement with genetic ID was 99.4%. This suggests that Koelz’s morphological description of *HOY* is useful for reliable ID of “classical” *HOY*, but more than a third of the genetic *HOY* were ID as species with different morphologies. Our results demonstrate that *HOY* exhibits such variable morphology that making reliable IDs for all *HOY* based on the idealized morphology we used is uncertain. However, our PCA and cluster analysis showed that based on the three most informative morphological characters (ORB, GRML, GRS), 88.8% of *HOY* can be discriminated from other ciscoes and agreement between intrinsic morphological clustering and genetic species assignment was 98.2% (Fig. 3c). These findings will be helpful in selecting the most discriminating morphological characters to make more accurate identification of ciscoes in the field and laboratory.

The uneven apportioning of putative hybrids among the six morphological species presented an unexpected potential relationship between genetic and morphological IDs: 24.1% of fish morphologically ID as *ZEN*, 52.2% ID as *NIG,* and 28.2% ID as *ART* were putative hybrids (Table 1). If the proportions of putative hybrids among morphological species were similar, this could be explained as genetic ambiguity arising from degraded DNA, but because the number of putative hybrids among the morphological species was significantly different from expected (17.1%), the differences may reflect hybridization or genotype differences that do not align with the four genetic clusters (*ART*, *HOY*, *KIY*, *REI*). The high proportion of non-hybrid *ZEN* and *NIG* identified as genetic *HOY* and the higher than expected number of putative hybrids in *ZEN* and *NIG* may reflect morphological variation in what constitutes genetic *HOY* and that there may be a correlation between morphological variants and putative hybrids. By contrast, the greater than expected number of putative hybrids in *ART* does not appear to be related to issues of ID as 50 of 52 morphologically ID fish matched genetic ID, suggesting that both putative hybrids and non-hybrids exhibited similar morphologies.

### Rediscovery of REI and uncertainty of ZEN and NIG

The taxonomic history of *REI* is complex and how it disappeared from Lake Superior is a product of the oftentimes convoluted process of scientific investigation. Koelz (1924) described *REI* from specimens collected in Lake Michigan and while additional specimens collected from lakes Nipigon, Superior, and Ontario were available, they were described later in Koelz (1929). Much later, Todd and Smith (1980) synonymized Lake Superior *REI* and *NIG* with *ZEN* based on exhibiting similar morphologies, which in effect asserted that *REI* and *NIG* never existed in Lake Superior. However, *REI* continued to be recognized as a valid species in lakes Michigan, Huron, and Ontario and their numbers dwindled rapidly after 1964 (Webb and Todd 1995). The last records of *REI* in lakes Ontario, Michigan, and Huron were 1964, 1974, and 1985, respectively, and afterwards Webb and Todd (1995) considered *REI* extinct throughout its known range. Given this history, we were surprised to discover *REI* to be not only present in Lake Superior but widespread in distribution.

Records of *ZEN* in Lake Superior have a long history dating back to the early 20^th^ century. This species was described from specimens collected in Lake Superior by Jordan and Evermann (1909), and Koelz (1929) reported that *ZEN* had a wide distribution in Lake Superior and was the most abundant deepwater cisco in the commercial chub fishery. While Koelz (1929) observed that both *ZEN* and *NIG* were abundant in the early 20^th^ century, he found *REI* to be relatively rare and only present west of line from the outlet of Nipigon Bay south to the Keweenaw peninsula (Fig. 1). Gorman and Todd (2007) provided a detailed review of commercial and assessment gillnet records that chronicle the decline of *ZEN* (including *NIG* and *REI*) during the 20^th^ century, trending from an initial high abundance to rarity by the 1970s, and near absence by the late 1980s (Gorman 2012). The acceptance of the synonymy of *REI* and *NIG* with *ZEN* was challenged by the discovery of large isotopic differences between *REI* and *ZEN* in early 20^th^ century Lake Superior (Schmidt et al. 2009). Moreover, Schmidt et al. (2009) showed that *ZEN* exhibited a high degree of isotopic separation from other Lake Superior ciscoes (except *NIG*). Building on this work, Blanke et al. (2018) examined amino acid-specific (A-AS) isotope signatures of *ZEN*, *NIG*, and *REI* from Lake Superior and showed that these species exhibited discrete A-AS trophic positions, indicative of a high degree of trophic niche segregation among these ciscoes. These findings suggest that in the early 20^th^ century, *ZEN* and *REI* functioned as distinct species with distinct trophic ecologies in Lake Superior. That their isotopic positions closely matched that of *ZEN* and *REI* in early 20^th^ century lakes Michigan and Huron (Schmidt et al. 2011), as well as their trophic positions in early 20^th^ century Lake Michigan (Blanke et al. 2018), suggests that *ZEN and REI* had a broad distribution across the LGL. Moreover, the concordance of trophic positions of *ART, HOY ZEN, KIY, REI*, and *NIG* (in part) in lakes Superior, Michigan and Huron and their similar patterns of trophic segregation in lakes Superior and Michigan in the early 20^th^ century suggests that these species had similar trophic niches and ecologies in the LGL (Schmidt et al. 2011, Blanke et al. 2018).

Given this historical and ecological background, we were surprised not to find *ZEN* as a distinct morphological and genetic species in our study. In the early 20^th^ century, the Lake Superior fish community and ecosystem was in the early stages of anthropogenic impacts stemming from European settlement that accelerated in the late 19^th^ century (Lawrie and Rahrer 1972). Environmental impacts continued during the 20^th^ century along with increasing commercial fishing (Koelz 1926; Smith 1968, 1972; Lawrie 1978) and the spread of non-native species rainbow smelt (*Osmerus mordax*) and sea lamprey (*Petromyzon marinus*) (Smith 1972, Christie 1974, Lawrie 1978). In response to these perturbations, the Lake Superior ecosystem and fish community changed accordingly, along with a decline in abundance of ciscoes, especially impacting *ZEN*, *NIG*, and *REI* (Lawrie and Rahrer 1972, Smith 1972, Christie 1974, Lawrie 1978, Bronte et al 2003, Gorman and Todd 2007, Gorman 2012). With the recovery of lake trout (*Salvelinus namaycush*) came the recovery of *ART*, *HOY*, and *KIY*, but not *ZEN* – including synonymized taxa *NIG*, *REI* (Bronte et al 2003, Gorman and Hoff 2009, Gorman 2012). These fish community changes can be seen in a comparison of isotopically inferred trophic niches in early 20^th^ vs early 21^st^ century Lake Superior: the niche breadth of forage fishes was slightly reduced and shifted (Blanke et al. 2018). The same comparison with Lake Michigan showed a greatly reduced niche breadth and a loss of all ciscoes except *HOY*, indicative of an impacted ecosystem with an impaired and simplified food web (Blanke et al. 2018). These findings suggest that despite the absence of *ZEN*, *NIG*, and *REI* in Blanke et al.’s (2018) samples, the Lake Superior ecosystem appears capable of supporting a more diverse assemblage of ciscoes than other LGL. The discovery of the presence of *REI* in our surveys demonstrates that Lake Superior has maintained the capability of supporting a more diverse assemblage of ciscoes. The relatively high-functioning state of the Lake Superior ecosystem demonstrated by Blanke et al. (2018) leaves open the possibility that *ZEN* may still be present, but that will not be resolved without further sampling.

Like *ZEN*, resolution of the status of *NIG* in Lake Superior has proven to be elusive. We concur with Todd and Smith (1980) that Lake Superior *NIG* as described and named by Koelz (1929; *C. nigripinnis cyanopterus*) likely represented large, old *ZEN* and its synonymy with *ZEN* was supported by Schmidt et al. (2009), showing that they had similar isotopic signatures. However, we observed ciscoes in Lake Superior that more closely aligned with *NIG* as described and named for lakes Michigan and Huron by Koelz (1929; *C. nigripinnis nigripinnis*), so we used Koelz’s description for Lake Michigan *NIG* as a basis for morphological ID of *NIG* in Lake Superior. We found *NIG* to be a rare morph in our survey of Lake Superior ciscoes, yielding 23 fish of which 12 were putative hybrids, 9 were genetic *HOY* and 2 were genetic *KIY* (Table 2), a result that leaves the status of *NIG* unresolved in Lake Superior.

### Ecological and evolutionary implications of morphological and genetic differentiation in coregonines

Morphological variation and lack of clear morphological differences among ciscoes has been problematic for identification and execution of ecological studies, the “coregonid problem” as identified by Svärdson (1949). Ackiss et al. (2020) provided an updated review of the coregonine problem considering the application of new genomic tools that have allowed discrimination of *ART*, *HOY*, and *KIY* (Ackiss et al. 2020, Lachance et al. 2021, Bernal et al. 2022, and Backenstose et al. 2024). Our work builds on these studies by integrating morphometrics with genomics to identify morphological characters that are associated with genetic differences. Our goal was to accurately discriminate the six ciscoes morphologically as described by Koelz (1929) for Lake Superior, particularly the rare ciscoes *ZEN*, *NIG*, and *REI*. The high level of morphological overlap among *ZEN, REI* and *NIG* as presented by Todd and Smith (1980) underscores the morphological ambiguity of these species in Lake Superior and led to *NIG* and *REI* being synonymized with *ZEN*. Despite the apparent lack of morphological characters to clearly distinguish *ZEN*, *NIG*, and *REI*, we found that *REI* could be distinguished as having the lowest count, shortest length, and greatest spacing of their gill rakers. Our finding of the distinctive morphology, genetics, and wide distribution of *REI* in Lake Superior (Figs. 1-3) is surprising given that this species was not considered to exist in Lake Superior (Todd and Smith 1980) and that it was considered extinct throughout its known range (Webb and Todd 1995). Our discovery of *REI* demonstrates that this species has been present in Lake Superior since its description by Koelz (1924, 1929). We now recognize that the synonymy of *NIG* and *REI* with *ZEN* obscured understanding geographic variation, distribution, and population trends among these species., e.g., Gorman and Todd (2007), Gorman (2012), and obscured differences in habitat associations and life history among these three ciscoes, e.g., Pratt (2012), Pratt and Chong (2012). The discovery of the presumed-extinct *REI* has implications for revising our earlier understanding of the diversity of ciscoes in the LGL and will lead to reinterpretation of findings from previous work to uncover ecological attributes, geographic distribution, and population trends that are now understood to apply to *REI*.

Morphological characters that allow maximum discrimination of ciscoes are likely related to life history and trophic and habitat specializations that contribute to niche segregation and species coexistence as has been found for European whitefish (*C. lavaretus* (*LAV*) complex; Kahilainen et al. 2011, Præbel et al. 2013, Häkli et al. 2018). Our integrated approach of applying genetics to identify morphological differences among ciscoes (“morphology informed by genetics”) facilitated interpretation of morphological differences among ciscoes in terms of adaptation and ecological function. The relationship between genetic and morphological uniqueness of ciscoes was explored with PCA and cluster analysis of morphological data in which fish were labeled by genetic species ID. The high level of discrimination of genetic species by morphometric characters and the agreement between clustering by genetic ID and morphometric characters demonstrates a close relationship between genotype and morphology for the four ciscoes. Relative eye size (ORB), mean length of middle gill rakers (GRML), and gill raker spacing (GRS) were the most informative morphometric characters that best distinguished the four ciscoes. Two other characters, pectoral fin length (PCL) and total gill raker count (GRTC), were highly correlated with the three most informative characters. Together these morphometric characters reflect differences in habitat use and have implications for adaptation and niche segregation. Smaller relative eye size appears associated with the diurnal pelagic life mode of *ART* (Stockwell et al. 2006, Gorman et al. 2012), while the large relative eye size of *HOY* and *KIY* appears associated with inhabiting dark hypolimnetic strata (Yule et al. 2007, Gorman et al. 2012), and associated retinal adaptations in *KIY* allow vision in profundal strata (Harrington et al. 2015). This pattern of segregation by light level and associated eye size is well documented in marine fishes: relative eye size increases with depth (Warrant and Locket 2004) or nocturnal life mode (Schmitz and Wainwright 2011). Similarly, eye size increases in subarctic whitefish dwelling in profundal vs pelagic habitats (Kahilainen and Østbye 2006). Together gill raker number, length, and spacing describe the characteristics of a buccal sieve used to filter food items and reflect trophic specialization that has been well documented in the radiation European whitefishes (Kahilainen and Østbye 2006, Kahilainen et al. 2011, Præbel et al. 2013; Roesch et al. 2013). The numerous, long gill rakers we found in *ART* correlate with specialization on zooplankton (Link and Hoff 1998; Anderson and Smith 1971; Gamble et al. 2011a, b; Rosinski et al. 2020), and the fewer, shorter, more widely spaced gill rakers we found in *KIY* and *REI* correlate with specialization on larger prey such as *Mysis* (Anderson and Smith 1971, Gamble et al. 2011a, Rosinski et al. 2020). *HOY* was distinguished by having a combination of relatively large eyes and numerous medium length gill rakers which may adapt this species to a more generalized diet of copepods, *Mysis*, and *Diporeia* (Anderson and Smith 1971; Gamble et al. 2011a, b; Rosinski et al. 2020). Pectoral fin length further defines differences in morphology that relate to niche segregation by reflecting different life modes and habitat use: *ART*, a pelagic species, preys on pelagic zooplankton and may benefit from reduced drag provided by short pectoral fins; *KIY*, a deep profundal species, undergoes extensive diel vertical migration (Hrabik et al. 2006, Gorman et al. 2012) and have long PCL, which provides increased lift when negatively buoyant (Alexander, 1990; Strand et al. 2005) after descending to deep strata following nighttime transits from shallower strata where they prey on *Mysis* (Hrabik et al. 2006, Gamble et al. 2011a; Rosinski et al. 2020). Other than Lake Superior, the LGL have suffered substantial losses of ecological and morphological diversity of ciscoes. The well-documented decline of the cisco assemblages in lakes Michigan and Huron during the 20^th^ century (Smith 1964, 1968) was accompanied by a loss of morphological diversity of ciscoes and implies a loss of ecological diversification and ecosystem services (Fedorowicz et al. 2024) and is reflected in habitat and diet shifts, changed trophic structure, and a loss of trophic niche breadth among Lake Michigan forage fishes (Bunnell et al. 2012, 2015; Blanke et al. 2018). The most notable losses were species with low gill raker counts, i.e., *C. johannae*, *REI*, and *ZEN*, which were deepwater benthic feeders presumably specializing on *Mysis* and *Diporeia* (Eshenroder et al. 2016). For Lake Superior, the presence of a complement of ciscoes with high gill raker counts (*ART, HOY*) and low gill raker counts (*KIY, REI*) indicates that its cisco complex has retained much of the morphological diversity that has been lost in lakes Michigan and Huron, which now includes a reduced abundance of *ART* and *HOY*. An increased understanding of the functional and ecological basis for morphological differences among extant and extirpated ciscoes can provide insights to their decline and disappearance in some LGL and persistence in others.

### Parallels in Adaptive Radiation of LGL Ciscoes and European Whitefish

The well-studied adaptive radiation of *LAV* in European post-glacial lakes (Schluter 2000, Bernatchez 2004) can serve as a model for understanding the radiation of ciscoes in the LGL. Amundsen et al. (2004a, b), Kahilainen and Lehtonen (2002), Kahilainen et al. (2004), and Kahilainen and Østbye (2006) described three *LAV* ecomorphs in northern Scandinavian lakes that showed niche segregation by habitat and food specialization and associated morphological adaptations. The remarkable concordance in the principal morphological characters that discriminate *ART, HOY, KIY* and *REI* (gill raker length, gill raker spacing, and eye diameter) and the distinct trophic niches of these species closely match those that discriminate *LAV* ecomorphs, suggesting a common pattern of adaptive radiation. Præbel et al. (2013) expanded on the earlier studies of *LAV* in Scandinavian lakes with integrated morphological, isotopic, and genetic analyses and concluded that these lakes were colonized by a common *LAV* ancestor after the retreat of glaciers 10-11k years before present and radiated independently in each lake by ecological specialization into as many as three similar ecomorphs. Häkli et al. (2018) concluded that the adaptive radiation of *LAV* in northern Scandinavian lakes was driven by diversifying selection to exploit benthic and pelagic resources primarily by niche segregation based on gill raker number and body size, and Fang et al. (2022) proposed that ecological specialization and allopatry contributed to reproductive isolation and more rapid evolution of ecologically distinct ecomorphs.

Adaptive radiation of *LAV* in Alpine lakes followed similar albeit more complex patterns of radiation and have yielded as many as six ecomorphs in two lakes and appears to have been facilitated by introgressive hybridization from *LAV* ecomorphs from different lake systems (De-Kayne et al. 2022). Results of genetic analysis of Alpine *LAV* ecomorphs suggests that polygenic architectures and duplicated genes in *LAV* may have contributed to rapid speciation in the absence of geographic isolation (De-Kayne et al. 2022). This conclusion is supported by evidence that past introgression from species now extinct may facilitate rapid evolution in extant forms to create ecomorphs lost through extinction caused by speciation reversal from hybridization (Frei et al. 2022, 2023). The pattern of adaptive radiation in European *LAV* fits West-Eberhard’s (2005) model of sympatric speciation whereby recombination and reorganization of ancestral phenotypes through genetic accommodation can facilitate evolution of novel phenotypes from preexisting phenotypes when stimulated by changes in the environment and the genome. Increased knowledge of the structure of the genome of *LAV* suggests that it harbors an accumulation of genetic diversity through history, hybridization, and genome reorganization and duplication that has the potential to give rise to novel phenotypes when the environment changes and an expansion of environmental resources appear (De-Kayne et al. 2022; Frei et al. 2022, 2023). In essence, *LAV*, as with many biological species, have an innate ability to evolve and can undergo rapid adaptive radiation in relatively short periods, e.g., <10000 years (West-Eberhard 2005; De-Kayne et al. 2022). Especially important in applying this understanding is recognizing that losses in diversity of coregonines from recent anthropogenic disturbances can impede restoration efforts (Frei et al. 2024).

The timing of the radiation of ciscoes in North America following the retreat of the Wisconsin glaciation parallels that of European *LAV* after the retreat of the Weichselian and Würm glaciations. An interesting contrast between the fish communities of North American and European glacial lakes is that whitefish underwent a remarkable radiation in Europe while ciscoes did so in North America. Reasons for this difference are speculative but may be related to profound differences in the size and depth of the principal post-glacial lakes: North America contains massive and deep lakes, e.g., LGL, while in Europe, they are much smaller and generally shallower. Differences in habitat scale and history may reflect evolution of ecomorphotypes along different resource axes: whitefish evolved rapidly along a horizontal littoral-pelagic resource axis in Europe (Schluter 2000; Siwertsson et al. 2010) whereas ciscoes in the LGL evolved along a vertical pelagic to profundal resource axis, e.g., Eshenroder and Burnham-Curtis (1999). In Alpine lakes, the post-glacial radiation of *LAV* into distinct ecomorphs resulted in a diversity of 22 recognized species (De-Kayne et al. 2022) and are being managed as such (Alexander and Seehausen 2021). Recently, Eshenroder et al. (2024) have proposed that the adaptive radiation of ciscoes in the LGL following the Wisconsin glaciation follows a similar evolutionary process as Alpine *LAV*, which would allow parallel evolution of extinct ecomorphs in each LGL that were lost during the Wisconsin glaciation. Alternatively, Backenstose et al. (2024) proposed a pre-Wisconsin glaciation origin for LGL ciscoes, requiring them to survive the Wisconsin glaciation in deepwater refugia. However, Eshenroder et al. (2024) provides evidence that suitable deepwater refugia likely did not exist during the Wisconsin glaciation. Additional genetic and genomic research on the evolution of North American ciscoes following the example of De-Kayne et al. (2022) and Frei et al. (2022, 2023) promises to provide new insights into the adaptive radiation of LGL ciscoes.

### Future directions in restoration and conservation of coregonines

We live in a time of rapid extinction of freshwater fishes and, while funding for research on game and commercial fishes is at a historic high, there is a dearth of research on endangered and non-game fish species (Guy et al. 2021). Especially important is recognizing that losses in diversity of coregonines from anthropogenic disturbances can impede restoration efforts (Frei et al. 2024). The identification of *REI* in Lake Superior represents a reversal of this trend. A recent review of peer-reviewed journal articles and reports that address coregonine biology and management in a broad sense dating back to 1970 showed that there were relatively few articles addressing conservation and restoration, but many articles addressing general topics of fisheries management, e.g., stock assessment, population dynamics, life history, aquaculture, stocking, and genetics (Bunnell et al. 2024). New efforts to reestablish ciscoes lost from LGL Michigan, Huron, Erie, and Ontario require a broad knowledge base of the evolutionary history of ciscoes and the ecology of extant forms, especially those in Lake Superior, which has the most diverse remaining assemblage of ciscoes in the LGL. Inferences about the ecology of ciscoes from their morphology emphasize the importance of building a knowledge base of taxonomy, life history, ecology, and geographic variation of coregonines that is needed to develop effective conservation and restoration strategies for extirpated or declining coregonine populations (Alexander and Seehausen 2021; Bunnell et al. 2023, 2024). In 2018, a multinational program including federal, state, provincial, tribal, and university partners was launched to restore and conserve coregonines in the LGL (Bunnell et al. 2023). That effort can benefit from using the collaborative, long-term studies of coregonine fishes in Alpine lakes in Switzerland as described in Alexander and Seehausen (2021) as a model for developing an integrated program of research and adaptive management to conserve, restore, or rehabilitate coregonines in the LGL.

## Supporting information

Supplement 2 with embedded table and references

Supplement 3 with text, Figure 1, Table 1 and references

Supplement 1 with Figure 1 and Table 1

## ACKNOWLEDGEMENTS

We recognize the following persons for their contributions to this project: Lori Evrard, Yana Grueneis, Jacob Czarnik-Neimeyer, Kara Winter, Katie Tapper, Paul Karpinski, and Sydney Phillips. Matthew Herbert (The Nature Conservancy), Cory Goldsworthy (Minnesota Dept. Natural Resources), U.S. Geological Survey, Ontario Ministry of Natural Resources, and Fisheries and Oceans Canada provided fish samples. Randy Eshenroder (Great Lakes Fishery Commission); and Andrew Honsey, David Bunnell, and Kevin Jones (U.S. Geological Survey) provided thoughtful reviews of the manuscript. This project was funded by the U.S. Environmental Protection Agency’s Great Lakes Restoration Initiative, Habitat and Species Focus Area, Coregonine Template. Any use of trade, firm, or product names is for descriptive purposes only and does not imply endorsement by the U.S. Government. The findings and conclusions in this article are those of the authors and do not necessarily represent the views of the U.S. Geological Survey, U.S. Fish and Wildlife Service, or Fisheries and Oceans Canada.

## Footnotes

1 The Anthropocene currently has no formal status in the Divisions of Geologic Time https://pubs.usgs.gov/fs/2018/3054/fs20183054.pdf). It is used by the authors cited here to informally indicate a time when human activities have significantly affected the global environment.

2 For the sake of clarity, we follow the names conventions for species in the *Common and Scientific Names of Fishes from the United States, Canada and Mexico* (Page et al. 2023). Our use of “species” for ciscoes is not meant to imply that ciscoes are monophyletic biological species *sensu* Mayr (1963).

## Notes

### Competing Interest Statement

The authors have declared no competing interest.

