## Supplement 2 with embedded table and references for "The search for Lake Superior ciscoes as described by W. Koelz in 1929 yields a surprising discovery: Shortnose Cisco (*Coregonus reighardi*), a presumed extinct species"

**Disclaimer:** Any use of trade, firm, or product names in this article is for descriptive purposes only and does not imply endorsement by the U.S. Government.

**Supplement 2.** Description of morphological measures and counts

Morphological measures and counts were assessed manually or from digital images. Type: DI (morphometric digital image measure), M (manual measure), C (categorical), R (ratio), % (percent). All measurements are mm. Source: K (Koelz 1929), E (Eshenroder et al. 2016), G (Gorman et al., this paper). 38 measures and counts used in morphometric analyses are indicated with a "‡". Measures and counts used in identification of fish with the Cisco Score Card are indicated with a "*". All measurements are in mm.

| # | Code | Description | Details | Type | Source |
|---|---|---|---|---|---|
| 1 | ADB ‡ | Adipose fin base | length of the base of the adipose fin from point of insertion to the distal end of the base | DI | K, G |
| 2 | AD-CA ‡ | Adipose-caudal fin distance | distance from the insertion of the adipose fin to the insertion of the first of the upper procurrent caudal rays (first of the extended caudal rays) | DI | K, G |
| 3 | ADL ‡ | Adipose fin length | length of the adipose fin from point of insertion to the distal end of fleshy lobe of the fin | DI | K, G |
| 4 | ANB ‡ | Anal fin base length | length of the base of the anal fin (anal fin base) | DI | K, G |
| 5 | AN-CA_M ‡ | Anal to caudal fin midline distance | distance from the insertion of the anal fin to the caudal peduncle visualized along the body midline with aid of vertical lines indicating location of character landmarks | DI | G |
| 6 | ANH ‡ | Anal fin height | height of the anal fin (anal fin height) | DI | K, G |

| | | | | | |
|---|---|---|---|---|---|
| 7 | BDD ‡ | Body depth at dorsal fin | body depth measured at the insertion of the dorsal fin | DI | E, G |
| 8 | BDM ‡ | Body depth, maximum | the greatest vertical depth of the body (this measure may be compromised by the state of bloating of the visceral body of the fish) | M, DI | K, G |
| 9 | BDM_ratio | Ratio of STL/BDM | calculated as STL/BDM | M (R) | G |
| 10 | DO-AD ‡ | Dorsal to adipose fin distance | distance from the anterior insertion of the dorsal fin to the anterior insertion of the adipose fin (dorsal to adipose) | DI | K, G |
| 11 | DO-AN_M ‡ | Dorsal to anal fin midline distance | distance from the insertion of the dorsal fin to the insertion of the anal fin visualized along the body midline with aid of vertical lines indicating location of character landmarks | DI | G |
| 12 | DOB ‡ | Dorsal fin base length | length of the base of the dorsal fin (dorsal fin base) | DI | K, G |
| 13 | DO-CA ‡ | Dorsal to caudal fin distance | distance from the insertion of the dorsal fin to the insertion of the first of the upper procurrent caudal rays (first of the extended caudal rays) | DI | G |
| 14 | DO-CA_M ‡ | Dorsal to caudal fin midline distance | distance from the insertion of the dorsal fin to the caudal peduncle visualized along the body midline with aid of vertical lines indicating location of character landmarks | DI | G |
| 15 | DOH ‡ | Dorsal fin height | height of the dorsal fin (dorsal fin height) | DI | K, E, G |
| 16 | EYE | Diameter of the eyeball | longitudinal diameter of the eyeball. Very close to the measure of the ORB | M | K |
| 17 | GRAL ‡ | Gill raker arch length | total length of gill raker arch measured along a line from the insertion of gill rakers at the distal end of the upper (shorter) arm to the central inflection, to the end of the insertion of gill rakers on the lower (longer) arm including arch with nubs | DI | G |

| | | | | | |
|---|---|---|---|---|---|
| 18 | GRL * | Gill raker length | Length of the middle gill rakers measured in situ with calipers (not individual lengths) | M | G |
| 19 | GRL_cat * | Gill raker length, category | Average length of middle gill rakers. 1: short (< 7 mm), 2: med-short (7-8 mm). 3: medium (9-11), 4: med-long (12-14 mm), 5: long (> 14 mm) | M (C) | G |
| 20 | GRLA ‡ | Gill raker length, average | Calculated as average length of the 3 closest rakers to the inflection point on the lower arm of the gill raker arch | DI | G |
| 21 | GRLM ‡ | Gill raker length, maximum | Length of the longest of the three middle gill rakers | DI | G |
| 22 | GRS ‡ | Gill raker spacing | Center-to-center distance of gill rakers calculated as GRAL/GRTC. The inverse of GRS is GRD, gill raker density (#/mm) | DI | G |
| 23 | GRTC (TGR) ‡ | Gill raker total count | total number of gill rakers; nubs are not included (coded as TGR in Eshenroder et al. 2016) | DI | K, E, G |
| 24 | GRTC_cat* | Gill raker total count, category | 1: low (<35), 2: med-low (35-37), 3: med. (38-40), 4: med-high (41-44), 5: high (>44) | M (C) | G |
| 25 | HDD ‡ | Head depth | distance from the occiput to the outer edge of the boundary between the suboperculum and the interoperculum | DI | K, G |
| 26 | HDL (HLL) ‡* | Head length | distance from the junction of the premaxillaries (tip of snout) to the extreme bony margin of the operculum, not including the operculum membrane, as measured parallel to the longitudinal axis of the body (coded as HLL in Eshenroder et al. 2016) | M, DI | K, E, G |
| 27 | HDL_ratio | Ratio of STL/HDL | calculated as STL/HDL | M (R) | K, E, G |
| 28 | MDL (MDB) ‡ | Mandible length | distance from the anterior symphysis of the lower jaw to the posterior edge of the mandibular-cranial joint (coded as MDB in Eshenroder et al. 2016) | M, DI | K, E, G |

| | | | | | |
|---|---|---|---|---|---|
| 29 | MDL_cat * | Mandible length category relative to tip of snout | 1: subterminal, 2: even-included, 3: extended | M (C) | G |
| 30 | MXL ‡ | Maxillary length | distance from the symphysis of the premaxillaries to the caudal end of the maxillary bone (upper jaw) | M, DI | K, E, G |
| 31 | OC-DO_M ‡ | Occiput to dorsal fin midline distance | distance from the occiput to the insertion of dorsal fin visualized along the body midline with aid of vertical lines indicating location of character landmarks | DI | G |
| 32 | ODD (OID) ‡* | Occiput-dorsal distance | distance from the occiput located at the dorsal margin of the skull to the insertion of first dorsal fin ray (coded as OID on Cisco Score Card) | M, DI | G |
| 33 | OPD (OPT) ‡* | Occiput-pectoral distance | lateral distance visualized from a vertical line marking the occiput to the tip of the pectoral fin aligned parallel to the body axis (coded as OPT on Cisco Score Card) | M, DI | G |
| 34 | ORB (OOL) ‡* | Orbital length (eye opening diameter) | distance between the anterior fleshy margin and the posterior bony margin of the orbit (calipers aligned against the margins of the orbital rim (Vuorinen et al. 1993) (Coded as OLL in Eshenroder et al. 2016) | M, DI | E, G |
| 35 | ORB_cat * | Orbit size category based on HDL/ORB | categories of HDL/ORB, 1: small (>4.4), 2: medium (>4.2-4.4), 3: med-large (>4.0-4.2), 4: large (<4.0) | M (C) | G |
| 36 | ORB_ratio * | Ratio HDL/ORB | calculated as HDL/ORB | M (R) | K, E, G |
| 37 | PAD‡ | Pelvic-anal fin distance | distance from insertion of the pelvic fin to the insertion of anal fin | DI | K, E, G |
| 38 | PCL ‡* | Pectoral fin length | length of pectoral fin | M, DI | K, E, G |
| 39 | PCL% * | % pectoral fin length calculated from OPD/ODD | calculated as OPD/ODD (alternate to PCL/PDD, #40) | M (%) | G |

| | | | | | |
|---|---|---|---|---|---|
| 40 | PCL/PDD% | % pectoral fin length calculated from PCL/PDD | calculated as PCL/PDD (alternate to PCL%, OPD/ODD, #39) | M (%) | G |
| 41 | PCL_cat * | Pectoral fin length category based on OPD/ODD | 1: short (≤60%), 2: med-short (61-65%), 3: med. (66-70%), 4: long (>70%) | M (C) | G |
| 42 | PDD (PCD) | Pectoral-dorsal distance | lateral distance from Insertion of pectoral fin to insertion of dorsal fin. Can be calculated as ODD-OPD+PCL. (coded as PCD on Cisco Score Card) | M, DI | G |
| 43 | PMA ‡ | Premaxillary angle | angle between the anterior margin of the premaxillae and the horizontal axis of the head, in degrees | DI | E, G |
| 44 | PMA_cat * | Premaxillary angle, category | 1: straight, 2: low, 3: medium, 4: high | M (C) | G |
| 45 | POL ‡ | Snout length; preorbital length (snout length) | distance from the tip of snout to anterior fleshy margin of the orbital rim with calipers anchored against the margins of the rim (Pre-Orbital Length; Vuorinen et al. 1993) | M, DI | K, E, G |
| 46 | PPD ‡ | Pectoral-pelvic distance | distance from the insertion of pectoral fin to the insertion of pelvic fin | DI | K, E, G |
| 47 | PVD ‡* | Pelvic-vent distance | distance from insertion of the pelvic fin to the center of the vent | M, DI | G |
| 48 | PVL ‡* | Pelvic fin length | length of the pelvic fin | M, DI | K, E, G |
| 49 | PVL% * | % pelvic fin length | calculated as PVL/PVD | M (%) | G |
| 50 | PVL/PAD% | % pelvic fin length relative to PAD | calculated as PVL/PAD, alternative to PVL% (PVL/PVD, #49) | DI (%) | G |
| 51 | PVL_cat * | Pelvic fin length category based on PVL/PVD | 1: ≤60%, 2: 65-75%, 3: 75-85%, 4: ≥90% | M (C) | G |

| | | | | | |
|---|---|---|---|---|---|
| 52 | SN-AN ‡ | Snout to anal fin distance | distance from the junction of the premaxillaries to the base of the first anal ray (snout to anal) | DI | K, G |
| 53 | SN-DO ‡ | Snout to dorsal fin distance | distance from the junction of the premaxillaries to the base of the first dorsal ray (snout to dorsal) | DI | K, G |
| 54 | SN-DO_M ‡ | Snout to dorsal fin midline distance | distance from tip of snout to the insertion of the dorsal fin visualized along the body midline with aid of vertical lines indicating location of character landmarks | DI | G |
| 55 | SN-OC ‡ | Premaxillary-occiput distance | distance from the junction of the premaxillaries to the end of the supraoccipital bone, not to the beginning of the scales (occiput) | DI | K, G |
| 56 | SN-OC_M ‡ | Snout to occiput midline distance | distance from tip of snout to the occiput visualized along the body midline with aid of vertical lines indicating location of character landmarks | DI | G |
| 57 | STL* | Standard length | Tip of snout to the bony terminus of the caudal peduncle | M, DI | K, E, G |
