## Supplement 3 with text, Figure 1, Table 1 and references for "The search for Lake Superior ciscoes as described by W. Koelz in 1929 yields a surprising discovery: Shortnose Cisco (*Coregonus reighardi*), a presumed extinct species"

**Disclaimer:** Any use of trade, firm, or product names in this article is for descriptive purposes only and does not imply endorsement by the U.S. Government.

Bureau approved

**Supplement 3.** Identification of ciscoes using a scoring matrix, the "Cisco Score Card"

Dichotomous keys do not yield reliable identification of ciscoes due to overlapping morphological variation among species and changing character states with ontogeny. Alternatively, species can be distinguished based on combinations of characters, but without a structured approach, reliable identifications are uncertain. To address these impediments to reliable identification of ciscoes, we developed a tabular scoring matrix (Cisco Score Card, CSC) to guide and facilitate reliable identification of Lake Superior ciscoes *Coregonus artedi* (*ART*), *C. hoyi* (*HOY*), *C. kiyi* (*KIY*), *C. nigripinnis* (*NIG*), *C. reighardi* (*REI*), *C. zenithicus* (*ZEN*). The CSC was designed as a self-guide to identifying ciscoes in a field or laboratory setting and does not require specialized training nor specialized tools other than an inexpensive set of calipers. Importantly, the CSC provides a written record of identification for review and redaction (Suppl. 3, Fig. 1). The CSC was designed for identification of adults, the life stage when morphological character states are fully expressed, and the life stage on which species descriptions in Koelz (1929) and Eshenroder et al. (2016) are based. The CSC can be used with younger life stages, but with the caveat that less reliable identifications will result. We recognized that accurate identification is not possible for every fish, but most fish can be identified with a relative measure of confidence using a combination of character states that are definitive for each species.

The CSC uses a combination of seven primary and five secondary morphological characters (Suppl. 3, Table 1) that are scored according to the level of match with the definitive character states for each of the six cisco species. Most of these characters are based on those used by Koelz (1929) and Eshenroder et al. (2016) to distinguish cisco species. Definitive character states represent those that distinguish the six species in descriptions provided in Koelz (1929) and Eshenroder et al. (2016). A matching morphological character state, i.e., falling within the range of definitive character states for one of the six species, was assigned a character score of "1", and a mismatching character state, i.e., not within range of definitive character states for a species, was assigned a "0". When a character state was judged as having a marginal or partial match, i.e., intermediate between two species, a character score of "0.5" was assigned. Character scores were assigned for each morphological character for each of the six species. Scores for primary morphological characters were double-weighted, and scores for secondary morphological characters were single-weighted. Character scores were summed for each species to yield a composite species score (range, 0 - 19) and the species with the highest composite score represented the consensus species. Ties and near-ties occurred frequently due to morphological variation and conflicting combinations of character states. A conformation score, ranging from 5 to 10, was generated for the consensus species that reflected how well the fish's morphological character states matched the definitive character states for that species. A fish with a conformation score of 5 reflects an equivocal identity: the combination of character states does not match any species. A score of 10 reflects an unequivocal identity: a perfect match with the definitive character states for one of the six species. Conformation scores reflect the level of confidence in the species identification: conformation scores <6.0 represent no confidence, scores 6.0 to <7.0 represent low confidence, and scores of 7.0 to < 8.0 represent moderate confidence and scores ≥ 8 to 10.0 represent high confidence. Fish with conformation scores <6.0 were recorded as unidentified, fish with scores ≥6.0 to <7.0 were assigned a provisional identification, and fish with scores ≥7.0 were assigned a putative identification. Fish with ties in species composite scores or near-ties (difference ≤3.0) received a conformation score of ≤6.0. In cases where a composite score is ≥10 for the consensus species but the fish presented a low character score in a defining primary character, that fish received a lower conformation score. An example would be a fish that received the highest composite score for *KIY* (score ≥10) and second highest score for *HOY* (score within 3.0) and the gill raker length was too long for *KIY*. After a review of measured characters by the two persons, this fish might be given a provisional identification as a *KIY* with a lower conformation score, e.g., <7.0. We emphasize that the CSC method does not yield absolute identifications, but conformation scores reflect the level of confidence in the identification.

The identification of fish was conducted independently by two persons well-trained in identifying ciscoes. The first person completes manual morphological measurements, evaluates the morphological character states, assigns character scores for each morphological character for each of the six cisco species on the CSC, generates composite scores for each species, selects the consensus species, generates a conformation score for the consensus species, and proposes a species identification. The second person independently reviews the measurements, character scores, composite scores, selection of consensus species and conformation score, species identification, and makes corrections as needed. Then the two persons review the results together and come to agreement on a finalized consensus species, conformation score, and species identification. This Delphi approach addresses the potential subjectivity in identifying ciscoes using combinations of variable characters and provides a permanent record of the identification process that can be reviewed and redacted later, such as when new information indicates that the weighting of character scores warrants revision. For example, the findings of our research indicate that relative eye size, gill raker length and gill raker spacing are the most informative morphological characters for discriminating the Lake Superior ciscoes and should be given greater weight in assigning scores and deciding a species ID and conformation score.

The CSC method addresses our goal to create a more quantifiable method of identification of ciscoes that is subject to review and revision. Toward that end, each CSC record was 1) scanned and stored digitally, 2) entered into an electronic database, 3) accompanied by digital photographs

Bureau approved

of the fish and gill rakers, 4) linked to a database of morphometric data derived from digital images, 5) linked to databases of age, diet, and genetic analyses.

| LAKE SUPERIOR CISCO SCORE CARD v8c | | Spec. # | | | | Species | | | Identifiers | | | Date |
|---|---|---|---|---|---|---|---|---|---|---|---|---|
| *Character* | √ | **Artedi (202)** | √ | **Nigripinnis (207)** | √ | **Kiyi (206)** | √ | **Hoyi (204)** | √ | **Zenithicus (210)** | √ | **Reighardi (208)** |
| ***Mandible length** | | ~even-include | | ~even-include | | extended | | extended | | subterm | | subterm |
| ***Premax angle** | | low or none-F | | low -D | | ~straight -C | | ~straight -B | | medium -A | | high -E |
| ***Eye size** | | small-med | | med-large | | large | | med-large | | small-med | | small-med |
| ***gill raker length** | | med-long,long | | med-long | | shrt,med-short | | med | | med-short,med | | short |
| ***gill raker count** | | med-high | | med | | low, med-low | | med | | med-low | | low |
| ***Pectoral fin length** | | short ≤60% | | med 65-70% | | long 70-85% | | med 65-70% | | med 65-70% | | med-shrt 60-65% |
| ***Pelvic fin length** | | short ≤60%-A | | med 65-75%-B | | long >90%- C | | m-lg 75-85%-D | | med 65-75%-B | | med 65-75%-B |
| ***body shape** | | elongate -A | | ovate -C | | sub-ovate -D | | sub-ovate -D | | sub-terete -B | | sub-terete -B |
| ***body wall** | | thick | | med-thick | | thin | | thin | | medium | | medium |
| *pectoral pigment* | | low | | dark | | med-dark | | med | | med | | low |
| *pelvic pigment* | | none to low | | low | | none to low | | none to low | | none to low | | none to low |
| *pigment dorsal* | | med-grn-blu | | dark-gry | | dark-blu-gry | | med-amb | | med-amb-other | | low-med-amb |
| **Scores (# checked)** | | | | | | | | | | | | |
| Consensus sp. | | Conform. Score | | | | Other pigmnt/color | | | | Up/low jaw pgmnt | | |
| TL STL | Rel. age (GR condition) | | | | | Color,rel. size: caud,dors,anl,adip | | | | | | |
| Comments | | | | | | | | | | Eye size ratio: < 4.0 -large; > 4.0-4.2 -med-large; > 4.2-4.4 -med; > 4.4 -small | | |

ocpt-ins dor_____ ocpt-pec tip_____ / _____%, pec ln _____ ins plv-vnt_____ plv ln_____ /_____% orb_____hd ln_____/ratio_____ ,eye_____

Notes ______________________________________________________________

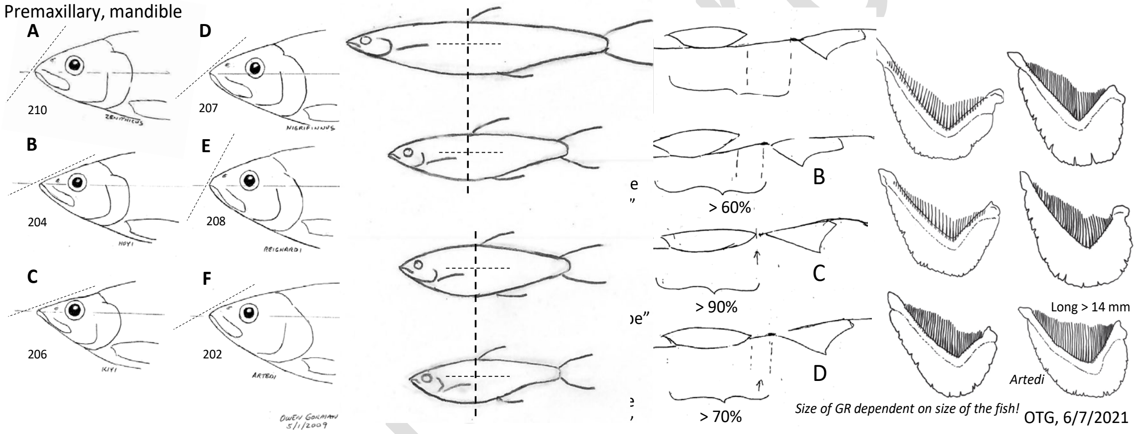


**Supplement 3, Figure 1.** Cisco Score Card (CSC) for Lake Superior Ciscoes used for identification of ciscoes. A matrix of six species and 12 morphological characters is shown. The first seven characters are primary diagnostic characters largely responsible for identification of ciscoes and are double-weighted. The last five characters are secondary characters and less critical and often dependent on the condition of the fish, e.g., partial decomposition, faded pigment and coloration. Description of morphological characters recorded on the CSC are listed in Suppl. 3, Table 1.

**Supplement 3, Table 1.** Description of morphological characters and measures used in the Cisco Score Card. For primary characters eye size, gill raker length, pectoral fin length, pelvic fin length, scores are based on measured characters. For *Characters based on measures and counts*, numbers following "#" refer to morphological characters listed in Supplement 2, *Morphological measures and counts*. Characters derived from Koelz (1929) are denoted with a "*".

| ***Primary characters*** | **unit** | **description** |
|---|---|---|
| Mandible length* | category | Estimated from diagrams: even (F), ~even-included (D), extended (B, C), subterminal (A, E) |
| Premax angle* | category | Estimated from diagrams: low or none (F), low (D), ~straight (C), ~straight (B), medium (A), high (E) |
| Eye size* | ratio | Calculated from ORB_ratio: small (>4.4), medium (4.2-4.4), medium-large (4.0-4.2); large (<4.0) |
| Gill raker length | category | Categorized from GRL: short (< 7 mm), medium short (7-8 mm), medium (9-11 mm), medium long (12-14 mm), long (>14 mm) |
| Gill raker count* | category | Estimated from diagrams: low (*REI*, <35), medium-low (*KIY*, 35-37), medium (*HOY, ZEN*, 38-40), medium-high (*ART, NIG*, 41-44), high (*ART*, >44) |
| Pectoral fin length* | category | Calculated from PCL%: short (≤ 60%), medium-short (60-65%), medium (65-70%), long (>70%) |
| Pelvic fin length* | category | Calculated from PVL%: short (≤ 60%), medium (65-75%), medium-long (75-85%), long (>90%) |
| ***Secondary characters*** | | |
| Body shape | category | Estimated from diagrams: A, elongate (sub-terete); B, sub-terete (D-shaped); C, ovate (egg-shaped); D, sub-ovate (P-shaped) |
| Body wall | category | Estimated from example photos: thin, medium, medium thick, thick |
| Pectoral fin pigment | category | Estimated from example photos: low, medium, medium-dark, dark |
| Pelvic fin pigment | category | Estimated from example photos: none, low |
| Dorsal body pigment, color | category | Estimated from example photos. pigment: low, medium, dark; color: green, blue, gray, blue-gray, amber, other |
| **Characters based on measures and counts** | | |
| GRL, #18 | mm | length of the middle gill rakers measured in situ with calipers; reflects average length of the three middle gill rakers |
| GRL_cat, #19 | category | average length of middle gill rakers. 1: short (< 7 mm), 2: med-short (7-8 mm). 3: medium (9-11), 4: med-long (12-14 mm), 5: long (> 14 mm) |
| GRTC_cat, #24* | category | gill raker total count. 1: low (REI, <35), 2: med-low (KIY, 35-37), 3: med. (HOY, ZEN, 38-40), 4: med-high (ART, NIG, 41-44), 5: high (ART, >44) |
| HDL (HLL), #26* | mm | head length: tip of snout to bony margin of operculum parallel to the body midline |

Bureau approved

| | | |
|---|---|---|
| ODD (OID), #32 | mm | occiput-dorsal distance: distance from occiput to insertion of first dorsal fin ray |
| OPD (OPT), #33 | mm | occiput-pectoral distance: lateral distance visualized from a vertical line marking the occiput to the tip of the pectoral fin aligned parallel to the body axis |
| ORB (OOL), #34* | mm | orbital length (eye opening diameter) |
| ORB_cat, #35 | category | categories of HDL/ORB: 1: small (>4.4), 2: medium (>4.2-4.4), 3: med-large (>4.0-4.2), 4: large (<4.0) |
| ORB_ratio, #36* | ratio | ratio of head length/orbital length (HDL/ORB) |
| PCL _cat, #41 | category | pectoral fin length category relative to OPD/ODD: 1: short (≤60%), 2: med-short (61-65%), 3: med. (66-70%), 4: long (>70%) |
| PCL%, #39 | % | % pectoral fin length, calculated as OPD/ODD |
| PCL, #38* | mm | pectoral fin length |
| PCL/PDD%, #40 | % | pectoral length/pectoral-dorsal distance: PCL/PDD (alternate to PCL%) |
| PDD (PCD), #42 | mm | pectoral-dorsal distance: lateral distance from Insertion of pectoral fin to insertion of dorsal fin, calculated as ODD-OPD+PCL |
| PVD, #47 | mm | pelvic-vent distance: insertion of pelvic fin to vent distance |
| PVL%, #49 | % | % pelvic fin length, calculated as PVL/PVD |
| PVL, #48* | mm | pelvic fin length |
| PVL_cat, #50 | category | pelvic fin length category relative to PVL/PVD: 1: ≤60%, 2: 65-75%, 3: 75-85%, 4: ≥90% |

Bureau approved
