## Supplement 1 with Figure 1 and Table 1 for "The search for Lake Superior ciscoes as described by W. Koelz in 1929 yields a surprising discovery: Shortnose Cisco (*Coregonus reighardi*), a presumed extinct species"

**Disclaimer:** Any use of trade, firm, or product names in this article is for descriptive purposes only and does not imply endorsement by the U.S. Government.

**Supplement 1.** Results of STRUCTURE Bayesian cluster analysis of Principal Components Analysis (PCA) scores of species-associated allele frequency differences

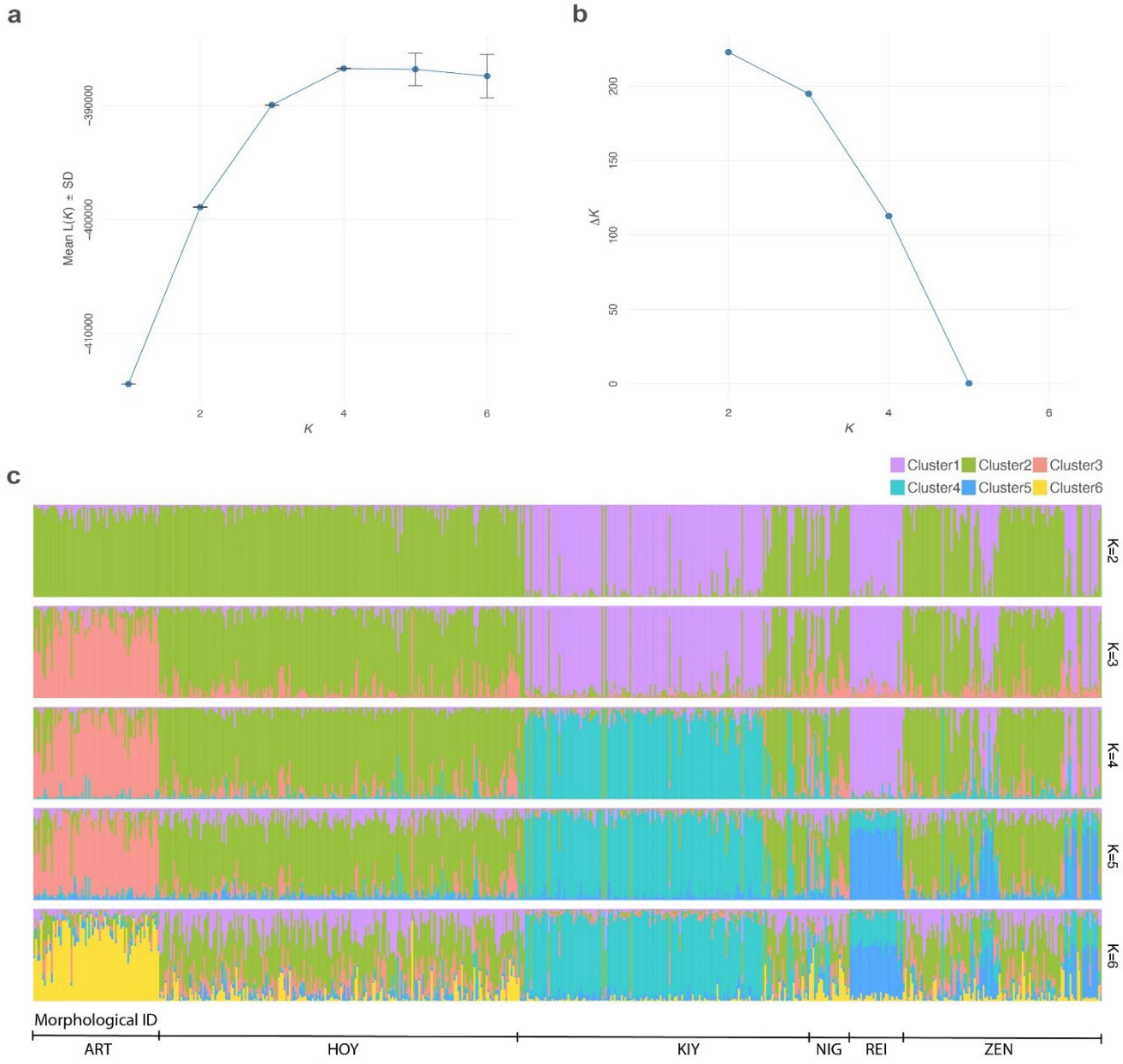


**Supplement 1, Figure 1.** Mean maximum likelihood (a) and Evanno (b) plots indicated support for *K*=4 and *K*=2 clusters, respectively, and STRUCTURE plots for *K*=2-6 (c).

**Supplement 1, Table 1.** STRUCTURE Q-scores for the 602 genotyped ciscoes for clusters K = 2 - 6.

| MEL_ID | K=2 | | K=3 | | | K=4 | |
|---|---|---|---|---|---|---|---|
| | X2.Cluster1 | X2.Cluster2 | X3.Cluster1 | X3.Cluster2 | X3.Cluster3 | X4.Cluster1 | X4.Cluster2 |
| ART7919 | 0.0108 | 0.9892 | 0.0134 | 0.4796 | 0.507 | 0.0134 | 0.4774 |
| ART7923 | 0.0142 | 0.9858 | 0.0132 | 0.2154 | 0.7712 | 0.019 | 0.2066 |
| ART7924 | 0.038 | 0.962 | 0.0386 | 0.468 | 0.4934 | 0.0172 | 0.4694 |
| ART7925 | 0.0412 | 0.9588 | 0.0424 | 0.464 | 0.4936 | 0.061 | 0.4474 |
| ART7928 | 0.0332 | 0.9668 | 0.0192 | 0.0272 | 0.9532 | 0.0112 | 0.0174 |
| ART7929 | 0.049 | 0.951 | 0.0578 | 0.6684 | 0.2736 | 0.0784 | 0.6538 |
| ART7930 | 0.1052 | 0.8948 | 0.1224 | 0.67 | 0.2074 | 0.0174 | 0.6684 |
| ART7933 | 0.0498 | 0.9502 | 0.0528 | 0.382 | 0.5654 | 0.016 | 0.3754 |
| ART7945 | 0.038 | 0.962 | 0.0242 | 0.1786 | 0.7974 | 0.0148 | 0.1698 |
| ART7946 | 0.0082 | 0.9918 | 0.0096 | 0.3738 | 0.6166 | 0.0084 | 0.3714 |
| ART8565 | 0.0158 | 0.9842 | 0.021 | 0.6778 | 0.3012 | 0.017 | 0.6766 |
| ART8574 | 0.0148 | 0.9852 | 0.0104 | 0.007 | 0.9822 | 0.0078 | 0.0048 |
| ART8575 | 0.0112 | 0.9888 | 0.01 | 0.2224 | 0.7674 | 0.01 | 0.2154 |
| ART8576 | 0.0474 | 0.9526 | 0.0354 | 0.0134 | 0.9512 | 0.0172 | 0.008 |
| ART8578 | 0.106 | 0.894 | 0.065 | 0.114 | 0.8212 | 0.0816 | 0.0906 |
| ART8579 | 0.1882 | 0.8118 | 0.164 | 0.1652 | 0.6704 | 0.0074 | 0.1424 |
| ART8580 | 0.1212 | 0.8788 | 0.0496 | 0.0132 | 0.9374 | 0.029 | 0.008 |
| ART8581 | 0.0514 | 0.9486 | 0.031 | 0.0184 | 0.9504 | 0.0102 | 0.01 |
| ART8582 | 0.023 | 0.977 | 0.0096 | 0.0254 | 0.9652 | 0.0112 | 0.0182 |
| ART8583 | 0.0724 | 0.9276 | 0.0326 | 0.008 | 0.9594 | 0.0086 | 0.0058 |
| ART8590 | 0.1628 | 0.8372 | 0.065 | 0.0684 | 0.8666 | 0.0688 | 0.0622 |
| ART8591 | 0.0322 | 0.9678 | 0.0486 | 0.7376 | 0.2138 | 0.0438 | 0.7312 |
| ART8593 | 0.1248 | 0.8752 | 0.0586 | 0.162 | 0.7788 | 0.0534 | 0.1514 |
| ART8594 | 0.2708 | 0.7292 | 0.238 | 0.0626 | 0.6994 | 0.2346 | 0.0582 |
| ART8595 | 0.0908 | 0.9092 | 0.0564 | 0.2132 | 0.7306 | 0.0644 | 0.2018 |
| ART8596 | 0.0458 | 0.9542 | 0.0292 | 0.2764 | 0.6946 | 0.0142 | 0.2684 |
| ART8607 | 0.028 | 0.972 | 0.0084 | 0.0054 | 0.9864 | 0.0038 | 0.003 |
| ART8608 | 0.1508 | 0.8492 | 0.1076 | 0.0382 | 0.854 | 0.0422 | 0.0322 |
| ART8609 | 0.011 | 0.989 | 0.0072 | 0.066 | 0.9268 | 0.0092 | 0.0534 |
| ART8610 | 0.3072 | 0.6928 | 0.2506 | 0.1014 | 0.6482 | 0.0116 | 0.0642 |
| ART8611 | 0.0432 | 0.9568 | 0.016 | 0.0092 | 0.9746 | 0.014 | 0.0058 |
| ART8612 | 0.2704 | 0.7296 | 0.2276 | 0.0338 | 0.7386 | 0.0164 | 0.0228 |
| ART8613 | 0.0152 | 0.9848 | 0.0074 | 0.0036 | 0.9888 | 0.0034 | 0.0022 |
| ART8615 | 0.0356 | 0.9644 | 0.0182 | 0.1868 | 0.7952 | 0.015 | 0.1812 |
| ART8616 | 0.04 | 0.96 | 0.0328 | 0.0342 | 0.933 | 0.0048 | 0.0248 |
| ART8617 | 0.0728 | 0.9272 | 0.0284 | 0.0054 | 0.9662 | 0.0284 | 0.003 |
| ART8618 | 0.1852 | 0.8148 | 0.0916 | 0.013 | 0.8954 | 0.0894 | 0.0082 |
| ART8622 | 0.0266 | 0.9734 | 0.016 | 0.2314 | 0.7526 | 0.022 | 0.2248 |
| ART8624 | 0.0342 | 0.9658 | 0.0076 | 0.0858 | 0.9066 | 0.005 | 0.0762 |
| ART8625 | 0.1214 | 0.8786 | 0.0294 | 0.0568 | 0.9142 | 0.0094 | 0.044 |
| ART8626 | 0.0338 | 0.9662 | 0.016 | 0.006 | 0.978 | 0.0054 | 0.0034 |

| | | | | | | | |
|---|---|---|---|---|---|---|---|
| ART8628 | 0.2564 | 0.7436 | 0.098 | 0.0038 | 0.898 | 0.0696 | 0.0026 |
| ART8629 | 0.0134 | 0.9866 | 0.0098 | 0.1184 | 0.8718 | 0.0068 | 0.1102 |
| ART8630 | 0.0456 | 0.9544 | 0.0294 | 0.197 | 0.774 | 0.0058 | 0.177 |
| ART8631 | 0.024 | 0.976 | 0.0076 | 0.2472 | 0.7452 | 0.0054 | 0.247 |
| ART8632 | 0.1298 | 0.8702 | 0.0104 | 0.0048 | 0.9848 | 0.0062 | 0.003 |
| ART8633 | 0.0772 | 0.9228 | 0.0252 | 0.2938 | 0.6806 | 0.0298 | 0.2862 |
| ART8634 | 0.1924 | 0.8076 | 0.076 | 0.0108 | 0.9132 | 0.0308 | 0.0068 |
| ART8635 | 0.0076 | 0.9924 | 0.0046 | 0.0064 | 0.9886 | 0.0028 | 0.004 |
| ART8636 | 0.0738 | 0.9262 | 0.0616 | 0.0678 | 0.871 | 0.0144 | 0.0516 |
| ART8637 | 0.1158 | 0.8842 | 0.0898 | 0.0342 | 0.876 | 0.0042 | 0.015 |
| ART8638 | 0.0084 | 0.9916 | 0.0078 | 0.4272 | 0.565 | 0.0134 | 0.4252 |
| ART8639 | 0.181 | 0.819 | 0.055 | 0.0696 | 0.8754 | 0.0188 | 0.054 |
| ART8640 | 0.0732 | 0.9268 | 0.068 | 0.4704 | 0.4618 | 0.0914 | 0.4532 |
| ART8641 | 0.1602 | 0.8398 | 0.1286 | 0.3072 | 0.564 | 0.0166 | 0.2992 |
| ART8642 | 0.12 | 0.88 | 0.085 | 0.2244 | 0.6904 | 0.1426 | 0.1932 |
| ART8643 | 0.034 | 0.966 | 0.0098 | 0.0144 | 0.976 | 0.0084 | 0.01 |
| ART8644 | 0.0134 | 0.9866 | 0.014 | 0.337 | 0.6492 | 0.013 | 0.3348 |
| ART8645 | 0.1014 | 0.8986 | 0.051 | 0.088 | 0.861 | 0.079 | 0.0558 |
| ART8647 | 0.0314 | 0.9686 | 0.018 | 0.1404 | 0.8416 | 0.0184 | 0.1356 |
| ART8648 | 0.0294 | 0.9706 | 0.0154 | 0.3 | 0.6846 | 0.0106 | 0.2956 |
| ART8649 | 0.0206 | 0.9794 | 0.0196 | 0.0962 | 0.8844 | 0.0348 | 0.0776 |
| ART8650 | 0.1948 | 0.8052 | 0.0684 | 0.1738 | 0.758 | 0.0322 | 0.1706 |
| ART8651 | 0.0886 | 0.9114 | 0.0088 | 0.0068 | 0.9846 | 0.0064 | 0.0048 |
| ART8652 | 0.102 | 0.898 | 0.0482 | 0.051 | 0.9008 | 0.0112 | 0.0328 |
| ART8653 | 0.0062 | 0.9938 | 0.0058 | 0.293 | 0.7014 | 0.0056 | 0.2902 |
| ART8654 | 0.0046 | 0.9954 | 0.006 | 0.4004 | 0.5938 | 0.0038 | 0.397 |
| ART8655 | 0.0912 | 0.9088 | 0.0142 | 0.014 | 0.972 | 0.0072 | 0.008 |
| COR8439 | 0.0216 | 0.9784 | 0.012 | 0.0926 | 0.8954 | 0.0204 | 0.0854 |
| COR8440 | 0.216 | 0.784 | 0.1556 | 0.0146 | 0.8298 | 0.039 | 0.0108 |
| COR8441 | 0.11 | 0.89 | 0.0822 | 0.317 | 0.6012 | 0.012 | 0.311 |
| HOY6647 | 0.0054 | 0.9946 | 0.009 | 0.9364 | 0.0546 | 0.0106 | 0.945 |
| HOY6649 | 0.0108 | 0.9892 | 0.0204 | 0.8428 | 0.137 | 0.0114 | 0.845 |
| HOY6650 | 0.0112 | 0.9888 | 0.0196 | 0.9332 | 0.0472 | 0.0278 | 0.9308 |
| HOY6651 | 0.0048 | 0.9952 | 0.0066 | 0.7612 | 0.2318 | 0.0182 | 0.7606 |
| HOY6653 | 0.0222 | 0.9778 | 0.0322 | 0.9614 | 0.0068 | 0.0086 | 0.9566 |
| HOY6654 | 0.0234 | 0.9766 | 0.033 | 0.9614 | 0.0056 | 0.0302 | 0.9508 |
| HOY6660 | 0.0114 | 0.9886 | 0.0186 | 0.9614 | 0.0198 | 0.008 | 0.9648 |
| HOY6661 | 0.0064 | 0.9936 | 0.0182 | 0.9468 | 0.035 | 0.0254 | 0.944 |
| HOY6664 | 0.0972 | 0.9028 | 0.094 | 0.5538 | 0.3524 | 0.0158 | 0.5498 |
| HOY6665 | 0.0054 | 0.9946 | 0.009 | 0.7656 | 0.2252 | 0.0044 | 0.7648 |
| HOY6666 | 0.0052 | 0.9948 | 0.008 | 0.6998 | 0.2922 | 0.0098 | 0.6996 |
| HOY6668 | 0.0134 | 0.9866 | 0.0146 | 0.983 | 0.0024 | 0.01 | 0.9796 |
| HOY6669 | 0.0382 | 0.9618 | 0.0692 | 0.8324 | 0.0986 | 0.0108 | 0.835 |
| HOY6670 | 0.0302 | 0.9698 | 0.0454 | 0.9502 | 0.0044 | 0.009 | 0.9462 |

| | | | | | | | |
|---|---|---|---|---|---|---|---|
| HOY6672 | 0.0078 | 0.9922 | 0.0192 | 0.9644 | 0.0164 | 0.0072 | 0.966 |
| HOY6673 | 0.0066 | 0.9934 | 0.0104 | 0.9812 | 0.0082 | 0.0026 | 0.9796 |
| HOY6695 | 0.0052 | 0.9948 | 0.0114 | 0.9684 | 0.0204 | 0.0106 | 0.9694 |
| HOY6696 | 0.0372 | 0.9628 | 0.0582 | 0.7558 | 0.1864 | 0.0154 | 0.7516 |
| HOY6697 | 0.0066 | 0.9934 | 0.0138 | 0.8826 | 0.1036 | 0.004 | 0.885 |
| HOY6702 | 0.0052 | 0.9948 | 0.009 | 0.9592 | 0.0316 | 0.0066 | 0.9636 |
| HOY6704 | 0.0062 | 0.9938 | 0.0116 | 0.8594 | 0.129 | 0.0052 | 0.8624 |
| HOY6705 | 0.0116 | 0.9884 | 0.032 | 0.9472 | 0.0208 | 0.0236 | 0.9436 |
| HOY6706 | 0.008 | 0.992 | 0.0148 | 0.8772 | 0.108 | 0.0152 | 0.8834 |
| HOY6708 | 0.0258 | 0.9742 | 0.0658 | 0.8426 | 0.0918 | 0.0222 | 0.8494 |
| HOY6710 | 0.0204 | 0.9796 | 0.0238 | 0.9726 | 0.0034 | 0.0088 | 0.9704 |
| HOY6711 | 0.0782 | 0.9218 | 0.1008 | 0.8954 | 0.0042 | 0.028 | 0.894 |
| HOY6714 | 0.0142 | 0.9858 | 0.0408 | 0.8844 | 0.075 | 0.0884 | 0.8636 |
| HOY6715 | 0.0106 | 0.9894 | 0.0232 | 0.9572 | 0.0196 | 0.0158 | 0.957 |
| HOY6718 | 0.021 | 0.979 | 0.0388 | 0.9528 | 0.0078 | 0.0258 | 0.9484 |
| HOY6720 | 0.0036 | 0.9964 | 0.0066 | 0.8876 | 0.106 | 0.0036 | 0.892 |
| HOY6722 | 0.0578 | 0.9422 | 0.0842 | 0.722 | 0.1936 | 0.0326 | 0.7266 |
| HOY6723 | 0.0482 | 0.9518 | 0.0756 | 0.8796 | 0.0444 | 0.0672 | 0.877 |
| HOY6728 | 0.0154 | 0.9846 | 0.028 | 0.8722 | 0.1 | 0.0786 | 0.8486 |
| HOY6730 | 0.0026 | 0.9974 | 0.004 | 0.8734 | 0.1224 | 0.0032 | 0.8768 |
| HOY6733 | 0.007 | 0.993 | 0.0114 | 0.8382 | 0.1504 | 0.0062 | 0.8416 |
| HOY6734 | 0.059 | 0.941 | 0.0672 | 0.6748 | 0.2582 | 0.1052 | 0.6502 |
| HOY6735 | 0.0952 | 0.9048 | 0.1224 | 0.7412 | 0.1362 | 0.1496 | 0.7358 |
| HOY6736 | 0.2366 | 0.7634 | 0.255 | 0.6956 | 0.0494 | 0.0232 | 0.6962 |
| HOY6737 | 0.0214 | 0.9786 | 0.0438 | 0.9218 | 0.0348 | 0.0468 | 0.915 |
| HOY6738 | 0.0174 | 0.9826 | 0.037 | 0.9246 | 0.038 | 0.0964 | 0.8776 |
| HOY6739 | 0.005 | 0.995 | 0.0076 | 0.7842 | 0.2082 | 0.0062 | 0.7894 |
| HOY6740 | 0.008 | 0.992 | 0.0138 | 0.8102 | 0.176 | 0.037 | 0.8026 |
| HOY6741 | 0.2062 | 0.7938 | 0.2132 | 0.67 | 0.1166 | 0.2276 | 0.6838 |
| HOY6742 | 0.0592 | 0.9408 | 0.0566 | 0.5224 | 0.4208 | 0.0104 | 0.5156 |
| HOY6744 | 0.0216 | 0.9784 | 0.0512 | 0.8558 | 0.093 | 0.0482 | 0.8538 |
| HOY6745 | 0.0074 | 0.9926 | 0.0134 | 0.8702 | 0.1166 | 0.004 | 0.8712 |
| HOY6746 | 0.1506 | 0.8494 | 0.1818 | 0.8016 | 0.0166 | 0.0222 | 0.8062 |
| HOY6747 | 0.01 | 0.99 | 0.0176 | 0.8014 | 0.1806 | 0.0786 | 0.7742 |
| HOY6748 | 0.0056 | 0.9944 | 0.0128 | 0.9626 | 0.0244 | 0.0058 | 0.9656 |
| HOY6749 | 0.0078 | 0.9922 | 0.0132 | 0.7916 | 0.195 | 0.007 | 0.7952 |
| HOY6750 | 0.01 | 0.99 | 0.019 | 0.9502 | 0.0304 | 0.1052 | 0.8806 |
| HOY6751 | 0.036 | 0.964 | 0.0892 | 0.8944 | 0.0166 | 0.0442 | 0.898 |
| HOY6752 | 0.0182 | 0.9818 | 0.0248 | 0.9692 | 0.006 | 0.0304 | 0.9554 |
| HOY6755 | 0.0986 | 0.9014 | 0.1546 | 0.777 | 0.0682 | 0.0246 | 0.7754 |
| HOY6756 | 0.043 | 0.957 | 0.0756 | 0.9144 | 0.0096 | 0.0332 | 0.9132 |
| HOY6757 | 0.0036 | 0.9964 | 0.0056 | 0.8838 | 0.1104 | 0.0084 | 0.8916 |
| HOY6758 | 0.0026 | 0.9974 | 0.005 | 0.9872 | 0.008 | 0.0026 | 0.9884 |
| HOY6759 | 0.0194 | 0.9806 | 0.0274 | 0.9694 | 0.0034 | 0.011 | 0.968 |

| | | | | | | | |
|---|---|---|---|---|---|---|---|
| HOY6761 | 0.0874 | 0.9126 | 0.1274 | 0.7666 | 0.1062 | 0.0788 | 0.77 |
| HOY6762 | 0.0174 | 0.9826 | 0.0496 | 0.8954 | 0.0552 | 0.0354 | 0.8986 |
| HOY6763 | 0.0782 | 0.9218 | 0.1146 | 0.7732 | 0.1122 | 0.0206 | 0.7774 |
| HOY6764 | 0.0052 | 0.9948 | 0.009 | 0.9802 | 0.0108 | 0.0074 | 0.98 |
| HOY6766 | 0.008 | 0.992 | 0.0122 | 0.8428 | 0.1448 | 0.018 | 0.8456 |
| HOY6777 | 0.0202 | 0.9798 | 0.035 | 0.8706 | 0.0944 | 0.1326 | 0.8178 |
| HOY6792 | 0.0048 | 0.9952 | 0.0062 | 0.9908 | 0.003 | 0.003 | 0.99 |
| HOY6793 | 0.0026 | 0.9974 | 0.0036 | 0.9936 | 0.0034 | 0.0018 | 0.993 |
| HOY6795 | 0.0026 | 0.9974 | 0.0036 | 0.9748 | 0.022 | 0.003 | 0.9798 |
| HOY6796 | 0.0062 | 0.9938 | 0.0092 | 0.7024 | 0.2886 | 0.0044 | 0.6998 |
| HOY6798 | 0.0546 | 0.9454 | 0.0498 | 0.6022 | 0.348 | 0.0304 | 0.6 |
| HOY6800 | 0.0114 | 0.9886 | 0.015 | 0.6446 | 0.3404 | 0.0214 | 0.639 |
| HOY6801 | 0.015 | 0.985 | 0.0254 | 0.8022 | 0.172 | 0.0184 | 0.801 |
| HOY6803 | 0.005 | 0.995 | 0.007 | 0.98 | 0.0132 | 0.006 | 0.9812 |
| HOY6805 | 0.134 | 0.866 | 0.1582 | 0.7104 | 0.1314 | 0.015 | 0.7056 |
| HOY6806 | 0.0718 | 0.9282 | 0.0624 | 0.5668 | 0.3708 | 0.0294 | 0.5634 |
| HOY6808 | 0.0104 | 0.9896 | 0.0178 | 0.9716 | 0.0108 | 0.0204 | 0.964 |
| HOY6810 | 0.0156 | 0.9844 | 0.0292 | 0.9664 | 0.0044 | 0.0512 | 0.9374 |
| HOY6811 | 0.0756 | 0.9244 | 0.122 | 0.8572 | 0.0208 | 0.0354 | 0.861 |
| HOY6813 | 0.0036 | 0.9964 | 0.0068 | 0.9614 | 0.032 | 0.0042 | 0.9694 |
| HOY6815 | 0.0052 | 0.9948 | 0.0102 | 0.9426 | 0.0472 | 0.0068 | 0.9494 |
| HOY6817 | 0.0302 | 0.9698 | 0.0704 | 0.8236 | 0.1062 | 0.0228 | 0.8268 |
| HOY6818 | 0.1462 | 0.8538 | 0.1864 | 0.7406 | 0.0728 | 0.1256 | 0.7512 |
| HOY6822 | 0.0136 | 0.9864 | 0.0244 | 0.8616 | 0.1138 | 0.0168 | 0.865 |
| HOY6823 | 0.0026 | 0.9974 | 0.0048 | 0.9286 | 0.0668 | 0.0136 | 0.9368 |
| HOY6828 | 0.0094 | 0.9906 | 0.018 | 0.9502 | 0.032 | 0.0692 | 0.9116 |
| HOY6830 | 0.0128 | 0.9872 | 0.0306 | 0.957 | 0.0122 | 0.014 | 0.958 |
| HOY6836 | 0.0128 | 0.9872 | 0.0188 | 0.9762 | 0.005 | 0.0088 | 0.974 |
| HOY6839 | 0.013 | 0.987 | 0.0378 | 0.9116 | 0.0506 | 0.036 | 0.9156 |
| HOY6840 | 0.012 | 0.988 | 0.018 | 0.9762 | 0.006 | 0.0054 | 0.9738 |
| HOY6841 | 0.057 | 0.943 | 0.0728 | 0.9182 | 0.0092 | 0.0412 | 0.9158 |
| HOY6842 | 0.0062 | 0.9938 | 0.0102 | 0.9342 | 0.0556 | 0.0158 | 0.9424 |
| HOY6843 | 0.0036 | 0.9964 | 0.005 | 0.9916 | 0.0034 | 0.004 | 0.9908 |
| HOY6844 | 0.0128 | 0.9872 | 0.024 | 0.9688 | 0.0072 | 0.0154 | 0.9644 |
| HOY6846 | 0.032 | 0.968 | 0.0564 | 0.9378 | 0.0058 | 0.0564 | 0.923 |
| HOY6847 | 0.0068 | 0.9932 | 0.0108 | 0.8234 | 0.1654 | 0.007 | 0.8286 |
| HOY6848 | 0.0284 | 0.9716 | 0.0472 | 0.9282 | 0.0246 | 0.05 | 0.9174 |
| HOY6849 | 0.0114 | 0.9886 | 0.0204 | 0.947 | 0.0326 | 0.046 | 0.9298 |
| HOY6952 | 0.006 | 0.994 | 0.0116 | 0.9316 | 0.057 | 0.0086 | 0.9346 |
| HOY6953 | 0.0554 | 0.9446 | 0.1016 | 0.7992 | 0.0994 | 0.0308 | 0.8018 |
| HOY6954 | 0.0566 | 0.9434 | 0.1078 | 0.8638 | 0.0282 | 0.058 | 0.8672 |
| HOY6956 | 0.011 | 0.989 | 0.0374 | 0.9114 | 0.051 | 0.017 | 0.9184 |
| HOY6963 | 0.0194 | 0.9806 | 0.0344 | 0.9234 | 0.0422 | 0.0462 | 0.9104 |
| HOY6966 | 0.0118 | 0.9882 | 0.0224 | 0.9514 | 0.0262 | 0.0108 | 0.9544 |

| | | | | | | | |
|---|---|---|---|---|---|---|---|
| HOY6967 | 0.0052 | 0.9948 | 0.0082 | 0.8136 | 0.1776 | 0.007 | 0.8172 |
| HOY6968 | 0.0134 | 0.9866 | 0.032 | 0.9506 | 0.0176 | 0.0526 | 0.928 |
| HOY6970 | 0.0046 | 0.9954 | 0.0092 | 0.9764 | 0.0148 | 0.0064 | 0.9778 |
| HOY6974 | 0.0036 | 0.9964 | 0.005 | 0.9848 | 0.0102 | 0.006 | 0.9848 |
| HOY6975 | 0.0088 | 0.9912 | 0.0202 | 0.9716 | 0.008 | 0.0066 | 0.9708 |
| HOY6978 | 0.0142 | 0.9858 | 0.0292 | 0.934 | 0.037 | 0.0724 | 0.9028 |
| HOY6979 | 0.016 | 0.984 | 0.0382 | 0.8476 | 0.1146 | 0.0064 | 0.8416 |
| HOY6980 | 0.0232 | 0.9768 | 0.0524 | 0.9176 | 0.03 | 0.0118 | 0.9166 |
| HOY6981 | 0.1206 | 0.8794 | 0.1264 | 0.5846 | 0.289 | 0.0658 | 0.5936 |
| HOY6982 | 0.0278 | 0.9722 | 0.0256 | 0.5612 | 0.4132 | 0.0166 | 0.558 |
| HOY6983 | 0.009 | 0.991 | 0.011 | 0.9842 | 0.0048 | 0.0062 | 0.9824 |
| HOY6984 | 0.0122 | 0.9878 | 0.0216 | 0.9648 | 0.014 | 0.0148 | 0.9646 |
| HOY6985 | 0.009 | 0.991 | 0.0152 | 0.9804 | 0.0044 | 0.0048 | 0.9786 |
| HOY6986 | 0.209 | 0.791 | 0.2392 | 0.6882 | 0.0728 | 0.099 | 0.7004 |
| HOY6987 | 0.0192 | 0.9808 | 0.038 | 0.8258 | 0.1364 | 0.012 | 0.8274 |
| HOY6991 | 0.0046 | 0.9954 | 0.0098 | 0.9828 | 0.0078 | 0.0048 | 0.9826 |
| HOY6992 | 0.0218 | 0.9782 | 0.0472 | 0.9454 | 0.0078 | 0.0476 | 0.93 |
| HOY6996 | 0.0072 | 0.9928 | 0.0138 | 0.9556 | 0.0304 | 0.0066 | 0.955 |
| HOY6998 | 0.0048 | 0.9952 | 0.0076 | 0.762 | 0.2302 | 0.0038 | 0.7634 |
| HOY6999 | 0.1482 | 0.8518 | 0.1428 | 0.5672 | 0.2902 | 0.1506 | 0.5594 |
| HOY7000 | 0.0532 | 0.9468 | 0.0964 | 0.8854 | 0.0178 | 0.065 | 0.882 |
| HOY7001 | 0.0672 | 0.9328 | 0.116 | 0.8472 | 0.037 | 0.017 | 0.8454 |
| HOY7002 | 0.0116 | 0.9884 | 0.019 | 0.8648 | 0.1168 | 0.0078 | 0.8656 |
| HOY7012 | 0.0156 | 0.9844 | 0.0268 | 0.7882 | 0.185 | 0.0074 | 0.7876 |
| HOY7013 | 0.1886 | 0.8114 | 0.2046 | 0.789 | 0.0066 | 0.075 | 0.7968 |
| HOY7016 | 0.0098 | 0.9902 | 0.0206 | 0.971 | 0.0084 | 0.0406 | 0.9476 |
| HOY7017 | 0.0182 | 0.9818 | 0.0272 | 0.9666 | 0.0064 | 0.0078 | 0.9634 |
| HOY7019 | 0.0126 | 0.9874 | 0.0254 | 0.8828 | 0.0912 | 0.0226 | 0.8846 |
| HOY7026 | 0.0424 | 0.9576 | 0.0868 | 0.851 | 0.0616 | 0.0794 | 0.851 |
| HOY7034 | 0.253 | 0.747 | 0.2614 | 0.6502 | 0.0886 | 0.0398 | 0.6572 |
| HOY7035 | 0.0174 | 0.9826 | 0.0478 | 0.9048 | 0.0474 | 0.0118 | 0.9096 |
| HOY7036 | 0.0082 | 0.9918 | 0.0168 | 0.9268 | 0.0566 | 0.0064 | 0.9338 |
| HOY7038 | 0.5154 | 0.4846 | 0.5066 | 0.4856 | 0.0076 | 0.031 | 0.488 |
| HOY7039 | 0.0158 | 0.9842 | 0.03 | 0.9634 | 0.0066 | 0.0592 | 0.9296 |
| HOY7043 | 0.3166 | 0.6834 | 0.2972 | 0.6986 | 0.004 | 0.0158 | 0.6922 |
| HOY7044 | 0.0078 | 0.9922 | 0.0202 | 0.932 | 0.0476 | 0.0082 | 0.9402 |
| HOY7045 | 0.0114 | 0.9886 | 0.019 | 0.9714 | 0.0096 | 0.0042 | 0.966 |
| HOY7046 | 0.027 | 0.973 | 0.0686 | 0.8998 | 0.0316 | 0.0094 | 0.89 |
| HOY7047 | 0.019 | 0.981 | 0.044 | 0.9478 | 0.0078 | 0.0086 | 0.9382 |
| HOY7048 | 0.0074 | 0.9926 | 0.0118 | 0.9752 | 0.0128 | 0.007 | 0.9754 |
| HOY7049 | 0.0258 | 0.9742 | 0.008 | 0.0472 | 0.945 | 0.0046 | 0.0372 |
| HOY7054 | 0.0052 | 0.9948 | 0.0078 | 0.9818 | 0.0106 | 0.0066 | 0.9824 |
| HOY7055 | 0.0128 | 0.9872 | 0.0226 | 0.924 | 0.0532 | 0.0098 | 0.9306 |
| HOY7056 | 0.0186 | 0.9814 | 0.054 | 0.933 | 0.0132 | 0.0084 | 0.9248 |

| | | | | | | | |
|---|---|---|---|---|---|---|---|
| HOY7058 | 0.0102 | 0.9898 | 0.017 | 0.7346 | 0.2484 | 0.0192 | 0.7302 |
| HOY7059 | 0.0124 | 0.9876 | 0.0222 | 0.9682 | 0.0094 | 0.0146 | 0.965 |
| HOY7060 | 0.0446 | 0.9554 | 0.0762 | 0.7842 | 0.1396 | 0.0186 | 0.7888 |
| HOY7061 | 0.0054 | 0.9946 | 0.0106 | 0.9458 | 0.0434 | 0.0094 | 0.9528 |
| HOY7062 | 0.0136 | 0.9864 | 0.0264 | 0.965 | 0.0082 | 0.0296 | 0.954 |
| HOY7063 | 0.0792 | 0.9208 | 0.0878 | 0.9066 | 0.006 | 0.012 | 0.8982 |
| HOY7064 | 0.0122 | 0.9878 | 0.0244 | 0.9622 | 0.014 | 0.007 | 0.961 |
| HOY7065 | 0.0546 | 0.9454 | 0.0526 | 0.5812 | 0.3662 | 0.0396 | 0.579 |
| HOY7066 | 0.01 | 0.99 | 0.0314 | 0.9504 | 0.0184 | 0.018 | 0.9496 |
| HOY7067 | 0.1416 | 0.8584 | 0.1698 | 0.817 | 0.0132 | 0.0232 | 0.8164 |
| HOY7068 | 0.038 | 0.962 | 0.083 | 0.9 | 0.0168 | 0.0872 | 0.884 |
| HOY7069 | 0.1626 | 0.8374 | 0.1944 | 0.7938 | 0.0118 | 0.0474 | 0.803 |
| HOY7070 | 0.0098 | 0.9902 | 0.0122 | 0.9846 | 0.0034 | 0.0132 | 0.978 |
| HOY7071 | 0.329 | 0.671 | 0.3302 | 0.519 | 0.151 | 0.0292 | 0.5156 |
| HOY7072 | 0.0194 | 0.9806 | 0.0386 | 0.9292 | 0.0322 | 0.0198 | 0.9316 |
| HOY7073 | 0.0072 | 0.9928 | 0.0122 | 0.9668 | 0.0212 | 0.0228 | 0.9584 |
| HOY7074 | 0.1738 | 0.8262 | 0.216 | 0.7696 | 0.0142 | 0.1332 | 0.7758 |
| HOY7075 | 0.043 | 0.957 | 0.0754 | 0.915 | 0.0094 | 0.0426 | 0.912 |
| HOY7077 | 0.0362 | 0.9638 | 0.0926 | 0.895 | 0.0122 | 0.0716 | 0.8892 |
| HOY7078 | 0.0528 | 0.9472 | 0.098 | 0.8952 | 0.0068 | 0.079 | 0.8888 |
| HOY7080 | 0.062 | 0.938 | 0.0798 | 0.672 | 0.248 | 0.015 | 0.6744 |
| HOY7082 | 0.0088 | 0.9912 | 0.0114 | 0.6672 | 0.3214 | 0.0112 | 0.6678 |
| HOY7083 | 0.0274 | 0.9726 | 0.0416 | 0.8044 | 0.1542 | 0.068 | 0.7936 |
| HOY7084 | 0.0168 | 0.9832 | 0.0506 | 0.9402 | 0.0094 | 0.0342 | 0.9366 |
| HOY7085 | 0.0202 | 0.9798 | 0.0204 | 0.5442 | 0.4352 | 0.0196 | 0.5404 |
| HOY7086 | 0.0184 | 0.9816 | 0.0286 | 0.9578 | 0.0138 | 0.0162 | 0.9564 |
| HOY7087 | 0.0498 | 0.9502 | 0.0904 | 0.8696 | 0.0402 | 0.0186 | 0.8738 |
| HOY7088 | 0.0122 | 0.9878 | 0.0226 | 0.8624 | 0.115 | 0.011 | 0.8664 |
| HOY7089 | 0.0046 | 0.9954 | 0.0064 | 0.9878 | 0.0058 | 0.0038 | 0.9874 |
| HOY7090 | 0.0042 | 0.9958 | 0.0048 | 0.9922 | 0.0032 | 0.0032 | 0.9914 |
| HOY7110 | 0.0058 | 0.9942 | 0.0094 | 0.9754 | 0.0152 | 0.0042 | 0.9766 |
| HOY7111 | 0.2312 | 0.7688 | 0.2154 | 0.365 | 0.4198 | 0.0262 | 0.3604 |
| HOY7112 | 0.4478 | 0.5522 | 0.4424 | 0.5492 | 0.0086 | 0.0068 | 0.5298 |
| HOY7113 | 0.1926 | 0.8074 | 0.2292 | 0.7364 | 0.0342 | 0.0408 | 0.7426 |
| HOY7114 | 0.0582 | 0.9418 | 0.057 | 0.5578 | 0.3852 | 0.0262 | 0.5636 |
| HOY7115 | 0.0072 | 0.9928 | 0.0114 | 0.8032 | 0.1854 | 0.0032 | 0.8024 |
| HOY7117 | 0.0044 | 0.9956 | 0.0076 | 0.9696 | 0.0228 | 0.004 | 0.976 |
| HOY7118 | 0.0182 | 0.9818 | 0.024 | 0.9706 | 0.0052 | 0.01 | 0.969 |
| HOY7119 | 0.0608 | 0.9392 | 0.0658 | 0.5782 | 0.356 | 0.0706 | 0.5702 |
| HOY7120 | 0.014 | 0.986 | 0.025 | 0.8398 | 0.1354 | 0.0166 | 0.8428 |
| HOY7121 | 0.0164 | 0.9836 | 0.032 | 0.9198 | 0.0482 | 0.0154 | 0.9288 |
| HOY7122 | 0.0112 | 0.9888 | 0.0208 | 0.9756 | 0.0036 | 0.0104 | 0.973 |
| HOY7123 | 0.005 | 0.995 | 0.0076 | 0.9854 | 0.0068 | 0.0068 | 0.985 |
| HOY7124 | 0.0242 | 0.9758 | 0.0462 | 0.8606 | 0.0932 | 0.0118 | 0.8608 |

| | | | | | | | |
|---|---|---|---|---|---|---|---|
| HOY7125 | 0.005 | 0.995 | 0.008 | 0.887 | 0.1048 | 0.0078 | 0.8956 |
| HOY7126 | 0.0136 | 0.9864 | 0.0176 | 0.979 | 0.0034 | 0.0054 | 0.9766 |
| HOY7127 | 0.096 | 0.904 | 0.1206 | 0.731 | 0.148 | 0.0466 | 0.7352 |
| HOY7128 | 0.0172 | 0.9828 | 0.0222 | 0.6714 | 0.3064 | 0.0196 | 0.669 |
| HOY7129 | 0.2116 | 0.7884 | 0.2184 | 0.6068 | 0.175 | 0.0054 | 0.5864 |
| HOY7130 | 0.0144 | 0.9856 | 0.0268 | 0.9588 | 0.014 | 0.0226 | 0.9546 |
| HOY7131 | 0.007 | 0.993 | 0.011 | 0.757 | 0.2318 | 0.0158 | 0.755 |
| HOY7132 | 0.0532 | 0.9468 | 0.0442 | 0.4614 | 0.4944 | 0.015 | 0.4608 |
| HOY7134 | 0.0668 | 0.9332 | 0.0696 | 0.6178 | 0.3126 | 0.0626 | 0.614 |
| HOY7135 | 0.0764 | 0.9236 | 0.0902 | 0.468 | 0.4418 | 0.0428 | 0.4728 |
| HOY7136 | 0.0078 | 0.9922 | 0.0108 | 0.703 | 0.2858 | 0.014 | 0.7014 |
| HOY7137 | 0.0042 | 0.9958 | 0.0066 | 0.772 | 0.2214 | 0.0046 | 0.7744 |
| KIY3488 | 0.5614 | 0.4386 | 0.532 | 0.1976 | 0.2702 | 0.0064 | 0.1748 |
| KIY3489 | 0.0098 | 0.9902 | 0.02 | 0.9718 | 0.0082 | 0.0192 | 0.9658 |
| KIY3491 | 0.1946 | 0.8054 | 0.1834 | 0.8122 | 0.0042 | 0.0114 | 0.803 |
| KIY3492 | 0.0074 | 0.9926 | 0.0136 | 0.977 | 0.0096 | 0.008 | 0.977 |
| KIY3494 | 0.9864 | 0.0136 | 0.9542 | 0.0418 | 0.004 | 0.19 | 0.0228 |
| KIY3495 | 0.918 | 0.082 | 0.8816 | 0.03 | 0.0886 | 0.0134 | 0.0136 |
| KIY3497 | 0.9934 | 0.0066 | 0.9634 | 0.0102 | 0.026 | 0.0182 | 0.0056 |
| KIY3498 | 0.0042 | 0.9958 | 0.0068 | 0.9404 | 0.0528 | 0.0066 | 0.9514 |
| KIY3499 | 0.995 | 0.005 | 0.9794 | 0.007 | 0.0138 | 0.0198 | 0.0046 |
| KIY3500 | 0.924 | 0.076 | 0.8736 | 0.123 | 0.0036 | 0.0062 | 0.0884 |
| KIY3502 | 0.9524 | 0.0476 | 0.9034 | 0.0462 | 0.0504 | 0.0074 | 0.0186 |
| KIY3503 | 0.948 | 0.052 | 0.91 | 0.0462 | 0.0442 | 0.049 | 0.028 |
| KIY3504 | 0.915 | 0.085 | 0.8686 | 0.121 | 0.0104 | 0.013 | 0.0864 |
| KIY3505 | 0.9886 | 0.0114 | 0.9608 | 0.0306 | 0.0086 | 0.0162 | 0.0202 |
| KIY3507 | 0.9968 | 0.0032 | 0.9888 | 0.0088 | 0.0024 | 0.1072 | 0.0048 |
| KIY3509 | 0.9944 | 0.0056 | 0.9554 | 0.0062 | 0.0384 | 0.2146 | 0.0038 |
| KIY3511 | 0.996 | 0.004 | 0.9848 | 0.0132 | 0.0022 | 0.0112 | 0.0048 |
| KIY3514 | 0.0376 | 0.9624 | 0.0448 | 0.9516 | 0.0036 | 0.023 | 0.9474 |
| KIY3516 | 0.9806 | 0.0194 | 0.9402 | 0.0566 | 0.0034 | 0.0308 | 0.0312 |
| KIY3518 | 0.976 | 0.024 | 0.9392 | 0.0336 | 0.0266 | 0.0234 | 0.0156 |
| KIY3521 | 0.9656 | 0.0344 | 0.8888 | 0.1078 | 0.0034 | 0.01 | 0.0662 |
| KIY3522 | 0.988 | 0.012 | 0.96 | 0.0292 | 0.0112 | 0.0942 | 0.0176 |
| KIY3533 | 0.9888 | 0.0112 | 0.9684 | 0.0252 | 0.0066 | 0.0372 | 0.0142 |
| KIY3534 | 0.5286 | 0.4714 | 0.5284 | 0.445 | 0.0264 | 0.2896 | 0.435 |
| KIY3535 | 0.9932 | 0.0068 | 0.9768 | 0.013 | 0.0102 | 0.0086 | 0.0072 |
| KIY3537 | 0.991 | 0.009 | 0.9676 | 0.0154 | 0.017 | 0.0386 | 0.0074 |
| KIY3538 | 0.9944 | 0.0056 | 0.9806 | 0.0122 | 0.0072 | 0.1496 | 0.007 |
| KIY3542 | 0.9974 | 0.0026 | 0.9862 | 0.0036 | 0.0102 | 0.015 | 0.002 |
| KIY3543 | 0.9836 | 0.0164 | 0.9322 | 0.0056 | 0.0626 | 0.0294 | 0.003 |
| KIY3544 | 0.9878 | 0.0122 | 0.9638 | 0.0168 | 0.0198 | 0.0082 | 0.0066 |
| KIY3545 | 0.9944 | 0.0056 | 0.9736 | 0.0238 | 0.0024 | 0.0278 | 0.013 |
| KIY3547 | 0.966 | 0.034 | 0.9178 | 0.0656 | 0.0168 | 0.018 | 0.0346 |

| | | | | | | | |
|---|---|---|---|---|---|---|---|
| KIY3548 | 0.944 | 0.056 | 0.9176 | 0.037 | 0.0454 | 0.0202 | 0.026 |
| KIY3550 | 0.9854 | 0.0146 | 0.9454 | 0.0112 | 0.0434 | 0.0122 | 0.0064 |
| KIY3551 | 0.9956 | 0.0044 | 0.9878 | 0.0086 | 0.0038 | 0.0728 | 0.0054 |
| KIY3553 | 0.9978 | 0.0022 | 0.9914 | 0.0042 | 0.0046 | 0.2376 | 0.0024 |
| KIY3554 | 0.9828 | 0.0172 | 0.95 | 0.0338 | 0.0164 | 0.0808 | 0.018 |
| KIY3555 | 0.9488 | 0.0512 | 0.9 | 0.0576 | 0.0428 | 0.0194 | 0.0546 |
| KIY3556 | 0.996 | 0.004 | 0.9862 | 0.009 | 0.0046 | 0.0634 | 0.0068 |
| KIY3557 | 0.9842 | 0.0158 | 0.9498 | 0.0262 | 0.0242 | 0.0038 | 0.006 |
| KIY3558 | 0.9326 | 0.0674 | 0.8842 | 0.105 | 0.0112 | 0.0678 | 0.0924 |
| KIY3559 | 0.8976 | 0.1024 | 0.856 | 0.1398 | 0.0042 | 0.0152 | 0.1262 |
| KIY3565 | 0.9426 | 0.0574 | 0.883 | 0.11 | 0.007 | 0.0222 | 0.0912 |
| KIY3568 | 0.972 | 0.028 | 0.9254 | 0.0642 | 0.0102 | 0.0374 | 0.0512 |
| KIY3570 | 0.9826 | 0.0174 | 0.9438 | 0.0416 | 0.0146 | 0.0384 | 0.0262 |
| KIY3571 | 0.992 | 0.008 | 0.974 | 0.0202 | 0.0058 | 0.0374 | 0.013 |
| KIY3572 | 0.9956 | 0.0044 | 0.985 | 0.0096 | 0.0056 | 0.1852 | 0.0064 |
| KIY3702 | 0.0036 | 0.9964 | 0.0062 | 0.981 | 0.013 | 0.0088 | 0.9794 |
| KIY3703 | 0.9952 | 0.0048 | 0.9832 | 0.0102 | 0.0062 | 0.0148 | 0.005 |
| KIY3706 | 0.0082 | 0.9918 | 0.018 | 0.9666 | 0.0154 | 0.0128 | 0.966 |
| KIY3707 | 0.8512 | 0.1488 | 0.8138 | 0.1786 | 0.0074 | 0.014 | 0.16 |
| KIY3710 | 0.9948 | 0.0052 | 0.9824 | 0.0124 | 0.0056 | 0.2302 | 0.0062 |
| KIY3711 | 0.9868 | 0.0132 | 0.9196 | 0.0096 | 0.0702 | 0.0116 | 0.0064 |
| KIY3712 | 0.998 | 0.002 | 0.9896 | 0.003 | 0.0076 | 0.0066 | 0.002 |
| KIY3713 | 0.8568 | 0.1432 | 0.8178 | 0.0826 | 0.1002 | 0.0194 | 0.0494 |
| KIY3715 | 0.9834 | 0.0166 | 0.9486 | 0.039 | 0.013 | 0.0098 | 0.0104 |
| KIY3716 | 0.9978 | 0.0022 | 0.9888 | 0.003 | 0.0082 | 0.0346 | 0.002 |
| KIY3717 | 0.9876 | 0.0124 | 0.9584 | 0.0334 | 0.0082 | 0.0344 | 0.0254 |
| KIY3718 | 0.9192 | 0.0808 | 0.8914 | 0.03 | 0.0786 | 0.018 | 0.0226 |
| KIY3719 | 0.9574 | 0.0426 | 0.8962 | 0.099 | 0.0056 | 0.0098 | 0.0636 |
| KIY3726 | 0.922 | 0.078 | 0.881 | 0.076 | 0.0436 | 0.0662 | 0.0558 |
| KIY3729 | 0.9748 | 0.0252 | 0.9324 | 0.0618 | 0.006 | 0.0048 | 0.0224 |
| KIY3732 | 0.9812 | 0.0188 | 0.9428 | 0.0512 | 0.0058 | 0.0056 | 0.018 |
| KIY3733 | 0.005 | 0.995 | 0.0086 | 0.9706 | 0.0212 | 0.0184 | 0.9654 |
| KIY3734 | 0.996 | 0.004 | 0.9832 | 0.008 | 0.009 | 0.007 | 0.004 |
| KIY3737 | 0.9876 | 0.0124 | 0.9578 | 0.0204 | 0.022 | 0.041 | 0.01 |
| KIY3738 | 0.996 | 0.004 | 0.9882 | 0.0082 | 0.0034 | 0.059 | 0.005 |
| KIY3739 | 0.994 | 0.006 | 0.9764 | 0.0106 | 0.0132 | 0.008 | 0.005 |
| KIY3740 | 0.9814 | 0.0186 | 0.9432 | 0.05 | 0.0068 | 0.0136 | 0.0286 |
| KIY3741 | 0.9634 | 0.0366 | 0.9362 | 0.0316 | 0.0324 | 0.0162 | 0.0166 |
| KIY3742 | 0.996 | 0.004 | 0.988 | 0.0086 | 0.0034 | 0.0654 | 0.0048 |
| KIY3743 | 0.994 | 0.006 | 0.9722 | 0.0078 | 0.02 | 0.0804 | 0.0054 |
| KIY3744 | 0.9914 | 0.0086 | 0.9538 | 0.0108 | 0.0356 | 0.0276 | 0.005 |
| KIY3745 | 0.9954 | 0.0046 | 0.982 | 0.0096 | 0.0084 | 0.0202 | 0.0052 |
| KIY3746 | 0.887 | 0.113 | 0.8506 | 0.1364 | 0.0132 | 0.0512 | 0.122 |
| KIY3747 | 0.996 | 0.004 | 0.979 | 0.0052 | 0.0154 | 0.0154 | 0.0034 |

| | | | | | | | |
|---|---|---|---|---|---|---|---|
| KIY3749 | 0.996 | 0.004 | 0.987 | 0.0068 | 0.0068 | 0.005 | 0.0032 |
| KIY3750 | 0.8886 | 0.1114 | 0.8566 | 0.1338 | 0.0098 | 0.0056 | 0.1026 |
| KIY3752 | 0.9922 | 0.0078 | 0.9668 | 0.014 | 0.0192 | 0.087 | 0.006 |
| KIY3753 | 0.9924 | 0.0076 | 0.9736 | 0.0142 | 0.0122 | 0.0398 | 0.0086 |
| KIY3754 | 0.9902 | 0.0098 | 0.9326 | 0.0074 | 0.0602 | 0.03 | 0.0046 |
| KIY3756 | 0.9766 | 0.0234 | 0.9206 | 0.0748 | 0.0046 | 0.026 | 0.041 |
| KIY3757 | 0.9262 | 0.0738 | 0.8884 | 0.0822 | 0.0294 | 0.0078 | 0.0326 |
| KIY3758 | 0.9678 | 0.0322 | 0.918 | 0.0752 | 0.0068 | 0.039 | 0.051 |
| KIY3759 | 0.9912 | 0.0088 | 0.968 | 0.0286 | 0.0034 | 0.0068 | 0.017 |
| KIY3760 | 0.1056 | 0.8944 | 0.1262 | 0.7674 | 0.1066 | 0.0198 | 0.7724 |
| KIY3766 | 0.9946 | 0.0054 | 0.9782 | 0.0108 | 0.0108 | 0.0264 | 0.0056 |
| KIY3767 | 0.9906 | 0.0094 | 0.9684 | 0.0222 | 0.0092 | 0.05 | 0.02 |
| KIY3773 | 0.9112 | 0.0888 | 0.8734 | 0.058 | 0.0684 | 0.017 | 0.0448 |
| KIY3774 | 0.9742 | 0.0258 | 0.9228 | 0.0234 | 0.0536 | 0.0558 | 0.0164 |
| KIY3775 | 0.989 | 0.011 | 0.9636 | 0.0318 | 0.0044 | 0.0536 | 0.0202 |
| KIY3776 | 0.5896 | 0.4104 | 0.5628 | 0.4318 | 0.0052 | 0.1116 | 0.4278 |
| KIY3777 | 0.9874 | 0.0126 | 0.933 | 0.0068 | 0.0608 | 0.034 | 0.0044 |
| KIY3778 | 0.9822 | 0.0178 | 0.9334 | 0.063 | 0.0036 | 0.0152 | 0.039 |
| KIY3779 | 0.9758 | 0.0242 | 0.936 | 0.0588 | 0.0058 | 0.0106 | 0.0294 |
| KIY3780 | 0.986 | 0.014 | 0.9478 | 0.0478 | 0.0044 | 0.1464 | 0.0254 |
| KIY3781 | 0.9958 | 0.0042 | 0.9856 | 0.0078 | 0.0068 | 0.292 | 0.0042 |
| KIY3782 | 0.9922 | 0.0078 | 0.9724 | 0.0236 | 0.0036 | 0.0722 | 0.0128 |
| KIY3783 | 0.9896 | 0.0104 | 0.9264 | 0.0098 | 0.064 | 0.08 | 0.0056 |
| KIY3784 | 0.9934 | 0.0066 | 0.9784 | 0.0132 | 0.008 | 0.0504 | 0.0106 |
| KIY3785 | 0.9952 | 0.0048 | 0.9826 | 0.0128 | 0.0048 | 0.01 | 0.0044 |
| KIY3786 | 0.9728 | 0.0272 | 0.9218 | 0.0688 | 0.0094 | 0.0126 | 0.0646 |
| KIY3787 | 0.8418 | 0.1582 | 0.8218 | 0.1036 | 0.0748 | 0.0054 | 0.0668 |
| KIY3788 | 0.9628 | 0.0372 | 0.9104 | 0.0798 | 0.01 | 0.031 | 0.0556 |
| KIY3789 | 0.989 | 0.011 | 0.958 | 0.0368 | 0.0052 | 0.0046 | 0.011 |
| KIY3790 | 0.9838 | 0.0162 | 0.95 | 0.0152 | 0.0348 | 0.1116 | 0.0126 |
| KIY3792 | 0.9766 | 0.0234 | 0.9392 | 0.0342 | 0.0266 | 0.0068 | 0.0132 |
| KIY3794 | 0.9924 | 0.0076 | 0.969 | 0.0094 | 0.0216 | 0.0182 | 0.005 |
| KIY3795 | 0.9976 | 0.0024 | 0.9916 | 0.004 | 0.0042 | 0.0354 | 0.003 |
| KIY3796 | 0.7186 | 0.2814 | 0.7028 | 0.2844 | 0.0126 | 0.0408 | 0.2786 |
| KIY3797 | 0.9434 | 0.0566 | 0.9034 | 0.0728 | 0.024 | 0.014 | 0.0392 |
| KIY3798 | 0.9584 | 0.0416 | 0.9208 | 0.0718 | 0.0074 | 0.046 | 0.0574 |
| KIY3799 | 0.7568 | 0.2432 | 0.7234 | 0.0774 | 0.1992 | 0.1666 | 0.0566 |
| KIY3800 | 0.996 | 0.004 | 0.98 | 0.0056 | 0.0144 | 0.033 | 0.0036 |
| KIY3801 | 0.986 | 0.014 | 0.9632 | 0.0308 | 0.0058 | 0.0164 | 0.0178 |
| KIY3802 | 0.9926 | 0.0074 | 0.976 | 0.0192 | 0.0046 | 0.0112 | 0.0098 |
| KIY3803 | 0.9664 | 0.0336 | 0.9312 | 0.0612 | 0.0076 | 0.114 | 0.0468 |
| KIY3804 | 0.4938 | 0.5062 | 0.4874 | 0.5044 | 0.0082 | 0.009 | 0.4946 |
| KIY3805 | 0.6784 | 0.3216 | 0.6378 | 0.3594 | 0.003 | 0.009 | 0.3444 |
| KIY3806 | 0.9648 | 0.0352 | 0.9144 | 0.0712 | 0.014 | 0.0134 | 0.0542 |

| | | | | | | | |
|---|---|---|---|---|---|---|---|
| KIY3807 | 0.995 | 0.005 | 0.9768 | 0.0072 | 0.016 | 0.0232 | 0.0042 |
| KIY3808 | 0.887 | 0.113 | 0.8558 | 0.117 | 0.0272 | 0.0134 | 0.1002 |
| KIY3809 | 0.987 | 0.013 | 0.9512 | 0.0174 | 0.0314 | 0.0888 | 0.009 |
| KIY3810 | 0.789 | 0.211 | 0.761 | 0.2138 | 0.025 | 0.206 | 0.2048 |
| KIY3811 | 0.791 | 0.209 | 0.77 | 0.0816 | 0.1484 | 0.1956 | 0.0574 |
| KIY3812 | 0.9914 | 0.0086 | 0.9702 | 0.0192 | 0.0106 | 0.0546 | 0.009 |
| KIY3814 | 0.0206 | 0.9794 | 0.047 | 0.9046 | 0.0482 | 0.0412 | 0.9054 |
| KIY3815 | 0.9934 | 0.0066 | 0.9792 | 0.0162 | 0.0048 | 0.0734 | 0.0096 |
| KIY3816 | 0.9678 | 0.0322 | 0.9246 | 0.0422 | 0.0332 | 0.0136 | 0.0246 |
| KIY3817 | 0.998 | 0.002 | 0.9944 | 0.003 | 0.0024 | 0.0176 | 0.002 |
| KIY3818 | 0.9272 | 0.0728 | 0.883 | 0.0558 | 0.0612 | 0.154 | 0.0346 |
| KIY3819 | 0.9824 | 0.0176 | 0.947 | 0.0394 | 0.0136 | 0.2418 | 0.0214 |
| KIY3820 | 0.9526 | 0.0474 | 0.8944 | 0.101 | 0.0048 | 0.027 | 0.0818 |
| KIY3821 | 0.997 | 0.003 | 0.9906 | 0.006 | 0.0034 | 0.0732 | 0.004 |
| KIY3822 | 0.9952 | 0.0048 | 0.9852 | 0.0092 | 0.0054 | 0.0178 | 0.005 |
| KIY3826 | 0.9 | 0.1 | 0.8566 | 0.1396 | 0.0038 | 0.061 | 0.138 |
| KIY3827 | 0.9268 | 0.0732 | 0.8738 | 0.1226 | 0.0034 | 0.004 | 0.0908 |
| KIY3828 | 0.9642 | 0.0358 | 0.9096 | 0.0332 | 0.0572 | 0.0252 | 0.0108 |
| KIY3829 | 0.0254 | 0.9746 | 0.0498 | 0.9428 | 0.007 | 0.0316 | 0.9392 |
| KIY3830 | 0.7136 | 0.2864 | 0.595 | 0.004 | 0.4008 | 0.0158 | 0.003 |
| KIY3832 | 0.1478 | 0.8522 | 0.1846 | 0.7918 | 0.0236 | 0.0346 | 0.793 |
| KIY3834 | 0.5384 | 0.4616 | 0.5246 | 0.4566 | 0.0186 | 0.0296 | 0.4592 |
| KIY3835 | 0.464 | 0.536 | 0.4692 | 0.503 | 0.0282 | 0.0516 | 0.5032 |
| KIY3837 | 0.0124 | 0.9876 | 0.0218 | 0.8662 | 0.1116 | 0.014 | 0.8708 |
| KIY3838 | 0.0132 | 0.9868 | 0.023 | 0.9704 | 0.0064 | 0.0182 | 0.965 |
| KIY3839 | 0.0066 | 0.9934 | 0.0132 | 0.9578 | 0.0288 | 0.0088 | 0.9624 |
| KIY3840 | 0.0086 | 0.9914 | 0.0118 | 0.9836 | 0.0046 | 0.0044 | 0.9824 |
| KIY3841 | 0.1198 | 0.8802 | 0.1254 | 0.641 | 0.234 | 0.054 | 0.647 |
| KIY3842 | 0.0044 | 0.9956 | 0.0074 | 0.9738 | 0.019 | 0.0036 | 0.9768 |
| KIY3843 | 0.0158 | 0.9842 | 0.0214 | 0.9656 | 0.0132 | 0.0036 | 0.9576 |
| KIY3844 | 0.0162 | 0.9838 | 0.0288 | 0.7962 | 0.1752 | 0.0164 | 0.7974 |
| KIY3845 | 0.0132 | 0.9868 | 0.0208 | 0.9676 | 0.0112 | 0.0132 | 0.966 |
| KIY3846 | 0.9916 | 0.0084 | 0.9688 | 0.0246 | 0.0068 | 0.0136 | 0.0218 |
| KIY3847 | 0.9776 | 0.0224 | 0.9374 | 0.0504 | 0.0122 | 0.0248 | 0.0314 |
| KIY3849 | 0.3372 | 0.6628 | 0.3318 | 0.6614 | 0.0066 | 0.0076 | 0.6516 |
| KIY3850 | 0.429 | 0.571 | 0.4094 | 0.3786 | 0.212 | 0.0178 | 0.3702 |
| KIY3851 | 0.0212 | 0.9788 | 0.0366 | 0.896 | 0.0676 | 0.0358 | 0.8924 |
| KIY3852 | 0.0304 | 0.9696 | 0.0758 | 0.8792 | 0.045 | 0.0292 | 0.8846 |
| KIY3853 | 0.0072 | 0.9928 | 0.0156 | 0.969 | 0.0154 | 0.0108 | 0.9696 |
| KIY3854 | 0.0312 | 0.9688 | 0.0362 | 0.9538 | 0.0102 | 0.047 | 0.9338 |
| KIY3857 | 0.1552 | 0.8448 | 0.1432 | 0.5532 | 0.3036 | 0.0718 | 0.5622 |
| KIY3858 | 0.0524 | 0.9476 | 0.0842 | 0.8664 | 0.0494 | 0.0178 | 0.8692 |
| KIY3860 | 0.0104 | 0.9896 | 0.0218 | 0.8574 | 0.1206 | 0.0078 | 0.8604 |
| KIY3862 | 0.7718 | 0.2282 | 0.7542 | 0.2198 | 0.026 | 0.8604 | 0.1312 |

| | | | | | | | |
|---|---|---|---|---|---|---|---|
| NIG3031 | 0.5098 | 0.4902 | 0.4732 | 0.1984 | 0.3286 | 0.014 | 0.1766 |
| NIG3032 | 0.0392 | 0.9608 | 0.0338 | 0.4834 | 0.4826 | 0.0568 | 0.4656 |
| NIG3033 | 0.1468 | 0.8532 | 0.099 | 0.2492 | 0.6514 | 0.0856 | 0.2426 |
| NIG3071 | 0.7074 | 0.2926 | 0.6932 | 0.2782 | 0.0288 | 0.0082 | 0.2542 |
| NIG3079 | 0.0622 | 0.9378 | 0.1 | 0.79 | 0.11 | 0.0088 | 0.776 |
| NIG3089 | 0.5804 | 0.4196 | 0.5448 | 0.2104 | 0.2452 | 0.085 | 0.2048 |
| NIG3091 | 0.1112 | 0.8888 | 0.1222 | 0.5988 | 0.2784 | 0.0958 | 0.6036 |
| NIG3094 | 0.0088 | 0.9912 | 0.0142 | 0.6964 | 0.2896 | 0.0062 | 0.6968 |
| NIG3097 | 0.1296 | 0.8704 | 0.1492 | 0.6978 | 0.153 | 0.1638 | 0.6946 |
| NIG3104 | 0.993 | 0.007 | 0.9542 | 0.0078 | 0.0376 | 0.0214 | 0.005 |
| NIG3109 | 0.6014 | 0.3986 | 0.5884 | 0.3444 | 0.0674 | 0.0768 | 0.3416 |
| NIG3110 | 0.5568 | 0.4432 | 0.5442 | 0.4054 | 0.0502 | 0.323 | 0.4046 |
| NIG3114 | 0.0126 | 0.9874 | 0.0326 | 0.9472 | 0.02 | 0.0122 | 0.9514 |
| NIG3116 | 0.0078 | 0.9922 | 0.0124 | 0.7578 | 0.2298 | 0.0088 | 0.7566 |
| NIG3119 | 0.0044 | 0.9956 | 0.0064 | 0.7816 | 0.212 | 0.0056 | 0.7836 |
| NIG3123 | 0.2022 | 0.7978 | 0.2134 | 0.6726 | 0.1142 | 0.0624 | 0.683 |
| NIG3124 | 0.0776 | 0.9224 | 0.1274 | 0.7926 | 0.0802 | 0.1124 | 0.7934 |
| NIG3128 | 0.029 | 0.971 | 0.0252 | 0.4704 | 0.5046 | 0.0326 | 0.4604 |
| NIG3129 | 0.0332 | 0.9668 | 0.0394 | 0.55 | 0.4106 | 0.0228 | 0.5482 |
| NIG3130 | 0.029 | 0.971 | 0.0328 | 0.6134 | 0.3538 | 0.0428 | 0.6018 |
| NIG3135 | 0.0062 | 0.9938 | 0.01 | 0.7968 | 0.193 | 0.0158 | 0.796 |
| NIG3136 | 0.006 | 0.994 | 0.013 | 0.8866 | 0.1002 | 0.005 | 0.8962 |
| NIG3137 | 0.0032 | 0.9968 | 0.0048 | 0.822 | 0.1734 | 0.0062 | 0.8262 |
| REI0070 | 0.9948 | 0.0052 | 0.9558 | 0.006 | 0.038 | 0.9878 | 0.002 |
| REI0071 | 0.9808 | 0.0192 | 0.8782 | 0.0064 | 0.115 | 0.8842 | 0.0074 |
| REI0072 | 0.9286 | 0.0714 | 0.8768 | 0.083 | 0.0398 | 0.9242 | 0.053 |
| REI0075 | 0.9902 | 0.0098 | 0.9548 | 0.0168 | 0.0284 | 0.9752 | 0.0088 |
| REI0076 | 0.9904 | 0.0096 | 0.9036 | 0.008 | 0.0882 | 0.9706 | 0.0036 |
| REI0077 | 0.9952 | 0.0048 | 0.9326 | 0.005 | 0.0624 | 0.989 | 0.002 |
| REI0078 | 0.9116 | 0.0884 | 0.799 | 0.0144 | 0.1862 | 0.9724 | 0.0048 |
| REI0080 | 0.9894 | 0.0106 | 0.964 | 0.0258 | 0.0102 | 0.9486 | 0.0116 |
| REI0081 | 0.9924 | 0.0076 | 0.9304 | 0.0074 | 0.0626 | 0.9614 | 0.004 |
| REI0082 | 0.825 | 0.175 | 0.7762 | 0.0814 | 0.1422 | 0.9104 | 0.0608 |
| REI0083 | 0.858 | 0.142 | 0.794 | 0.0468 | 0.1594 | 0.9282 | 0.0504 |
| REI0084 | 0.9884 | 0.0116 | 0.8786 | 0.0072 | 0.114 | 0.9526 | 0.0042 |
| REI0085 | 0.769 | 0.231 | 0.7056 | 0.0692 | 0.225 | 0.9014 | 0.0624 |
| REI0086 | 0.9834 | 0.0166 | 0.8828 | 0.0106 | 0.1066 | 0.9544 | 0.009 |
| REI0087 | 0.99 | 0.01 | 0.7988 | 0.0038 | 0.1972 | 0.98 | 0.002 |
| REI0089 | 0.996 | 0.004 | 0.9844 | 0.006 | 0.0096 | 0.9796 | 0.0028 |
| REI0090 | 0.9882 | 0.0118 | 0.9548 | 0.0258 | 0.0194 | 0.9802 | 0.0112 |
| REI0091 | 0.9292 | 0.0708 | 0.8348 | 0.0064 | 0.1588 | 0.9286 | 0.0042 |
| REI0092 | 0.9914 | 0.0086 | 0.9686 | 0.022 | 0.0094 | 0.8306 | 0.0048 |
| REI0093 | 0.8582 | 0.1418 | 0.8064 | 0.085 | 0.1084 | 0.939 | 0.0472 |
| REI0094 | 0.9774 | 0.0226 | 0.8458 | 0.0058 | 0.1484 | 0.9556 | 0.005 |

| | | | | | | | |
|---|---|---|---|---|---|---|---|
| REI0095 | 0.9786 | 0.0214 | 0.8284 | 0.0074 | 0.164 | 0.9722 | 0.0042 |
| REI0096 | 0.9936 | 0.0064 | 0.968 | 0.0086 | 0.0234 | 0.8264 | 0.01 |
| REI0099 | 0.992 | 0.008 | 0.8848 | 0.0052 | 0.11 | 0.949 | 0.0044 |
| ZEN0212 | 0.0266 | 0.9734 | 0.0442 | 0.8054 | 0.1508 | 0.129 | 0.7614 |
| ZEN0213 | 0.0048 | 0.9952 | 0.0068 | 0.5916 | 0.4012 | 0.0038 | 0.5936 |
| ZEN0217 | 0.0052 | 0.9948 | 0.0078 | 0.9824 | 0.0098 | 0.0056 | 0.9834 |
| ZEN0218 | 0.0046 | 0.9954 | 0.0092 | 0.9274 | 0.0632 | 0.0072 | 0.936 |
| ZEN0219 | 0.1018 | 0.8982 | 0.104 | 0.5356 | 0.3606 | 0.0372 | 0.547 |
| ZEN0225 | 0.0228 | 0.9772 | 0.0318 | 0.6802 | 0.2882 | 0.0102 | 0.6774 |
| ZEN0229 | 0.2906 | 0.7094 | 0.3016 | 0.6894 | 0.0094 | 0.0494 | 0.694 |
| ZEN0230 | 0.0558 | 0.9442 | 0.0908 | 0.7888 | 0.1198 | 0.0218 | 0.7912 |
| ZEN0232 | 0.008 | 0.992 | 0.0214 | 0.9378 | 0.0408 | 0.0098 | 0.9412 |
| ZEN0233 | 0.008 | 0.992 | 0.0132 | 0.8422 | 0.1448 | 0.0118 | 0.8432 |
| ZEN0234 | 0.042 | 0.958 | 0.0696 | 0.7796 | 0.1508 | 0.0388 | 0.7834 |
| ZEN0236 | 0.0068 | 0.9932 | 0.0102 | 0.9848 | 0.0056 | 0.0182 | 0.9738 |
| ZEN0237 | 0.0358 | 0.9642 | 0.0572 | 0.9336 | 0.009 | 0.029 | 0.9332 |
| ZEN0238 | 0.0034 | 0.9966 | 0.0058 | 0.9804 | 0.014 | 0.0032 | 0.9834 |
| ZEN0239 | 0.0036 | 0.9964 | 0.0064 | 0.9718 | 0.0216 | 0.0036 | 0.976 |
| ZEN0240 | 0.0132 | 0.9868 | 0.042 | 0.9394 | 0.0186 | 0.0656 | 0.9148 |
| ZEN0241 | 0.0026 | 0.9974 | 0.004 | 0.9902 | 0.0058 | 0.0036 | 0.9908 |
| ZEN0242 | 0.0072 | 0.9928 | 0.0106 | 0.7474 | 0.242 | 0.0114 | 0.7476 |
| ZEN0243 | 0.0046 | 0.9954 | 0.0068 | 0.7296 | 0.2636 | 0.0036 | 0.7292 |
| ZEN0245 | 0.0508 | 0.9492 | 0.0888 | 0.852 | 0.0594 | 0.1156 | 0.8358 |
| ZEN0246 | 0.0448 | 0.9552 | 0.071 | 0.9182 | 0.0108 | 0.0284 | 0.9186 |
| ZEN0247 | 0.006 | 0.994 | 0.0106 | 0.9812 | 0.0082 | 0.005 | 0.9808 |
| ZEN0249 | 0.9942 | 0.0058 | 0.9816 | 0.0144 | 0.0042 | 0.0086 | 0.008 |
| ZEN0253 | 0.016 | 0.984 | 0.0436 | 0.8776 | 0.0788 | 0.0116 | 0.88 |
| ZEN0254 | 0.041 | 0.959 | 0.0616 | 0.8644 | 0.0744 | 0.0856 | 0.8572 |
| ZEN0255 | 0.0026 | 0.9974 | 0.0036 | 0.9668 | 0.0298 | 0.0028 | 0.9748 |
| ZEN0258 | 0.0102 | 0.9898 | 0.0272 | 0.9554 | 0.0178 | 0.0136 | 0.9578 |
| ZEN0277 | 0.509 | 0.491 | 0.496 | 0.433 | 0.0708 | 0.4974 | 0.4496 |
| ZEN0278 | 0.0352 | 0.9648 | 0.0706 | 0.9038 | 0.0252 | 0.0166 | 0.9002 |
| ZEN0279 | 0.0338 | 0.9662 | 0.086 | 0.9002 | 0.0134 | 0.0276 | 0.9024 |
| ZEN0281 | 0.2852 | 0.7148 | 0.306 | 0.6808 | 0.0134 | 0.2524 | 0.6798 |
| ZEN0282 | 0.3886 | 0.6114 | 0.3886 | 0.605 | 0.0066 | 0.3308 | 0.6028 |
| ZEN0283 | 0.1118 | 0.8882 | 0.1434 | 0.7754 | 0.0812 | 0.056 | 0.7882 |
| ZEN0284 | 0.0836 | 0.9164 | 0.1166 | 0.8798 | 0.004 | 0.098 | 0.8692 |
| ZEN0285 | 0.1422 | 0.8578 | 0.1592 | 0.6868 | 0.154 | 0.0664 | 0.6988 |
| ZEN0289 | 0.1946 | 0.8054 | 0.2128 | 0.6604 | 0.127 | 0.0248 | 0.658 |
| ZEN0290 | 0.006 | 0.994 | 0.0106 | 0.8722 | 0.117 | 0.0276 | 0.8702 |
| ZEN0293 | 0.0184 | 0.9816 | 0.03 | 0.7908 | 0.1794 | 0.0158 | 0.79 |
| ZEN0296 | 0.9576 | 0.0424 | 0.892 | 0.0396 | 0.0684 | 0.8984 | 0.06 |
| ZEN0297 | 0.7096 | 0.2904 | 0.6854 | 0.2156 | 0.0992 | 0.157 | 0.21 |
| ZEN0299 | 0.2316 | 0.7684 | 0.2282 | 0.4874 | 0.2846 | 0.0118 | 0.4776 |

| | | | | | | | |
|---|---|---|---|---|---|---|---|
| ZEN0300 | 0.0882 | 0.9118 | 0.1208 | 0.8686 | 0.0104 | 0.0346 | 0.8674 |
| ZEN0303 | 0.038 | 0.962 | 0.0608 | 0.7772 | 0.1618 | 0.026 | 0.781 |
| ZEN0305 | 0.4638 | 0.5362 | 0.4626 | 0.4856 | 0.0518 | 0.535 | 0.4494 |
| ZEN0317 | 0.998 | 0.002 | 0.988 | 0.003 | 0.009 | 0.8678 | 0.002 |
| ZEN0319 | 0.6078 | 0.3922 | 0.589 | 0.1938 | 0.217 | 0.273 | 0.1894 |
| ZEN0333 | 0.8796 | 0.1204 | 0.8474 | 0.0798 | 0.073 | 0.8998 | 0.0758 |
| ZEN0343 | 0.9928 | 0.0072 | 0.9068 | 0.005 | 0.0878 | 0.9592 | 0.0028 |
| ZEN0349 | 0.9918 | 0.0082 | 0.8896 | 0.0056 | 0.1048 | 0.9536 | 0.003 |
| ZEN0351 | 0.822 | 0.178 | 0.8072 | 0.1746 | 0.0184 | 0.0694 | 0.1672 |
| ZEN0354 | 0.756 | 0.244 | 0.7176 | 0.1678 | 0.1144 | 0.7048 | 0.1848 |
| ZEN0356 | 0.9904 | 0.0096 | 0.9522 | 0.0138 | 0.0338 | 0.9886 | 0.003 |
| ZEN0361 | 0.9956 | 0.0044 | 0.8906 | 0.0034 | 0.1062 | 0.9792 | 0.003 |
| ZEN0365 | 0.9936 | 0.0064 | 0.9564 | 0.0084 | 0.0354 | 0.9824 | 0.004 |
| ZEN0367 | 0.41 | 0.59 | 0.4062 | 0.5222 | 0.0714 | 0.3998 | 0.539 |
| ZEN0378 | 0.3702 | 0.6298 | 0.384 | 0.556 | 0.0602 | 0.4542 | 0.5186 |
| ZEN0379 | 0.9808 | 0.0192 | 0.93 | 0.03 | 0.04 | 0.8608 | 0.0474 |
| ZEN0383 | 0.996 | 0.004 | 0.9644 | 0.0056 | 0.0302 | 0.9664 | 0.0024 |
| ZEN0384 | 0.4754 | 0.5246 | 0.471 | 0.483 | 0.0458 | 0.2916 | 0.4822 |
| ZEN0388 | 0.5198 | 0.4802 | 0.5096 | 0.407 | 0.0834 | 0.016 | 0.3978 |
| ZEN0389 | 0.1126 | 0.8874 | 0.1196 | 0.5348 | 0.3456 | 0.101 | 0.5326 |
| ZEN0390 | 0.0094 | 0.9906 | 0.0172 | 0.8636 | 0.1192 | 0.009 | 0.8652 |
| ZEN0391 | 0.006 | 0.994 | 0.0086 | 0.9874 | 0.0042 | 0.005 | 0.986 |
| ZEN0392 | 0.0116 | 0.9884 | 0.021 | 0.8554 | 0.1236 | 0.0116 | 0.8598 |
| ZEN0393 | 0.0232 | 0.9768 | 0.0676 | 0.9048 | 0.0274 | 0.0122 | 0.9074 |
| ZEN0395 | 0.0524 | 0.9476 | 0.0618 | 0.9336 | 0.0046 | 0.0102 | 0.923 |
| ZEN0397 | 0.015 | 0.985 | 0.0208 | 0.9746 | 0.0046 | 0.0088 | 0.9712 |
| ZEN0398 | 0.194 | 0.806 | 0.2372 | 0.7388 | 0.0234 | 0.0098 | 0.7306 |
| ZEN0399 | 0.029 | 0.971 | 0.0388 | 0.7082 | 0.2532 | 0.064 | 0.6928 |
| ZEN0400 | 0.0224 | 0.9776 | 0.0544 | 0.912 | 0.0338 | 0.0734 | 0.8936 |
| ZEN0402 | 0.0448 | 0.9552 | 0.081 | 0.8034 | 0.1154 | 0.0926 | 0.797 |
| ZEN0405 | 0.0046 | 0.9954 | 0.0074 | 0.799 | 0.1936 | 0.0066 | 0.8032 |
| ZEN0406 | 0.0378 | 0.9622 | 0.0776 | 0.8776 | 0.0448 | 0.039 | 0.8822 |
| ZEN0407 | 0.1584 | 0.8416 | 0.1224 | 0.2776 | 0.5998 | 0.1336 | 0.2652 |
| ZEN0410 | 0.0108 | 0.9892 | 0.0144 | 0.6718 | 0.3138 | 0.0106 | 0.6716 |
| ZEN0411 | 0.0286 | 0.9714 | 0.065 | 0.918 | 0.0168 | 0.0184 | 0.9234 |
| ZEN0412 | 0.0222 | 0.9778 | 0.0402 | 0.9018 | 0.058 | 0.011 | 0.9056 |
| ZEN0413 | 0.0096 | 0.9904 | 0.0134 | 0.6394 | 0.347 | 0.0278 | 0.6312 |
| ZEN0414 | 0.0086 | 0.9914 | 0.0224 | 0.95 | 0.0276 | 0.0286 | 0.9448 |
| ZEN0415 | 0.213 | 0.787 | 0.2472 | 0.7198 | 0.0324 | 0.009 | 0.7128 |
| ZEN0416 | 0.0366 | 0.9634 | 0.0856 | 0.891 | 0.0232 | 0.1126 | 0.8632 |
| ZEN0417 | 0.0134 | 0.9866 | 0.035 | 0.96 | 0.0054 | 0.0136 | 0.956 |
| ZEN0418 | 0.018 | 0.982 | 0.0368 | 0.902 | 0.0614 | 0.0402 | 0.9016 |
| ZEN0420 | 0.0182 | 0.9818 | 0.0422 | 0.9512 | 0.0066 | 0.0092 | 0.9478 |
| ZEN0422 | 0.0994 | 0.9006 | 0.1404 | 0.7408 | 0.1186 | 0.03 | 0.7434 |

| | | | | | | | |
|---|---|---|---|---|---|---|---|
| ZEN0423 | 0.096 | 0.904 | 0.1382 | 0.8268 | 0.0348 | 0.0356 | 0.836 |
| ZEN0424 | 0.004 | 0.996 | 0.007 | 0.9582 | 0.0348 | 0.0044 | 0.9662 |
| ZEN0425 | 0.2324 | 0.7676 | 0.2324 | 0.7638 | 0.004 | 0.0136 | 0.759 |
| ZEN0426 | 0.0326 | 0.9674 | 0.056 | 0.8034 | 0.1406 | 0.013 | 0.802 |
| ZEN0427 | 0.1156 | 0.8844 | 0.1286 | 0.8598 | 0.0116 | 0.018 | 0.855 |
| ZEN0429 | 0.1198 | 0.8802 | 0.1532 | 0.8252 | 0.0216 | 0.0424 | 0.8308 |
| ZEN0435 | 0.0562 | 0.9438 | 0.0874 | 0.8032 | 0.1098 | 0.057 | 0.811 |
| ZEN0436 | 0.0086 | 0.9914 | 0.0114 | 0.9848 | 0.0036 | 0.0114 | 0.9806 |
| ZEN0437 | 0.0096 | 0.9904 | 0.0196 | 0.9078 | 0.0726 | 0.0192 | 0.9172 |
| ZEN0439 | 0.0136 | 0.9864 | 0.031 | 0.8816 | 0.0876 | 0.0438 | 0.8816 |
| ZEN0442 | 0.0758 | 0.9242 | 0.082 | 0.4862 | 0.4314 | 0.0082 | 0.4702 |
| ZEN0444 | 0.054 | 0.946 | 0.0556 | 0.5628 | 0.3818 | 0.048 | 0.556 |
| ZEN0445 | 0.9856 | 0.0144 | 0.8908 | 0.0102 | 0.099 | 0.899 | 0.0056 |
| ZEN0446 | 0.7928 | 0.2072 | 0.747 | 0.1576 | 0.0958 | 0.826 | 0.1262 |
| ZEN0447 | 0.4752 | 0.5248 | 0.4748 | 0.516 | 0.0092 | 0.0976 | 0.5164 |
| ZEN0448 | 0.5276 | 0.4724 | 0.4858 | 0.1836 | 0.3304 | 0.0316 | 0.1732 |
| ZEN0449 | 0.9786 | 0.0214 | 0.8654 | 0.0046 | 0.1302 | 0.9638 | 0.003 |
| ZEN0450 | 0.9936 | 0.0064 | 0.941 | 0.0068 | 0.0522 | 0.98 | 0.0052 |
| ZEN0451 | 0.9888 | 0.0112 | 0.9552 | 0.0204 | 0.0242 | 0.9346 | 0.005 |
| ZEN0452 | 0.0052 | 0.9948 | 0.0064 | 0.4098 | 0.5834 | 0.007 | 0.4092 |
| ZEN0453 | 0.013 | 0.987 | 0.0166 | 0.5972 | 0.3862 | 0.0064 | 0.5978 |
| ZEN0454 | 0.0038 | 0.9962 | 0.0056 | 0.9906 | 0.0034 | 0.0094 | 0.9856 |
| ZEN0455 | 0.9424 | 0.0576 | 0.8784 | 0.039 | 0.0826 | 0.917 | 0.0368 |
| ZEN0457 | 0.9822 | 0.0178 | 0.898 | 0.0136 | 0.0884 | 0.8438 | 0.0186 |
| ZEN0458 | 0.9868 | 0.0132 | 0.8972 | 0.0106 | 0.0922 | 0.9652 | 0.005 |
| ZEN0459 | 0.9834 | 0.0166 | 0.906 | 0.0144 | 0.0794 | 0.9898 | 0.0034 |
| ZEN0462 | 0.0062 | 0.9938 | 0.0128 | 0.9244 | 0.0628 | 0.0086 | 0.9322 |
| ZEN0464 | 0.9886 | 0.0114 | 0.9172 | 0.0116 | 0.0716 | 0.969 | 0.007 |
| ZEN0465 | 0.9702 | 0.0298 | 0.875 | 0.0104 | 0.115 | 0.964 | 0.0062 |
| ZEN0469 | 0.8742 | 0.1258 | 0.8276 | 0.0686 | 0.1036 | 0.8652 | 0.0952 |
| ZEN0470 | 0.9414 | 0.0586 | 0.8902 | 0.0614 | 0.0486 | 0.9202 | 0.0612 |
| ZEN0472 | 0.1692 | 0.8308 | 0.1814 | 0.6112 | 0.2076 | 0.039 | 0.6138 |
| ZEN0475 | 0.0294 | 0.9706 | 0.053 | 0.9412 | 0.0058 | 0.0342 | 0.9354 |

| | | K=5 | | | | | K=6 |
|---|---|---|---|---|---|---|---|
| X4.Cluster3 | X4.Cluster4 | X5.Cluster1 | X5.Cluster2 | X5.Cluster3 | X5.Cluster4 | X5.Cluster5 | X6.Cluster1 |
| 0.5022 | 0.0068 | 0.0952 | 0.4064 | 0.4778 | 0.0066 | 0.014 | 0.2272 |
| 0.7676 | 0.0066 | 0.1796 | 0.067 | 0.6906 | 0.0046 | 0.0582 | 0.1054 |
| 0.4906 | 0.0228 | 0.1344 | 0.3732 | 0.4504 | 0.02 | 0.0214 | 0.156 |
| 0.4762 | 0.0152 | 0.1038 | 0.374 | 0.4542 | 0.0204 | 0.0478 | 0.2252 |
| 0.9594 | 0.0124 | 0.0212 | 0.0348 | 0.9122 | 0.0192 | 0.0126 | 0.026 |
| 0.2502 | 0.018 | 0.0866 | 0.596 | 0.2398 | 0.0214 | 0.0564 | 0.1628 |
| 0.2036 | 0.1104 | 0.0212 | 0.6606 | 0.1942 | 0.1092 | 0.015 | 0.0166 |
| 0.564 | 0.0444 | 0.0168 | 0.385 | 0.5344 | 0.0488 | 0.0144 | 0.0326 |
| 0.7998 | 0.016 | 0.0368 | 0.1756 | 0.757 | 0.019 | 0.0112 | 0.0748 |
| 0.6146 | 0.0058 | 0.1034 | 0.2928 | 0.5592 | 0.004 | 0.0404 | 0.0668 |
| 0.2938 | 0.0124 | 0.1582 | 0.546 | 0.2428 | 0.0064 | 0.0468 | 0.0738 |
| 0.9814 | 0.0062 | 0.0802 | 0.0036 | 0.8872 | 0.0042 | 0.025 | 0.047 |
| 0.769 | 0.0056 | 0.0708 | 0.1864 | 0.7186 | 0.005 | 0.0188 | 0.0924 |
| 0.9534 | 0.0214 | 0.0182 | 0.013 | 0.9114 | 0.035 | 0.022 | 0.018 |
| 0.8108 | 0.0168 | 0.2316 | 0.0048 | 0.6992 | 0.0072 | 0.0572 | 0.116 |
| 0.6768 | 0.1736 | 0.1954 | 0.0416 | 0.611 | 0.1124 | 0.0396 | 0.1232 |
| 0.9368 | 0.026 | 0.006 | 0.015 | 0.8992 | 0.051 | 0.0286 | 0.0074 |
| 0.9578 | 0.0216 | 0.0154 | 0.0208 | 0.9156 | 0.0354 | 0.0126 | 0.02 |
| 0.965 | 0.0054 | 0.0174 | 0.0406 | 0.9212 | 0.0066 | 0.0142 | 0.0158 |
| 0.9608 | 0.0244 | 0.0324 | 0.0102 | 0.9182 | 0.0268 | 0.0128 | 0.011 |
| 0.8496 | 0.0196 | 0.0188 | 0.0728 | 0.8162 | 0.0364 | 0.0556 | 0.0272 |
| 0.2034 | 0.0218 | 0.0814 | 0.6752 | 0.1796 | 0.0182 | 0.0454 | 0.048 |
| 0.7718 | 0.0234 | 0.0434 | 0.1428 | 0.7362 | 0.032 | 0.0456 | 0.0804 |
| 0.6738 | 0.0332 | 0.0288 | 0.0728 | 0.6398 | 0.0716 | 0.187 | 0.0336 |
| 0.7134 | 0.0202 | 0.0658 | 0.1836 | 0.672 | 0.0232 | 0.0552 | 0.052 |
| 0.6984 | 0.0188 | 0.0348 | 0.2634 | 0.659 | 0.0178 | 0.0248 | 0.0342 |
| 0.9876 | 0.0052 | 0.0094 | 0.0058 | 0.9676 | 0.0108 | 0.0062 | 0.0146 |
| 0.8564 | 0.0694 | 0.0126 | 0.0478 | 0.819 | 0.0866 | 0.034 | 0.0136 |
| 0.9332 | 0.0044 | 0.1488 | 0.0068 | 0.803 | 0.0024 | 0.0392 | 0.0802 |
| 0.6648 | 0.2598 | 0.0376 | 0.0726 | 0.625 | 0.254 | 0.0106 | 0.0562 |
| 0.9718 | 0.008 | 0.0122 | 0.0106 | 0.9378 | 0.0186 | 0.0208 | 0.0178 |
| 0.7478 | 0.2134 | 0.0282 | 0.0282 | 0.7058 | 0.2204 | 0.0178 | 0.0254 |
| 0.9894 | 0.005 | 0.0056 | 0.0038 | 0.9734 | 0.0132 | 0.0044 | 0.0086 |
| 0.7942 | 0.0094 | 0.1416 | 0.083 | 0.7246 | 0.006 | 0.045 | 0.085 |
| 0.939 | 0.0314 | 0.1196 | 0.0074 | 0.829 | 0.0112 | 0.033 | 0.0546 |
| 0.9586 | 0.0102 | 0.0064 | 0.0052 | 0.9298 | 0.0254 | 0.0336 | 0.0066 |
| 0.8838 | 0.0188 | 0.1524 | 0.0038 | 0.781 | 0.0144 | 0.0482 | 0.077 |
| 0.7454 | 0.0072 | 0.0176 | 0.2468 | 0.709 | 0.0082 | 0.0184 | 0.009 |
| 0.9132 | 0.0054 | 0.012 | 0.1074 | 0.8662 | 0.0058 | 0.0084 | 0.0086 |
| 0.922 | 0.0246 | 0.1664 | 0.0048 | 0.7826 | 0.0062 | 0.04 | 0.0888 |
| 0.9794 | 0.0116 | 0.01 | 0.0048 | 0.9486 | 0.0294 | 0.0072 | 0.0114 |

| | | | | | | | |
|---|---|---|---|---|---|---|---|
| 0.8908 | 0.0368 | 0.0096 | 0.0034 | 0.852 | 0.0714 | 0.0636 | 0.0052 |
| 0.8764 | 0.007 | 0.0272 | 0.1292 | 0.831 | 0.006 | 0.0066 | 0.0594 |
| 0.782 | 0.0352 | 0.1408 | 0.089 | 0.7202 | 0.0224 | 0.0274 | 0.109 |
| 0.7428 | 0.0044 | 0.1166 | 0.1584 | 0.679 | 0.0034 | 0.0428 | 0.094 |
| 0.9846 | 0.0058 | 0.0642 | 0.0028 | 0.9066 | 0.005 | 0.021 | 0.0366 |
| 0.673 | 0.011 | 0.034 | 0.2704 | 0.6428 | 0.0138 | 0.0382 | 0.0434 |
| 0.9146 | 0.0478 | 0.0458 | 0.0074 | 0.8592 | 0.0648 | 0.0226 | 0.0276 |
| 0.9904 | 0.0028 | 0.0104 | 0.009 | 0.9704 | 0.004 | 0.006 | 0.0126 |
| 0.8804 | 0.0542 | 0.1342 | 0.016 | 0.7974 | 0.0242 | 0.0284 | 0.0806 |
| 0.8754 | 0.1054 | 0.015 | 0.0224 | 0.8336 | 0.1234 | 0.0056 | 0.022 |
| 0.5572 | 0.0036 | 0.0416 | 0.407 | 0.5364 | 0.0042 | 0.0106 | 0.0878 |
| 0.881 | 0.0462 | 0.0424 | 0.0632 | 0.8278 | 0.0518 | 0.0148 | 0.0708 |
| 0.4368 | 0.0186 | 0.037 | 0.4484 | 0.4182 | 0.0258 | 0.0706 | 0.0734 |
| 0.566 | 0.1182 | 0.045 | 0.2826 | 0.5322 | 0.1212 | 0.0192 | 0.06 |
| 0.6528 | 0.0116 | 0.1422 | 0.1128 | 0.6136 | 0.0232 | 0.108 | 0.1152 |
| 0.9764 | 0.0058 | 0.02 | 0.0194 | 0.9394 | 0.01 | 0.0112 | 0.0324 |
| 0.6454 | 0.0066 | 0.067 | 0.2964 | 0.6146 | 0.008 | 0.0134 | 0.156 |
| 0.8512 | 0.0136 | 0.2296 | 0.0036 | 0.7084 | 0.0048 | 0.0536 | 0.117 |
| 0.8386 | 0.0072 | 0.0356 | 0.1346 | 0.803 | 0.0108 | 0.0158 | 0.0758 |
| 0.683 | 0.0108 | 0.0528 | 0.273 | 0.6518 | 0.0112 | 0.0114 | 0.1436 |
| 0.8792 | 0.0084 | 0.0434 | 0.0826 | 0.8332 | 0.0092 | 0.0312 | 0.0354 |
| 0.7534 | 0.0434 | 0.0266 | 0.1696 | 0.7188 | 0.0586 | 0.0266 | 0.0444 |
| 0.984 | 0.0052 | 0.0052 | 0.0104 | 0.9678 | 0.009 | 0.0072 | 0.0086 |
| 0.9068 | 0.049 | 0.0978 | 0.0242 | 0.8368 | 0.0322 | 0.009 | 0.1116 |
| 0.7008 | 0.0034 | 0.0424 | 0.2774 | 0.67 | 0.0034 | 0.0068 | 0.0658 |
| 0.595 | 0.004 | 0.0168 | 0.4092 | 0.559 | 0.0034 | 0.0114 | 0.0198 |
| 0.9752 | 0.0094 | 0.055 | 0.0118 | 0.908 | 0.0096 | 0.0158 | 0.0346 |
| 0.8886 | 0.0058 | 0.0268 | 0.0922 | 0.8548 | 0.0066 | 0.0196 | 0.042 |
| 0.8348 | 0.1156 | 0.0184 | 0.0128 | 0.7974 | 0.147 | 0.0246 | 0.0234 |
| 0.601 | 0.076 | 0.0228 | 0.3124 | 0.57 | 0.0806 | 0.0146 | 0.0232 |
| 0.0394 | 0.005 | 0.1984 | 0.746 | 0.0314 | 0.0046 | 0.02 | 0.5008 |
| 0.129 | 0.0148 | 0.1988 | 0.6658 | 0.1102 | 0.0126 | 0.013 | 0.4678 |
| 0.032 | 0.0092 | 0.1322 | 0.8086 | 0.0278 | 0.0092 | 0.022 | 0.232 |
| 0.2182 | 0.0032 | 0.0278 | 0.7474 | 0.2046 | 0.0028 | 0.0174 | 0.035 |
| 0.0044 | 0.0306 | 0.0762 | 0.865 | 0.003 | 0.0216 | 0.0344 | 0.1504 |
| 0.0034 | 0.0152 | 0.0082 | 0.9524 | 0.003 | 0.0152 | 0.021 | 0.0188 |
| 0.0114 | 0.0154 | 0.2202 | 0.749 | 0.0088 | 0.0118 | 0.0102 | 0.5616 |
| 0.0236 | 0.007 | 0.2326 | 0.6914 | 0.0086 | 0.004 | 0.0634 | 0.2642 |
| 0.3492 | 0.0852 | 0.077 | 0.4866 | 0.3286 | 0.0822 | 0.025 | 0.0984 |
| 0.2246 | 0.0062 | 0.0352 | 0.7358 | 0.215 | 0.0062 | 0.008 | 0.0732 |
| 0.2862 | 0.0042 | 0.0422 | 0.6836 | 0.2582 | 0.0036 | 0.0124 | 0.0266 |
| 0.0018 | 0.0088 | 0.1532 | 0.8266 | 0.0012 | 0.0086 | 0.01 | 0.3022 |
| 0.089 | 0.0656 | 0.0094 | 0.8394 | 0.0782 | 0.0642 | 0.0088 | 0.0184 |
| 0.003 | 0.0416 | 0.162 | 0.7854 | 0.0024 | 0.0398 | 0.0104 | 0.2528 |

| | | | | | | | |
|---|---|---|---|---|---|---|---|
| 0.0108 | 0.0156 | 0.1198 | 0.8498 | 0.009 | 0.0142 | 0.007 | 0.3774 |
| 0.005 | 0.0126 | 0.1052 | 0.8728 | 0.0046 | 0.0102 | 0.0072 | 0.173 |
| 0.0132 | 0.0068 | 0.2104 | 0.7588 | 0.0096 | 0.0062 | 0.015 | 0.5256 |
| 0.183 | 0.05 | 0.3194 | 0.4826 | 0.1172 | 0.0124 | 0.0684 | 0.347 |
| 0.095 | 0.016 | 0.1072 | 0.7854 | 0.0888 | 0.014 | 0.0044 | 0.1752 |
| 0.0242 | 0.0058 | 0.0228 | 0.9424 | 0.0198 | 0.0052 | 0.0096 | 0.0756 |
| 0.1238 | 0.0088 | 0.0696 | 0.7976 | 0.1182 | 0.0086 | 0.0064 | 0.2012 |
| 0.013 | 0.0198 | 0.134 | 0.8172 | 0.0094 | 0.0194 | 0.0198 | 0.2466 |
| 0.0926 | 0.0094 | 0.0392 | 0.8564 | 0.0804 | 0.0084 | 0.016 | 0.0612 |
| 0.0762 | 0.0522 | 0.3412 | 0.5448 | 0.0202 | 0.0136 | 0.0798 | 0.2822 |
| 0.0024 | 0.0182 | 0.172 | 0.7924 | 0.002 | 0.0178 | 0.0154 | 0.357 |
| 0.003 | 0.0754 | 0.1768 | 0.7098 | 0.0018 | 0.0254 | 0.0866 | 0.1178 |
| 0.0386 | 0.0092 | 0.1402 | 0.7538 | 0.0208 | 0.0086 | 0.0764 | 0.087 |
| 0.0118 | 0.0148 | 0.0308 | 0.9212 | 0.0104 | 0.0144 | 0.0234 | 0.0302 |
| 0.005 | 0.0206 | 0.17 | 0.7804 | 0.004 | 0.022 | 0.0236 | 0.437 |
| 0.0996 | 0.0048 | 0.1868 | 0.7146 | 0.0896 | 0.0042 | 0.0046 | 0.4188 |
| 0.1912 | 0.0494 | 0.1168 | 0.6318 | 0.1742 | 0.0516 | 0.0256 | 0.2476 |
| 0.024 | 0.0318 | 0.2256 | 0.6692 | 0.0112 | 0.0168 | 0.077 | 0.21 |
| 0.0658 | 0.007 | 0.0156 | 0.8486 | 0.0678 | 0.0072 | 0.061 | 0.0242 |
| 0.1172 | 0.0026 | 0.1778 | 0.7064 | 0.109 | 0.0026 | 0.0042 | 0.5112 |
| 0.1446 | 0.008 | 0.0342 | 0.8136 | 0.1356 | 0.0074 | 0.009 | 0.0648 |
| 0.2318 | 0.013 | 0.3682 | 0.3518 | 0.165 | 0.0088 | 0.1058 | 0.2598 |
| 0.0974 | 0.0168 | 0.1618 | 0.593 | 0.0956 | 0.0318 | 0.118 | 0.4246 |
| 0.0428 | 0.238 | 0.3944 | 0.395 | 0.0112 | 0.1022 | 0.0976 | 0.362 |
| 0.0192 | 0.0188 | 0.0822 | 0.8442 | 0.0176 | 0.0212 | 0.0342 | 0.2422 |
| 0.02 | 0.006 | 0.0516 | 0.8434 | 0.0154 | 0.008 | 0.082 | 0.1322 |
| 0.1996 | 0.0044 | 0.1056 | 0.691 | 0.1914 | 0.0046 | 0.0074 | 0.1366 |
| 0.1552 | 0.0052 | 0.08 | 0.7398 | 0.1414 | 0.0054 | 0.033 | 0.222 |
| 0.0616 | 0.0272 | 0.022 | 0.6788 | 0.0576 | 0.0536 | 0.1878 | 0.0204 |
| 0.4198 | 0.0544 | 0.2134 | 0.349 | 0.3766 | 0.033 | 0.028 | 0.3518 |
| 0.0766 | 0.0212 | 0.042 | 0.8412 | 0.059 | 0.022 | 0.0364 | 0.0648 |
| 0.111 | 0.0138 | 0.075 | 0.8048 | 0.0976 | 0.013 | 0.0098 | 0.209 |
| 0.012 | 0.1596 | 0.1144 | 0.7242 | 0.0068 | 0.1146 | 0.0396 | 0.0602 |
| 0.1428 | 0.0048 | 0.139 | 0.6526 | 0.1414 | 0.0058 | 0.0614 | 0.2508 |
| 0.0194 | 0.0094 | 0.1948 | 0.7738 | 0.0166 | 0.0088 | 0.0058 | 0.5548 |
| 0.188 | 0.0094 | 0.1632 | 0.6468 | 0.1742 | 0.0092 | 0.0064 | 0.381 |
| 0.0102 | 0.0042 | 0.0132 | 0.8934 | 0.009 | 0.0048 | 0.0794 | 0.0582 |
| 0.0102 | 0.0474 | 0.2018 | 0.7064 | 0.008 | 0.0484 | 0.0354 | 0.4058 |
| 0.0036 | 0.0102 | 0.0386 | 0.9216 | 0.003 | 0.0112 | 0.026 | 0.1068 |
| 0.0564 | 0.1436 | 0.1918 | 0.662 | 0.0268 | 0.0712 | 0.0484 | 0.108 |
| 0.0068 | 0.0468 | 0.0674 | 0.8564 | 0.0048 | 0.0448 | 0.0268 | 0.1434 |
| 0.0968 | 0.0034 | 0.0736 | 0.8296 | 0.0874 | 0.003 | 0.0064 | 0.2578 |
| 0.0054 | 0.0034 | 0.1438 | 0.8408 | 0.0044 | 0.003 | 0.0082 | 0.3048 |
| 0.0022 | 0.0188 | 0.1888 | 0.7776 | 0.002 | 0.0192 | 0.0124 | 0.516 |

| | | | | | | | |
|---|---|---|---|---|---|---|---|
| 0.1014 | 0.05 | 0.0968 | 0.703 | 0.0814 | 0.0494 | 0.0692 | 0.1012 |
| 0.038 | 0.0282 | 0.2136 | 0.7034 | 0.0262 | 0.0236 | 0.033 | 0.4606 |
| 0.1064 | 0.0954 | 0.3822 | 0.4458 | 0.0352 | 0.0212 | 0.1158 | 0.3058 |
| 0.0072 | 0.0052 | 0.1028 | 0.876 | 0.0058 | 0.0052 | 0.0102 | 0.2792 |
| 0.1306 | 0.0062 | 0.1518 | 0.7064 | 0.1222 | 0.0066 | 0.0134 | 0.4144 |
| 0.0442 | 0.0052 | 0.0392 | 0.8032 | 0.0422 | 0.0078 | 0.1074 | 0.21 |
| 0.002 | 0.005 | 0.0182 | 0.9702 | 0.0012 | 0.0046 | 0.006 | 0.0756 |
| 0.002 | 0.0028 | 0.1728 | 0.8186 | 0.002 | 0.0024 | 0.0046 | 0.2726 |
| 0.0148 | 0.0024 | 0.0124 | 0.9702 | 0.0128 | 0.002 | 0.0028 | 0.0462 |
| 0.2886 | 0.007 | 0.1532 | 0.5636 | 0.2708 | 0.0066 | 0.0056 | 0.4046 |
| 0.3418 | 0.0278 | 0.0292 | 0.5908 | 0.3224 | 0.0294 | 0.0284 | 0.024 |
| 0.3326 | 0.0068 | 0.0804 | 0.568 | 0.3146 | 0.0064 | 0.0304 | 0.1082 |
| 0.1664 | 0.014 | 0.2202 | 0.5946 | 0.1158 | 0.0068 | 0.0624 | 0.1866 |
| 0.0084 | 0.0044 | 0.0344 | 0.9502 | 0.0062 | 0.0038 | 0.0052 | 0.1208 |
| 0.1272 | 0.1522 | 0.3316 | 0.4718 | 0.0498 | 0.0432 | 0.1032 | 0.1998 |
| 0.3676 | 0.0396 | 0.1376 | 0.4578 | 0.3436 | 0.0386 | 0.0222 | 0.3638 |
| 0.0068 | 0.0086 | 0.016 | 0.954 | 0.005 | 0.0072 | 0.0182 | 0.0122 |
| 0.0026 | 0.0086 | 0.1646 | 0.7518 | 0.0018 | 0.0062 | 0.076 | 0.1964 |
| 0.0146 | 0.0888 | 0.0966 | 0.7932 | 0.0088 | 0.0632 | 0.0386 | 0.118 |
| 0.0218 | 0.0048 | 0.1258 | 0.8416 | 0.0188 | 0.0044 | 0.0094 | 0.3854 |
| 0.0366 | 0.0074 | 0.0396 | 0.9164 | 0.0296 | 0.0068 | 0.0076 | 0.1062 |
| 0.0974 | 0.0526 | 0.2132 | 0.6502 | 0.0616 | 0.0344 | 0.0412 | 0.2186 |
| 0.05 | 0.0732 | 0.0124 | 0.751 | 0.0448 | 0.0926 | 0.0994 | 0.0178 |
| 0.1022 | 0.016 | 0.226 | 0.656 | 0.0362 | 0.0078 | 0.0742 | 0.1394 |
| 0.0472 | 0.0022 | 0.169 | 0.7742 | 0.0452 | 0.0022 | 0.0094 | 0.4554 |
| 0.0138 | 0.0054 | 0.031 | 0.8986 | 0.0112 | 0.006 | 0.053 | 0.1066 |
| 0.0082 | 0.0196 | 0.1214 | 0.836 | 0.0066 | 0.0202 | 0.016 | 0.2266 |
| 0.0034 | 0.014 | 0.1008 | 0.8414 | 0.0024 | 0.012 | 0.0436 | 0.1676 |
| 0.0318 | 0.0166 | 0.1328 | 0.7928 | 0.0272 | 0.0182 | 0.029 | 0.4214 |
| 0.004 | 0.0166 | 0.1596 | 0.8112 | 0.0034 | 0.0142 | 0.0118 | 0.3798 |
| 0.0056 | 0.0374 | 0.152 | 0.7772 | 0.0036 | 0.025 | 0.0424 | 0.1244 |
| 0.0364 | 0.005 | 0.0492 | 0.9002 | 0.0332 | 0.0048 | 0.0122 | 0.1314 |
| 0.0024 | 0.0032 | 0.2236 | 0.7216 | 0.0012 | 0.0026 | 0.0506 | 0.2974 |
| 0.0048 | 0.0156 | 0.123 | 0.8472 | 0.0036 | 0.015 | 0.011 | 0.3324 |
| 0.0036 | 0.0172 | 0.0484 | 0.8808 | 0.003 | 0.0206 | 0.0476 | 0.074 |
| 0.1572 | 0.007 | 0.0184 | 0.8138 | 0.15 | 0.0064 | 0.0116 | 0.0242 |
| 0.0146 | 0.0184 | 0.063 | 0.8614 | 0.0128 | 0.0222 | 0.0408 | 0.1504 |
| 0.0164 | 0.008 | 0.1848 | 0.7612 | 0.0126 | 0.0078 | 0.0336 | 0.4844 |
| 0.0492 | 0.0078 | 0.2166 | 0.7298 | 0.0374 | 0.0064 | 0.0102 | 0.534 |
| 0.0872 | 0.0802 | 0.2524 | 0.6038 | 0.037 | 0.0246 | 0.0822 | 0.2076 |
| 0.0186 | 0.0562 | 0.0844 | 0.7904 | 0.0162 | 0.064 | 0.045 | 0.2134 |
| 0.0366 | 0.0282 | 0.145 | 0.7828 | 0.0254 | 0.0218 | 0.0252 | 0.1784 |
| 0.0296 | 0.0138 | 0.0522 | 0.8708 | 0.0238 | 0.0136 | 0.0398 | 0.077 |
| 0.0176 | 0.0172 | 0.1574 | 0.801 | 0.0124 | 0.0166 | 0.0128 | 0.4582 |

| | | | | | | | |
|---|---|---|---|---|---|---|---|
| 0.1708 | 0.005 | 0.1256 | 0.6918 | 0.1582 | 0.005 | 0.0192 | 0.2968 |
| 0.009 | 0.0106 | 0.1104 | 0.8258 | 0.005 | 0.0076 | 0.051 | 0.0528 |
| 0.0098 | 0.006 | 0.2308 | 0.6846 | 0.0048 | 0.0032 | 0.0764 | 0.2374 |
| 0.0066 | 0.0028 | 0.0758 | 0.91 | 0.0054 | 0.0026 | 0.0064 | 0.2264 |
| 0.005 | 0.0176 | 0.17 | 0.7582 | 0.0024 | 0.005 | 0.0644 | 0.1366 |
| 0.0168 | 0.0082 | 0.015 | 0.9072 | 0.0124 | 0.008 | 0.0574 | 0.0156 |
| 0.1042 | 0.0476 | 0.3538 | 0.495 | 0.0312 | 0.0094 | 0.1108 | 0.3418 |
| 0.0216 | 0.05 | 0.2168 | 0.7168 | 0.013 | 0.0406 | 0.0132 | 0.3768 |
| 0.2826 | 0.0582 | 0.2118 | 0.4366 | 0.251 | 0.0516 | 0.0488 | 0.2304 |
| 0.41 | 0.0156 | 0.1256 | 0.459 | 0.3904 | 0.0146 | 0.0104 | 0.3526 |
| 0.0034 | 0.0078 | 0.1506 | 0.8326 | 0.0024 | 0.0076 | 0.007 | 0.3642 |
| 0.009 | 0.0118 | 0.175 | 0.7896 | 0.007 | 0.011 | 0.0176 | 0.3934 |
| 0.003 | 0.0136 | 0.1928 | 0.7862 | 0.0022 | 0.0118 | 0.0068 | 0.4312 |
| 0.0576 | 0.1436 | 0.188 | 0.5428 | 0.0438 | 0.1512 | 0.0742 | 0.2898 |
| 0.1282 | 0.0322 | 0.0758 | 0.7618 | 0.1186 | 0.0338 | 0.01 | 0.1956 |
| 0.0054 | 0.0072 | 0.1326 | 0.8452 | 0.004 | 0.0064 | 0.0116 | 0.298 |
| 0.0044 | 0.018 | 0.0998 | 0.8388 | 0.0038 | 0.0206 | 0.0376 | 0.1644 |
| 0.025 | 0.013 | 0.1334 | 0.8168 | 0.0136 | 0.008 | 0.0282 | 0.2828 |
| 0.2272 | 0.0058 | 0.1384 | 0.638 | 0.213 | 0.0052 | 0.0054 | 0.2704 |
| 0.2594 | 0.031 | 0.0104 | 0.5716 | 0.25 | 0.049 | 0.1188 | 0.0178 |
| 0.011 | 0.0422 | 0.0466 | 0.846 | 0.0084 | 0.0484 | 0.0508 | 0.1468 |
| 0.0256 | 0.112 | 0.0916 | 0.7674 | 0.0182 | 0.1048 | 0.0184 | 0.1266 |
| 0.1112 | 0.0154 | 0.1572 | 0.7168 | 0.1032 | 0.0156 | 0.0072 | 0.481 |
| 0.1812 | 0.0238 | 0.137 | 0.6604 | 0.1464 | 0.0126 | 0.044 | 0.1698 |
| 0.0044 | 0.1242 | 0.0494 | 0.7652 | 0.0032 | 0.1262 | 0.0558 | 0.031 |
| 0.0048 | 0.007 | 0.0174 | 0.9394 | 0.0038 | 0.0074 | 0.0324 | 0.0188 |
| 0.0046 | 0.0244 | 0.1808 | 0.783 | 0.0032 | 0.024 | 0.0088 | 0.5302 |
| 0.0796 | 0.0134 | 0.1546 | 0.7414 | 0.0748 | 0.013 | 0.0166 | 0.245 |
| 0.0424 | 0.0274 | 0.288 | 0.6198 | 0.0216 | 0.0258 | 0.0452 | 0.48 |
| 0.0814 | 0.2214 | 0.258 | 0.4738 | 0.0496 | 0.19 | 0.0286 | 0.4646 |
| 0.0352 | 0.0432 | 0.2884 | 0.6402 | 0.0136 | 0.0148 | 0.0432 | 0.4762 |
| 0.0438 | 0.0158 | 0.1978 | 0.7438 | 0.0368 | 0.013 | 0.0086 | 0.537 |
| 0.0068 | 0.4744 | 0.101 | 0.3974 | 0.0054 | 0.474 | 0.0224 | 0.2426 |
| 0.0038 | 0.0076 | 0.0102 | 0.9272 | 0.0026 | 0.0082 | 0.052 | 0.0224 |
| 0.0032 | 0.289 | 0.2014 | 0.4996 | 0.0022 | 0.2462 | 0.0506 | 0.2374 |
| 0.0346 | 0.0172 | 0.1468 | 0.7898 | 0.0296 | 0.016 | 0.0182 | 0.2946 |
| 0.0068 | 0.0232 | 0.2946 | 0.612 | 0.0028 | 0.0072 | 0.0838 | 0.3168 |
| 0.0226 | 0.0776 | 0.2132 | 0.7004 | 0.0148 | 0.0562 | 0.015 | 0.5296 |
| 0.0054 | 0.0474 | 0.102 | 0.834 | 0.0038 | 0.0442 | 0.0154 | 0.196 |
| 0.009 | 0.0084 | 0.195 | 0.7752 | 0.0064 | 0.0074 | 0.0162 | 0.4782 |
| 0.952 | 0.0058 | 0.0184 | 0.067 | 0.9016 | 0.0064 | 0.0066 | 0.0186 |
| 0.0066 | 0.0044 | 0.0296 | 0.9554 | 0.0052 | 0.004 | 0.0058 | 0.0738 |
| 0.0388 | 0.0214 | 0.3036 | 0.5712 | 0.008 | 0.0048 | 0.1122 | 0.167 |
| 0.0088 | 0.0582 | 0.2478 | 0.6466 | 0.0038 | 0.011 | 0.0912 | 0.2008 |

| | | | | | | | |
|---|---|---|---|---|---|---|---|
| 0.2422 | 0.0084 | 0.1126 | 0.6324 | 0.2322 | 0.0084 | 0.0148 | 0.346 |
| 0.0064 | 0.0136 | 0.0236 | 0.943 | 0.0048 | 0.0138 | 0.015 | 0.0352 |
| 0.133 | 0.0594 | 0.297 | 0.5328 | 0.0508 | 0.013 | 0.1064 | 0.1602 |
| 0.0304 | 0.0072 | 0.0922 | 0.8602 | 0.0254 | 0.006 | 0.0162 | 0.141 |
| 0.005 | 0.0112 | 0.075 | 0.884 | 0.0042 | 0.0124 | 0.0248 | 0.169 |
| 0.0038 | 0.0858 | 0.0322 | 0.8714 | 0.0032 | 0.0822 | 0.0106 | 0.1182 |
| 0.0092 | 0.0222 | 0.021 | 0.945 | 0.0066 | 0.019 | 0.0082 | 0.0142 |
| 0.3552 | 0.026 | 0.1368 | 0.4666 | 0.3366 | 0.0282 | 0.0312 | 0.364 |
| 0.012 | 0.0208 | 0.0554 | 0.8906 | 0.0092 | 0.0168 | 0.0286 | 0.0556 |
| 0.0092 | 0.1508 | 0.0168 | 0.807 | 0.0074 | 0.1496 | 0.0188 | 0.0368 |
| 0.0076 | 0.0212 | 0.1866 | 0.7138 | 0.0056 | 0.026 | 0.0674 | 0.5232 |
| 0.0082 | 0.141 | 0.3312 | 0.5226 | 0.0042 | 0.058 | 0.084 | 0.392 |
| 0.0024 | 0.0066 | 0.0238 | 0.9576 | 0.002 | 0.0064 | 0.0108 | 0.0674 |
| 0.1524 | 0.303 | 0.0464 | 0.4806 | 0.1444 | 0.3094 | 0.019 | 0.113 |
| 0.0212 | 0.0274 | 0.075 | 0.8644 | 0.0164 | 0.027 | 0.0174 | 0.298 |
| 0.0132 | 0.0056 | 0.1458 | 0.7776 | 0.0066 | 0.004 | 0.0664 | 0.098 |
| 0.0088 | 0.0822 | 0.0252 | 0.7588 | 0.007 | 0.1062 | 0.1028 | 0.024 |
| 0.0064 | 0.039 | 0.1004 | 0.8224 | 0.0046 | 0.0374 | 0.035 | 0.18 |
| 0.0072 | 0.0318 | 0.0322 | 0.8682 | 0.006 | 0.039 | 0.0546 | 0.0416 |
| 0.004 | 0.0284 | 0.0394 | 0.8572 | 0.0032 | 0.0364 | 0.0644 | 0.0468 |
| 0.244 | 0.0666 | 0.114 | 0.5772 | 0.2268 | 0.0656 | 0.0162 | 0.1616 |
| 0.315 | 0.0066 | 0.1486 | 0.5402 | 0.2964 | 0.0066 | 0.0082 | 0.3698 |
| 0.1234 | 0.0152 | 0.011 | 0.8048 | 0.1192 | 0.018 | 0.047 | 0.0412 |
| 0.0066 | 0.0224 | 0.2642 | 0.645 | 0.003 | 0.0076 | 0.0806 | 0.2296 |
| 0.4292 | 0.0108 | 0.101 | 0.4602 | 0.4104 | 0.0122 | 0.0162 | 0.1614 |
| 0.0086 | 0.0186 | 0.2064 | 0.7538 | 0.0062 | 0.0152 | 0.0184 | 0.5106 |
| 0.0308 | 0.0766 | 0.216 | 0.674 | 0.0132 | 0.0336 | 0.0632 | 0.176 |
| 0.1064 | 0.0162 | 0.1052 | 0.77 | 0.0972 | 0.015 | 0.0126 | 0.2074 |
| 0.0042 | 0.0048 | 0.1496 | 0.8384 | 0.0032 | 0.0042 | 0.0044 | 0.3056 |
| 0.0022 | 0.0032 | 0.0714 | 0.9192 | 0.0018 | 0.0028 | 0.0048 | 0.1942 |
| 0.0102 | 0.0086 | 0.2568 | 0.6838 | 0.005 | 0.0046 | 0.0492 | 0.4764 |
| 0.4226 | 0.1908 | 0.0112 | 0.3696 | 0.4008 | 0.1978 | 0.0206 | 0.0126 |
| 0.0062 | 0.4574 | 0.2034 | 0.3264 | 0.0032 | 0.3724 | 0.0946 | 0.1504 |
| 0.0272 | 0.1896 | 0.129 | 0.622 | 0.0218 | 0.1968 | 0.0298 | 0.1732 |
| 0.3784 | 0.032 | 0.0248 | 0.5578 | 0.3616 | 0.0358 | 0.02 | 0.0858 |
| 0.1812 | 0.0128 | 0.061 | 0.7414 | 0.168 | 0.011 | 0.0188 | 0.096 |
| 0.0148 | 0.0054 | 0.046 | 0.9316 | 0.0118 | 0.0052 | 0.0048 | 0.0682 |
| 0.0038 | 0.0176 | 0.0616 | 0.9086 | 0.0028 | 0.0172 | 0.0094 | 0.2132 |
| 0.3394 | 0.0198 | 0.1234 | 0.4852 | 0.3084 | 0.02 | 0.0634 | 0.0872 |
| 0.1262 | 0.0142 | 0.0478 | 0.8032 | 0.113 | 0.0128 | 0.0232 | 0.0502 |
| 0.034 | 0.0214 | 0.078 | 0.8468 | 0.0288 | 0.019 | 0.0274 | 0.1324 |
| 0.0026 | 0.014 | 0.1332 | 0.8392 | 0.002 | 0.0124 | 0.0132 | 0.2216 |
| 0.0038 | 0.0044 | 0.013 | 0.9754 | 0.003 | 0.004 | 0.005 | 0.055 |
| 0.0814 | 0.0462 | 0.123 | 0.7552 | 0.0702 | 0.0416 | 0.0098 | 0.3796 |

| | | | | | | | |
|---|---|---|---|---|---|---|---|
| 0.092 | 0.0044 | 0.2258 | 0.6772 | 0.0694 | 0.0034 | 0.0236 | 0.5094 |
| 0.0022 | 0.016 | 0.258 | 0.6796 | 0.0012 | 0.0076 | 0.0536 | 0.459 |
| 0.1454 | 0.0728 | 0.119 | 0.6264 | 0.138 | 0.0804 | 0.036 | 0.3054 |
| 0.3006 | 0.011 | 0.1302 | 0.5546 | 0.2882 | 0.0106 | 0.0164 | 0.264 |
| 0.1702 | 0.2384 | 0.3284 | 0.346 | 0.1102 | 0.125 | 0.0902 | 0.276 |
| 0.0088 | 0.014 | 0.0366 | 0.9176 | 0.0062 | 0.0142 | 0.025 | 0.0424 |
| 0.2232 | 0.0054 | 0.1258 | 0.6412 | 0.2154 | 0.0054 | 0.0118 | 0.2546 |
| 0.492 | 0.0324 | 0.1064 | 0.3796 | 0.4686 | 0.033 | 0.0122 | 0.2264 |
| 0.2968 | 0.027 | 0.0958 | 0.539 | 0.2856 | 0.0338 | 0.0458 | 0.2364 |
| 0.4364 | 0.048 | 0.0134 | 0.4822 | 0.4138 | 0.0618 | 0.0288 | 0.024 |
| 0.2792 | 0.0056 | 0.1396 | 0.5774 | 0.2654 | 0.0056 | 0.0118 | 0.3606 |
| 0.2168 | 0.0044 | 0.0118 | 0.7752 | 0.2038 | 0.0036 | 0.0056 | 0.0196 |
| 0.2762 | 0.5426 | 0.0488 | 0.15 | 0.2626 | 0.5318 | 0.007 | 0.072 |
| 0.0054 | 0.0098 | 0.103 | 0.8658 | 0.0044 | 0.0098 | 0.0168 | 0.2776 |
| 0.003 | 0.1824 | 0.0986 | 0.7076 | 0.0026 | 0.1812 | 0.0102 | 0.2344 |
| 0.0066 | 0.0086 | 0.1486 | 0.8286 | 0.0046 | 0.0078 | 0.0102 | 0.4336 |
| 0.0026 | 0.7846 | 0.0138 | 0.0258 | 0.0022 | 0.8074 | 0.151 | 0.0262 |
| 0.0994 | 0.8738 | 0.0136 | 0.0162 | 0.0954 | 0.864 | 0.011 | 0.0174 |
| 0.0322 | 0.9442 | 0.0122 | 0.008 | 0.0296 | 0.935 | 0.0152 | 0.0066 |
| 0.0382 | 0.004 | 0.1084 | 0.824 | 0.0194 | 0.0028 | 0.0456 | 0.0708 |
| 0.0122 | 0.9634 | 0.0062 | 0.0056 | 0.0128 | 0.9608 | 0.0146 | 0.0086 |
| 0.004 | 0.9016 | 0.0272 | 0.0812 | 0.0022 | 0.877 | 0.0118 | 0.0618 |
| 0.0508 | 0.923 | 0.0258 | 0.0214 | 0.0436 | 0.8958 | 0.0128 | 0.0296 |
| 0.0456 | 0.8774 | 0.0322 | 0.0306 | 0.0378 | 0.8644 | 0.0352 | 0.0326 |
| 0.0114 | 0.8894 | 0.1074 | 0.0302 | 0.0052 | 0.8246 | 0.0328 | 0.0642 |
| 0.012 | 0.9518 | 0.0206 | 0.0228 | 0.0098 | 0.9346 | 0.0128 | 0.0318 |
| 0.002 | 0.886 | 0.007 | 0.0058 | 0.0014 | 0.9086 | 0.0766 | 0.008 |
| 0.0238 | 0.7578 | 0.0058 | 0.0052 | 0.0264 | 0.7928 | 0.1694 | 0.0056 |
| 0.0018 | 0.982 | 0.035 | 0.0078 | 0.001 | 0.9406 | 0.0156 | 0.0266 |
| 0.0024 | 0.0272 | 0.1286 | 0.8132 | 0.002 | 0.0254 | 0.0306 | 0.2598 |
| 0.0026 | 0.9356 | 0.0198 | 0.0318 | 0.002 | 0.9182 | 0.0286 | 0.0194 |
| 0.0244 | 0.9366 | 0.0214 | 0.0202 | 0.0212 | 0.9168 | 0.0204 | 0.0282 |
| 0.0034 | 0.9204 | 0.0224 | 0.0754 | 0.0022 | 0.8846 | 0.015 | 0.0298 |
| 0.01 | 0.8782 | 0.0282 | 0.0192 | 0.008 | 0.8746 | 0.0698 | 0.0238 |
| 0.0068 | 0.942 | 0.0162 | 0.019 | 0.0054 | 0.9302 | 0.0292 | 0.0192 |
| 0.0104 | 0.265 | 0.0186 | 0.434 | 0.0098 | 0.3086 | 0.2286 | 0.0302 |
| 0.0078 | 0.977 | 0.0078 | 0.0096 | 0.008 | 0.9676 | 0.007 | 0.0112 |
| 0.0174 | 0.9368 | 0.0114 | 0.0094 | 0.0172 | 0.9306 | 0.0312 | 0.0182 |
| 0.0048 | 0.8386 | 0.0072 | 0.01 | 0.0048 | 0.86 | 0.1178 | 0.0108 |
| 0.0098 | 0.9728 | 0.0066 | 0.0028 | 0.0098 | 0.9684 | 0.0118 | 0.0066 |
| 0.0688 | 0.8984 | 0.0062 | 0.0036 | 0.0708 | 0.9018 | 0.0174 | 0.0064 |
| 0.0164 | 0.9688 | 0.01 | 0.0108 | 0.0156 | 0.9562 | 0.0074 | 0.011 |
| 0.002 | 0.9574 | 0.0118 | 0.0156 | 0.0014 | 0.949 | 0.0222 | 0.0188 |
| 0.0196 | 0.9274 | 0.0378 | 0.0416 | 0.0142 | 0.8888 | 0.0176 | 0.0126 |

| | | | | | | | |
|---|---|---|---|---|---|---|---|
| 0.049 | 0.9052 | 0.1298 | 0.0068 | 0.0132 | 0.8088 | 0.0412 | 0.0724 |
| 0.0408 | 0.941 | 0.0078 | 0.0076 | 0.0434 | 0.9326 | 0.0088 | 0.0126 |
| 0.004 | 0.9176 | 0.1016 | 0.0032 | 0.0018 | 0.8442 | 0.0494 | 0.052 |
| 0.0024 | 0.7576 | 0.0032 | 0.0032 | 0.0024 | 0.8022 | 0.1886 | 0.0034 |
| 0.0192 | 0.8822 | 0.0104 | 0.0276 | 0.0172 | 0.8834 | 0.0608 | 0.012 |
| 0.0392 | 0.887 | 0.0426 | 0.0402 | 0.0292 | 0.861 | 0.0262 | 0.0394 |
| 0.0032 | 0.9268 | 0.007 | 0.008 | 0.0032 | 0.9386 | 0.0432 | 0.009 |
| 0.0128 | 0.9776 | 0.135 | 0.0024 | 0.0032 | 0.8212 | 0.0384 | 0.0734 |
| 0.0094 | 0.8304 | 0.0274 | 0.0922 | 0.0072 | 0.8302 | 0.0428 | 0.0398 |
| 0.0034 | 0.8546 | 0.0282 | 0.1154 | 0.003 | 0.8364 | 0.0172 | 0.0408 |
| 0.0096 | 0.8768 | 0.1552 | 0.0106 | 0.0034 | 0.7828 | 0.0482 | 0.086 |
| 0.007 | 0.9046 | 0.2152 | 0.0036 | 0.0024 | 0.7238 | 0.0548 | 0.1154 |
| 0.01 | 0.9256 | 0.0212 | 0.0298 | 0.009 | 0.9108 | 0.0294 | 0.0256 |
| 0.0052 | 0.9444 | 0.0098 | 0.0146 | 0.0042 | 0.9412 | 0.0298 | 0.0138 |
| 0.0042 | 0.8046 | 0.0052 | 0.009 | 0.0032 | 0.8402 | 0.1424 | 0.0072 |
| 0.0082 | 0.0034 | 0.0228 | 0.9604 | 0.0064 | 0.0032 | 0.007 | 0.0856 |
| 0.004 | 0.976 | 0.0084 | 0.0064 | 0.0036 | 0.9698 | 0.0116 | 0.0144 |
| 0.0096 | 0.0116 | 0.1788 | 0.7902 | 0.0088 | 0.0104 | 0.0116 | 0.4482 |
| 0.0072 | 0.8192 | 0.1992 | 0.0192 | 0.0032 | 0.718 | 0.0598 | 0.1034 |
| 0.0032 | 0.7604 | 0.008 | 0.0086 | 0.003 | 0.7972 | 0.1836 | 0.0126 |
| 0.0648 | 0.9172 | 0.0078 | 0.0076 | 0.0696 | 0.9064 | 0.009 | 0.0074 |
| 0.0048 | 0.9866 | 0.0042 | 0.0022 | 0.006 | 0.9818 | 0.0058 | 0.005 |
| 0.121 | 0.8102 | 0.1304 | 0.0094 | 0.08 | 0.7342 | 0.046 | 0.0682 |
| 0.013 | 0.967 | 0.1286 | 0.003 | 0.003 | 0.8282 | 0.0374 | 0.0674 |
| 0.005 | 0.9584 | 0.0034 | 0.0022 | 0.0054 | 0.963 | 0.0258 | 0.0032 |
| 0.007 | 0.9332 | 0.036 | 0.0242 | 0.0056 | 0.9086 | 0.0254 | 0.0366 |
| 0.072 | 0.8874 | 0.0144 | 0.0266 | 0.069 | 0.8746 | 0.0152 | 0.0182 |
| 0.005 | 0.9214 | 0.0266 | 0.0616 | 0.0034 | 0.8886 | 0.02 | 0.036 |
| 0.0462 | 0.8318 | 0.0236 | 0.058 | 0.0446 | 0.8266 | 0.0472 | 0.0364 |
| 0.0046 | 0.9684 | 0.0182 | 0.0242 | 0.0036 | 0.9396 | 0.0146 | 0.0268 |
| 0.004 | 0.9726 | 0.0682 | 0.0108 | 0.0024 | 0.895 | 0.024 | 0.0464 |
| 0.0122 | 0.0044 | 0.0174 | 0.9548 | 0.0114 | 0.004 | 0.0128 | 0.0364 |
| 0.0054 | 0.9838 | 0.0056 | 0.005 | 0.006 | 0.9772 | 0.006 | 0.0078 |
| 0.0194 | 0.9296 | 0.0156 | 0.0126 | 0.0184 | 0.9222 | 0.031 | 0.0214 |
| 0.003 | 0.9332 | 0.0052 | 0.0086 | 0.0022 | 0.9398 | 0.0442 | 0.0052 |
| 0.0082 | 0.9794 | 0.0744 | 0.0034 | 0.0044 | 0.891 | 0.0268 | 0.0378 |
| 0.0056 | 0.9524 | 0.0226 | 0.0322 | 0.0042 | 0.9248 | 0.016 | 0.0282 |
| 0.0326 | 0.935 | 0.015 | 0.022 | 0.0308 | 0.9184 | 0.0136 | 0.0172 |
| 0.0026 | 0.9274 | 0.0066 | 0.005 | 0.0022 | 0.937 | 0.0492 | 0.0064 |
| 0.016 | 0.8982 | 0.015 | 0.0068 | 0.0172 | 0.901 | 0.06 | 0.0146 |
| 0.0436 | 0.924 | 0.0098 | 0.0066 | 0.0444 | 0.921 | 0.0182 | 0.0118 |
| 0.0056 | 0.969 | 0.0062 | 0.0068 | 0.0054 | 0.9674 | 0.0144 | 0.0102 |
| 0.0106 | 0.8162 | 0.1576 | 0.0218 | 0.006 | 0.7598 | 0.0544 | 0.0824 |
| 0.0166 | 0.9648 | 0.0064 | 0.0036 | 0.0186 | 0.9598 | 0.0114 | 0.007 |

| | | | | | | | |
|---|---|---|---|---|---|---|---|
| 0.0036 | 0.988 | 0.0056 | 0.0048 | 0.0042 | 0.98 | 0.006 | 0.0068 |
| 0.0076 | 0.8836 | 0.1244 | 0.0256 | 0.0046 | 0.8126 | 0.0328 | 0.0754 |
| 0.0174 | 0.89 | 0.0072 | 0.0108 | 0.018 | 0.8984 | 0.0654 | 0.007 |
| 0.012 | 0.9394 | 0.0132 | 0.0116 | 0.011 | 0.9348 | 0.0294 | 0.0164 |
| 0.0736 | 0.892 | 0.0086 | 0.0056 | 0.0756 | 0.8906 | 0.0194 | 0.0074 |
| 0.0042 | 0.9288 | 0.1608 | 0.0046 | 0.0012 | 0.7876 | 0.0458 | 0.084 |
| 0.0282 | 0.9316 | 0.0302 | 0.0362 | 0.0232 | 0.8998 | 0.0106 | 0.0492 |
| 0.0076 | 0.9024 | 0.0252 | 0.0492 | 0.006 | 0.8892 | 0.0312 | 0.0344 |
| 0.0028 | 0.9732 | 0.0114 | 0.0212 | 0.0022 | 0.956 | 0.0094 | 0.0172 |
| 0.0996 | 0.108 | 0.0304 | 0.7514 | 0.0886 | 0.1152 | 0.014 | 0.0508 |
| 0.0132 | 0.9546 | 0.007 | 0.0094 | 0.012 | 0.9512 | 0.02 | 0.0068 |
| 0.0058 | 0.9238 | 0.0312 | 0.02 | 0.0048 | 0.9092 | 0.0342 | 0.0198 |
| 0.0834 | 0.8554 | 0.0556 | 0.0408 | 0.0566 | 0.8304 | 0.0166 | 0.0266 |
| 0.0508 | 0.8772 | 0.0186 | 0.0194 | 0.052 | 0.8724 | 0.038 | 0.0192 |
| 0.0032 | 0.9232 | 0.014 | 0.022 | 0.0028 | 0.9194 | 0.0416 | 0.0166 |
| 0.0032 | 0.4572 | 0.1022 | 0.3432 | 0.003 | 0.4716 | 0.0796 | 0.2736 |
| 0.0626 | 0.8988 | 0.0086 | 0.005 | 0.0638 | 0.8988 | 0.0236 | 0.0068 |
| 0.0028 | 0.943 | 0.0174 | 0.0482 | 0.0022 | 0.917 | 0.016 | 0.0294 |
| 0.0048 | 0.9558 | 0.0318 | 0.026 | 0.0034 | 0.919 | 0.0198 | 0.0306 |
| 0.0032 | 0.8248 | 0.0156 | 0.029 | 0.0026 | 0.8384 | 0.1146 | 0.0216 |
| 0.0036 | 0.6998 | 0.007 | 0.0056 | 0.0036 | 0.754 | 0.2302 | 0.007 |
| 0.003 | 0.9122 | 0.015 | 0.0182 | 0.0024 | 0.9076 | 0.0572 | 0.0174 |
| 0.0736 | 0.841 | 0.0072 | 0.0074 | 0.0766 | 0.8534 | 0.0552 | 0.007 |
| 0.0112 | 0.9278 | 0.0206 | 0.012 | 0.0096 | 0.9172 | 0.0402 | 0.0212 |
| 0.003 | 0.982 | 0.0088 | 0.0066 | 0.0026 | 0.9722 | 0.0094 | 0.013 |
| 0.0116 | 0.9114 | 0.0476 | 0.0448 | 0.007 | 0.879 | 0.0218 | 0.0516 |
| 0.0738 | 0.8538 | 0.0232 | 0.067 | 0.0684 | 0.8236 | 0.018 | 0.0184 |
| 0.01 | 0.9038 | 0.0208 | 0.06 | 0.0078 | 0.8838 | 0.0278 | 0.0192 |
| 0.0036 | 0.9808 | 0.0118 | 0.0174 | 0.003 | 0.9588 | 0.0088 | 0.0166 |
| 0.0288 | 0.8464 | 0.0176 | 0.0138 | 0.0318 | 0.8612 | 0.0752 | 0.0132 |
| 0.0216 | 0.9582 | 0.1076 | 0.0046 | 0.0062 | 0.8474 | 0.034 | 0.0632 |
| 0.0182 | 0.9584 | 0.0352 | 0.0064 | 0.0136 | 0.9268 | 0.0184 | 0.0184 |
| 0.003 | 0.9588 | 0.0054 | 0.0032 | 0.0034 | 0.9636 | 0.0242 | 0.0056 |
| 0.0128 | 0.6678 | 0.029 | 0.2636 | 0.01 | 0.6638 | 0.0336 | 0.048 |
| 0.0334 | 0.9136 | 0.1332 | 0.0098 | 0.0074 | 0.8096 | 0.0394 | 0.058 |
| 0.0078 | 0.8888 | 0.0396 | 0.0442 | 0.006 | 0.874 | 0.0362 | 0.0404 |
| 0.198 | 0.579 | 0.0222 | 0.062 | 0.1904 | 0.6024 | 0.1224 | 0.0418 |
| 0.0252 | 0.938 | 0.005 | 0.006 | 0.023 | 0.9432 | 0.023 | 0.0036 |
| 0.0056 | 0.9598 | 0.0112 | 0.0234 | 0.0048 | 0.9492 | 0.0116 | 0.0144 |
| 0.0036 | 0.9754 | 0.016 | 0.0136 | 0.003 | 0.9536 | 0.014 | 0.0146 |
| 0.0068 | 0.8324 | 0.0172 | 0.0498 | 0.006 | 0.8414 | 0.0856 | 0.0226 |
| 0.0056 | 0.4906 | 0.0974 | 0.408 | 0.0042 | 0.481 | 0.0088 | 0.2504 |
| 0.002 | 0.6444 | 0.1778 | 0.1824 | 0.0016 | 0.5696 | 0.069 | 0.1382 |
| 0.0104 | 0.9216 | 0.019 | 0.0634 | 0.0078 | 0.8962 | 0.0136 | 0.0292 |

| | | | | | | | |
|---|---|---|---|---|---|---|---|
| 0.013 | 0.9596 | 0.0048 | 0.0064 | 0.0134 | 0.9588 | 0.0166 | 0.0066 |
| 0.0268 | 0.8598 | 0.0374 | 0.0888 | 0.0216 | 0.8396 | 0.0126 | 0.0682 |
| 0.0252 | 0.8768 | 0.0078 | 0.0122 | 0.0272 | 0.8904 | 0.0624 | 0.0104 |
| 0.0152 | 0.5742 | 0.0864 | 0.1464 | 0.01 | 0.5834 | 0.1742 | 0.075 |
| 0.146 | 0.601 | 0.0184 | 0.0678 | 0.1386 | 0.624 | 0.1516 | 0.022 |
| 0.0148 | 0.9214 | 0.0142 | 0.0118 | 0.0132 | 0.9196 | 0.0414 | 0.02 |
| 0.0298 | 0.024 | 0.299 | 0.599 | 0.0128 | 0.0086 | 0.0802 | 0.3702 |
| 0.0032 | 0.9142 | 0.042 | 0.0088 | 0.0024 | 0.8896 | 0.0574 | 0.0366 |
| 0.024 | 0.9378 | 0.0126 | 0.0338 | 0.0232 | 0.9194 | 0.011 | 0.0238 |
| 0.002 | 0.978 | 0.0026 | 0.0022 | 0.0016 | 0.9828 | 0.0108 | 0.0026 |
| 0.0548 | 0.7568 | 0.0242 | 0.0386 | 0.0506 | 0.7672 | 0.1202 | 0.035 |
| 0.0078 | 0.729 | 0.0358 | 0.0232 | 0.006 | 0.7432 | 0.1922 | 0.0346 |
| 0.0048 | 0.8868 | 0.0234 | 0.082 | 0.0036 | 0.8708 | 0.02 | 0.0444 |
| 0.0022 | 0.9202 | 0.0072 | 0.0054 | 0.0022 | 0.9284 | 0.0572 | 0.0062 |
| 0.0046 | 0.973 | 0.0154 | 0.006 | 0.0042 | 0.958 | 0.0168 | 0.0174 |
| 0.0034 | 0.7978 | 0.0368 | 0.1174 | 0.0028 | 0.7966 | 0.0464 | 0.0564 |
| 0.0028 | 0.9024 | 0.0094 | 0.1074 | 0.002 | 0.8768 | 0.0046 | 0.015 |
| 0.0784 | 0.8854 | 0.0114 | 0.0232 | 0.0684 | 0.8788 | 0.0184 | 0.0106 |
| 0.0044 | 0.0242 | 0.0758 | 0.8674 | 0.0038 | 0.027 | 0.026 | 0.1702 |
| 0.4134 | 0.5678 | 0.0032 | 0.0036 | 0.4022 | 0.5824 | 0.0086 | 0.0032 |
| 0.0176 | 0.1546 | 0.024 | 0.7756 | 0.0142 | 0.1594 | 0.0266 | 0.0302 |
| 0.0154 | 0.4962 | 0.1062 | 0.3694 | 0.0102 | 0.4808 | 0.0332 | 0.208 |
| 0.024 | 0.4216 | 0.1368 | 0.3928 | 0.016 | 0.4114 | 0.0436 | 0.2096 |
| 0.101 | 0.0142 | 0.0132 | 0.8708 | 0.0918 | 0.0142 | 0.01 | 0.0794 |
| 0.004 | 0.0126 | 0.0322 | 0.9288 | 0.003 | 0.012 | 0.0242 | 0.051 |
| 0.0198 | 0.0094 | 0.1144 | 0.8542 | 0.0158 | 0.0082 | 0.0074 | 0.2894 |
| 0.0032 | 0.0104 | 0.1686 | 0.8052 | 0.0022 | 0.0088 | 0.015 | 0.4944 |
| 0.2276 | 0.0712 | 0.1366 | 0.5282 | 0.2166 | 0.081 | 0.0376 | 0.2546 |
| 0.0134 | 0.006 | 0.1886 | 0.7516 | 0.0048 | 0.003 | 0.052 | 0.284 |
| 0.0092 | 0.0296 | 0.255 | 0.6186 | 0.0032 | 0.0088 | 0.114 | 0.2132 |
| 0.1676 | 0.0182 | 0.0902 | 0.715 | 0.1586 | 0.02 | 0.016 | 0.1766 |
| 0.0072 | 0.0136 | 0.031 | 0.934 | 0.0056 | 0.0128 | 0.0162 | 0.1066 |
| 0.005 | 0.9594 | 0.0204 | 0.023 | 0.0038 | 0.9414 | 0.0118 | 0.0278 |
| 0.012 | 0.9318 | 0.0182 | 0.041 | 0.0102 | 0.9078 | 0.0226 | 0.0266 |
| 0.005 | 0.336 | 0.1308 | 0.529 | 0.0038 | 0.3294 | 0.007 | 0.2916 |
| 0.2212 | 0.3912 | 0.087 | 0.3034 | 0.2082 | 0.3896 | 0.0116 | 0.2388 |
| 0.0534 | 0.0182 | 0.0258 | 0.8828 | 0.0462 | 0.0184 | 0.0274 | 0.0648 |
| 0.0356 | 0.0508 | 0.0432 | 0.8514 | 0.0294 | 0.049 | 0.0264 | 0.0832 |
| 0.009 | 0.0102 | 0.186 | 0.751 | 0.0048 | 0.0046 | 0.0534 | 0.2762 |
| 0.0064 | 0.0132 | 0.1924 | 0.7528 | 0.0046 | 0.0128 | 0.0374 | 0.4672 |
| 0.2994 | 0.0674 | 0.0714 | 0.5098 | 0.283 | 0.0832 | 0.0526 | 0.1438 |
| 0.0362 | 0.0766 | 0.0752 | 0.8064 | 0.0286 | 0.0768 | 0.0134 | 0.2466 |
| 0.1126 | 0.019 | 0.207 | 0.6752 | 0.0922 | 0.0146 | 0.0108 | 0.445 |
| 0.0034 | 0.0046 | 0.0274 | 0.1332 | 0.0068 | 0.148 | 0.6848 | 0.0624 |

| | | | | | | | |
|---|---|---|---|---|---|---|---|
| 0.341 | 0.4688 | 0.0706 | 0.141 | 0.3148 | 0.4556 | 0.0182 | 0.1186 |
| 0.4658 | 0.0118 | 0.2622 | 0.2278 | 0.4052 | 0.0072 | 0.0978 | 0.1646 |
| 0.6396 | 0.032 | 0.0694 | 0.2118 | 0.6042 | 0.0484 | 0.0666 | 0.0832 |
| 0.033 | 0.705 | 0.2434 | 0.078 | 0.0136 | 0.6002 | 0.065 | 0.1522 |
| 0.102 | 0.1128 | 0.1672 | 0.6206 | 0.0958 | 0.1056 | 0.0108 | 0.3622 |
| 0.2434 | 0.467 | 0.257 | 0.0562 | 0.1952 | 0.4168 | 0.075 | 0.084 |
| 0.2642 | 0.0366 | 0.1306 | 0.4918 | 0.255 | 0.0532 | 0.0692 | 0.3576 |
| 0.2868 | 0.0104 | 0.2706 | 0.4344 | 0.2214 | 0.0042 | 0.0694 | 0.2334 |
| 0.1208 | 0.021 | 0.3432 | 0.434 | 0.0768 | 0.0224 | 0.1238 | 0.4054 |
| 0.042 | 0.932 | 0.026 | 0.0064 | 0.0382 | 0.913 | 0.0162 | 0.0126 |
| 0.0668 | 0.5148 | 0.0792 | 0.2826 | 0.062 | 0.5288 | 0.048 | 0.1874 |
| 0.027 | 0.2454 | 0.335 | 0.1286 | 0.0094 | 0.172 | 0.355 | 0.1828 |
| 0.0136 | 0.023 | 0.0252 | 0.9282 | 0.0108 | 0.0226 | 0.013 | 0.0698 |
| 0.2266 | 0.0078 | 0.103 | 0.6538 | 0.2104 | 0.0068 | 0.0262 | 0.1596 |
| 0.2062 | 0.0044 | 0.159 | 0.6336 | 0.198 | 0.004 | 0.0056 | 0.462 |
| 0.107 | 0.1476 | 0.2976 | 0.478 | 0.0534 | 0.0998 | 0.071 | 0.3088 |
| 0.0558 | 0.0386 | 0.1698 | 0.6422 | 0.051 | 0.0532 | 0.0836 | 0.3886 |
| 0.4966 | 0.0106 | 0.0468 | 0.4374 | 0.475 | 0.0124 | 0.029 | 0.1272 |
| 0.4048 | 0.0244 | 0.0526 | 0.5172 | 0.3734 | 0.0208 | 0.036 | 0.035 |
| 0.3424 | 0.013 | 0.139 | 0.4882 | 0.3244 | 0.0142 | 0.0342 | 0.3696 |
| 0.1838 | 0.0046 | 0.1218 | 0.6856 | 0.177 | 0.0052 | 0.0108 | 0.405 |
| 0.0876 | 0.0114 | 0.0368 | 0.8652 | 0.0778 | 0.0102 | 0.01 | 0.0482 |
| 0.1648 | 0.0024 | 0.0124 | 0.8248 | 0.1542 | 0.0024 | 0.0064 | 0.0482 |
| 0.0022 | 0.0078 | 0.0032 | 0.003 | 0.0112 | 0.1942 | 0.7882 | 0.0036 |
| 0.0182 | 0.0898 | 0.0066 | 0.0084 | 0.0376 | 0.2416 | 0.706 | 0.0066 |
| 0.0052 | 0.0178 | 0.03 | 0.0468 | 0.0094 | 0.1784 | 0.7356 | 0.0252 |
| 0.003 | 0.0136 | 0.0102 | 0.011 | 0.0076 | 0.1944 | 0.777 | 0.0098 |
| 0.0038 | 0.022 | 0.0048 | 0.0046 | 0.0218 | 0.1924 | 0.7766 | 0.0052 |
| 0.003 | 0.006 | 0.0028 | 0.0032 | 0.017 | 0.1868 | 0.7904 | 0.003 |
| 0.0184 | 0.0042 | 0.0044 | 0.0094 | 0.0518 | 0.1584 | 0.7764 | 0.006 |
| 0.0022 | 0.038 | 0.0076 | 0.0196 | 0.003 | 0.2126 | 0.7576 | 0.0098 |
| 0.0032 | 0.0318 | 0.0056 | 0.0062 | 0.0156 | 0.2066 | 0.7664 | 0.0052 |
| 0.0212 | 0.0078 | 0.0108 | 0.0716 | 0.0416 | 0.152 | 0.7236 | 0.0226 |
| 0.011 | 0.0104 | 0.0328 | 0.0456 | 0.0356 | 0.156 | 0.7304 | 0.019 |
| 0.0112 | 0.032 | 0.006 | 0.005 | 0.0326 | 0.1972 | 0.7592 | 0.0054 |
| 0.0314 | 0.0046 | 0.0084 | 0.0812 | 0.0608 | 0.1342 | 0.7154 | 0.0136 |
| 0.0098 | 0.0268 | 0.1044 | 0.0024 | 0.0148 | 0.1632 | 0.7152 | 0.0522 |
| 0.012 | 0.0058 | 0.0024 | 0.0026 | 0.0506 | 0.1612 | 0.783 | 0.0022 |
| 0.002 | 0.0152 | 0.004 | 0.0038 | 0.0034 | 0.2054 | 0.7838 | 0.004 |
| 0.002 | 0.0068 | 0.0114 | 0.0178 | 0.004 | 0.1864 | 0.7804 | 0.0126 |
| 0.0574 | 0.0098 | 0.0052 | 0.005 | 0.0776 | 0.1712 | 0.7406 | 0.0048 |
| 0.002 | 0.163 | 0.007 | 0.0118 | 0.0026 | 0.3194 | 0.6598 | 0.0076 |
| 0.0084 | 0.0054 | 0.0262 | 0.0442 | 0.0252 | 0.158 | 0.7466 | 0.0254 |
| 0.0278 | 0.0118 | 0.0044 | 0.0058 | 0.0538 | 0.174 | 0.7618 | 0.004 |

| | | | | | | | |
|---|---|---|---|---|---|---|---|
| 0.0114 | 0.012 | 0.0062 | 0.0052 | 0.0418 | 0.1712 | 0.7758 | 0.0064 |
| 0.005 | 0.1588 | 0.0062 | 0.0098 | 0.0086 | 0.317 | 0.6586 | 0.0084 |
| 0.008 | 0.039 | 0.006 | 0.0042 | 0.0298 | 0.2018 | 0.7584 | 0.0054 |
| 0.1036 | 0.0062 | 0.0644 | 0.7182 | 0.1044 | 0.0096 | 0.1038 | 0.1346 |
| 0.3974 | 0.005 | 0.0188 | 0.5946 | 0.3748 | 0.0044 | 0.007 | 0.0216 |
| 0.0058 | 0.0056 | 0.1966 | 0.7888 | 0.0046 | 0.0048 | 0.005 | 0.5654 |
| 0.0504 | 0.0064 | 0.1162 | 0.8224 | 0.0476 | 0.0058 | 0.0082 | 0.3088 |
| 0.3536 | 0.0622 | 0.0496 | 0.5058 | 0.3308 | 0.0702 | 0.0438 | 0.0448 |
| 0.2862 | 0.0264 | 0.3456 | 0.3488 | 0.1854 | 0.0052 | 0.115 | 0.1724 |
| 0.0068 | 0.2498 | 0.2272 | 0.4914 | 0.0038 | 0.1752 | 0.103 | 0.1508 |
| 0.111 | 0.0758 | 0.3346 | 0.518 | 0.0472 | 0.0242 | 0.0762 | 0.2658 |
| 0.032 | 0.0168 | 0.1718 | 0.7792 | 0.0222 | 0.0138 | 0.013 | 0.2884 |
| 0.1374 | 0.0076 | 0.0396 | 0.8146 | 0.123 | 0.0066 | 0.0158 | 0.0536 |
| 0.1414 | 0.0362 | 0.2994 | 0.5234 | 0.0782 | 0.0126 | 0.0864 | 0.2934 |
| 0.0032 | 0.0048 | 0.1788 | 0.7976 | 0.0032 | 0.0052 | 0.0158 | 0.5322 |
| 0.0066 | 0.0314 | 0.3024 | 0.6284 | 0.0024 | 0.0104 | 0.0566 | 0.4792 |
| 0.009 | 0.0044 | 0.1734 | 0.8108 | 0.0074 | 0.004 | 0.0048 | 0.3104 |
| 0.0164 | 0.0044 | 0.0574 | 0.921 | 0.0126 | 0.0042 | 0.005 | 0.2268 |
| 0.0088 | 0.0114 | 0.1612 | 0.7712 | 0.0074 | 0.0134 | 0.0468 | 0.3188 |
| 0.0034 | 0.0022 | 0.0596 | 0.931 | 0.003 | 0.0022 | 0.004 | 0.1566 |
| 0.2342 | 0.0066 | 0.042 | 0.719 | 0.2228 | 0.0062 | 0.0098 | 0.2428 |
| 0.2622 | 0.0048 | 0.144 | 0.5998 | 0.2474 | 0.0046 | 0.0046 | 0.3138 |
| 0.031 | 0.0176 | 0.0542 | 0.8036 | 0.0272 | 0.0244 | 0.0904 | 0.1598 |
| 0.007 | 0.0454 | 0.2776 | 0.641 | 0.0042 | 0.0178 | 0.0598 | 0.4394 |
| 0.0056 | 0.008 | 0.1126 | 0.8546 | 0.0044 | 0.0076 | 0.021 | 0.237 |
| 0.0038 | 0.9796 | 0.009 | 0.0108 | 0.003 | 0.9684 | 0.0092 | 0.0102 |
| 0.0654 | 0.0428 | 0.1786 | 0.7186 | 0.0524 | 0.0344 | 0.0162 | 0.4138 |
| 0.0394 | 0.0176 | 0.1416 | 0.7294 | 0.0382 | 0.0206 | 0.0702 | 0.3692 |
| 0.02 | 0.0022 | 0.0418 | 0.9364 | 0.0166 | 0.0022 | 0.0032 | 0.1966 |
| 0.011 | 0.018 | 0.1364 | 0.8198 | 0.0092 | 0.017 | 0.0172 | 0.2464 |
| 0.0148 | 0.0382 | 0.012 | 0.4542 | 0.0152 | 0.1214 | 0.3978 | 0.0152 |
| 0.018 | 0.065 | 0.0366 | 0.8716 | 0.0138 | 0.062 | 0.0158 | 0.069 |
| 0.0088 | 0.061 | 0.0494 | 0.8576 | 0.0068 | 0.0644 | 0.022 | 0.0832 |
| 0.006 | 0.0618 | 0.1744 | 0.5214 | 0.0042 | 0.083 | 0.2166 | 0.1754 |
| 0.0036 | 0.0626 | 0.3092 | 0.3074 | 0.0022 | 0.0758 | 0.3056 | 0.1996 |
| 0.0732 | 0.0828 | 0.0438 | 0.756 | 0.0614 | 0.096 | 0.0428 | 0.0452 |
| 0.0024 | 0.03 | 0.0234 | 0.8492 | 0.002 | 0.0446 | 0.0804 | 0.0506 |
| 0.1472 | 0.0876 | 0.1358 | 0.5784 | 0.132 | 0.0934 | 0.0608 | 0.2696 |
| 0.1208 | 0.1968 | 0.231 | 0.5114 | 0.0756 | 0.1344 | 0.0474 | 0.1994 |
| 0.0976 | 0.0044 | 0.0158 | 0.8644 | 0.0948 | 0.0046 | 0.0206 | 0.0398 |
| 0.1738 | 0.0206 | 0.0704 | 0.7386 | 0.1612 | 0.0188 | 0.011 | 0.2526 |
| 0.0062 | 0.0354 | 0.044 | 0.0466 | 0.0132 | 0.187 | 0.7092 | 0.0276 |
| 0.0928 | 0.5404 | 0.045 | 0.181 | 0.0936 | 0.5702 | 0.1102 | 0.088 |
| 0.2838 | 0.227 | 0.0204 | 0.4722 | 0.267 | 0.2286 | 0.0122 | 0.0266 |

| | | | | | | | |
|---|---|---|---|---|---|---|---|
| 0.0074 | 0.0906 | 0.1774 | 0.7124 | 0.004 | 0.0342 | 0.072 | 0.1046 |
| 0.1538 | 0.0394 | 0.0324 | 0.7644 | 0.1402 | 0.0418 | 0.0212 | 0.0898 |
| 0.0074 | 0.0088 | 0.019 | 0.446 | 0.0144 | 0.0912 | 0.4294 | 0.0506 |
| 0.0034 | 0.1268 | 0.0032 | 0.0022 | 0.0042 | 0.2992 | 0.6914 | 0.0024 |
| 0.1938 | 0.3436 | 0.1264 | 0.1078 | 0.1768 | 0.3638 | 0.2256 | 0.129 |
| 0.0142 | 0.0098 | 0.0204 | 0.0664 | 0.023 | 0.1732 | 0.717 | 0.0354 |
| 0.006 | 0.032 | 0.0088 | 0.0032 | 0.0244 | 0.2018 | 0.7614 | 0.0064 |
| 0.0158 | 0.0276 | 0.0048 | 0.0042 | 0.035 | 0.1972 | 0.7594 | 0.0042 |
| 0.0178 | 0.7456 | 0.0484 | 0.1432 | 0.0134 | 0.7494 | 0.0458 | 0.131 |
| 0.0314 | 0.0786 | 0.0496 | 0.162 | 0.0388 | 0.1892 | 0.5606 | 0.036 |
| 0.0022 | 0.006 | 0.0074 | 0.0062 | 0.0086 | 0.1886 | 0.7894 | 0.0098 |
| 0.0108 | 0.0076 | 0.0036 | 0.003 | 0.0312 | 0.1812 | 0.7816 | 0.0032 |
| 0.0026 | 0.0114 | 0.0068 | 0.0052 | 0.01 | 0.1938 | 0.7844 | 0.007 |
| 0.0196 | 0.0416 | 0.0088 | 0.5434 | 0.0222 | 0.1088 | 0.3166 | 0.0088 |
| 0.0192 | 0.0076 | 0.0954 | 0.4472 | 0.02 | 0.0778 | 0.3594 | 0.1318 |
| 0.0044 | 0.0876 | 0.0142 | 0.051 | 0.0098 | 0.2352 | 0.6896 | 0.0164 |
| 0.0028 | 0.0284 | 0.0034 | 0.003 | 0.0096 | 0.2144 | 0.769 | 0.0036 |
| 0.0194 | 0.2066 | 0.0126 | 0.487 | 0.0178 | 0.2532 | 0.2288 | 0.0222 |
| 0.0868 | 0.4988 | 0.1922 | 0.2594 | 0.0486 | 0.4314 | 0.0688 | 0.1216 |
| 0.3312 | 0.0352 | 0.14 | 0.4454 | 0.2996 | 0.0404 | 0.0746 | 0.0954 |
| 0.1118 | 0.0142 | 0.1234 | 0.7514 | 0.1058 | 0.0124 | 0.0068 | 0.3474 |
| 0.003 | 0.006 | 0.013 | 0.974 | 0.0022 | 0.0052 | 0.0054 | 0.028 |
| 0.1142 | 0.0148 | 0.1452 | 0.7234 | 0.1056 | 0.0162 | 0.0094 | 0.4524 |
| 0.019 | 0.0612 | 0.2078 | 0.6912 | 0.0102 | 0.0336 | 0.0574 | 0.3346 |
| 0.003 | 0.0638 | 0.0124 | 0.9172 | 0.002 | 0.0602 | 0.008 | 0.0344 |
| 0.0032 | 0.0168 | 0.1926 | 0.7782 | 0.0022 | 0.0146 | 0.0116 | 0.4264 |
| 0.0202 | 0.2394 | 0.0454 | 0.6954 | 0.012 | 0.2276 | 0.0198 | 0.048 |
| 0.2316 | 0.0112 | 0.0642 | 0.6494 | 0.2232 | 0.0162 | 0.0468 | 0.1818 |
| 0.0164 | 0.0166 | 0.195 | 0.7148 | 0.015 | 0.0178 | 0.0572 | 0.434 |
| 0.0876 | 0.023 | 0.137 | 0.692 | 0.073 | 0.0278 | 0.0702 | 0.155 |
| 0.186 | 0.0044 | 0.1108 | 0.696 | 0.1744 | 0.004 | 0.0148 | 0.206 |
| 0.03 | 0.049 | 0.032 | 0.8602 | 0.0252 | 0.0514 | 0.0312 | 0.135 |
| 0.5846 | 0.0164 | 0.074 | 0.2228 | 0.5554 | 0.0384 | 0.1094 | 0.1948 |
| 0.308 | 0.01 | 0.019 | 0.6694 | 0.2906 | 0.0084 | 0.0128 | 0.0242 |
| 0.011 | 0.0474 | 0.1788 | 0.749 | 0.0086 | 0.0492 | 0.0152 | 0.4762 |
| 0.0444 | 0.039 | 0.1886 | 0.7274 | 0.0316 | 0.0284 | 0.0238 | 0.4036 |
| 0.3352 | 0.0058 | 0.1122 | 0.538 | 0.321 | 0.006 | 0.0236 | 0.1886 |
| 0.0162 | 0.0104 | 0.0544 | 0.8984 | 0.0146 | 0.0112 | 0.0214 | 0.1336 |
| 0.0274 | 0.251 | 0.0748 | 0.6484 | 0.0202 | 0.247 | 0.0096 | 0.1234 |
| 0.0098 | 0.0144 | 0.1918 | 0.6874 | 0.0094 | 0.0204 | 0.0908 | 0.5046 |
| 0.0036 | 0.0272 | 0.0404 | 0.9158 | 0.003 | 0.0262 | 0.0148 | 0.0734 |
| 0.0412 | 0.0172 | 0.1266 | 0.7866 | 0.037 | 0.0196 | 0.0302 | 0.1914 |
| 0.0046 | 0.0388 | 0.1458 | 0.8044 | 0.0034 | 0.0382 | 0.0084 | 0.3226 |
| 0.1118 | 0.1148 | 0.0238 | 0.7288 | 0.0998 | 0.1262 | 0.0212 | 0.0442 |

| | | | | | | | |
|---|---|---|---|---|---|---|---|
| 0.0254 | 0.1028 | 0.1236 | 0.7548 | 0.0156 | 0.0606 | 0.0444 | 0.0642 |
| 0.0254 | 0.0042 | 0.081 | 0.8898 | 0.0208 | 0.0038 | 0.0048 | 0.2376 |
| 0.0028 | 0.225 | 0.1606 | 0.6014 | 0.0022 | 0.2212 | 0.0146 | 0.423 |
| 0.1352 | 0.0498 | 0.1654 | 0.6584 | 0.0964 | 0.0218 | 0.0582 | 0.1186 |
| 0.0078 | 0.1192 | 0.172 | 0.6918 | 0.0058 | 0.1168 | 0.0136 | 0.4874 |
| 0.0134 | 0.1134 | 0.0242 | 0.8128 | 0.0108 | 0.1188 | 0.0334 | 0.0782 |
| 0.0892 | 0.0426 | 0.248 | 0.5992 | 0.0638 | 0.0344 | 0.0544 | 0.4826 |
| 0.0022 | 0.0058 | 0.1852 | 0.7958 | 0.002 | 0.0058 | 0.0114 | 0.5132 |
| 0.0514 | 0.012 | 0.136 | 0.7946 | 0.043 | 0.0118 | 0.0146 | 0.179 |
| 0.0614 | 0.0132 | 0.1026 | 0.79 | 0.0582 | 0.0144 | 0.0352 | 0.1524 |
| 0.4292 | 0.093 | 0.0232 | 0.4702 | 0.4036 | 0.095 | 0.0078 | 0.0438 |
| 0.373 | 0.0234 | 0.1054 | 0.4876 | 0.3464 | 0.0246 | 0.0364 | 0.1922 |
| 0.0058 | 0.09 | 0.009 | 0.0062 | 0.025 | 0.2492 | 0.7108 | 0.008 |
| 0.0152 | 0.0322 | 0.0298 | 0.1164 | 0.0296 | 0.1682 | 0.6556 | 0.0338 |
| 0.0064 | 0.3796 | 0.1212 | 0.4102 | 0.0046 | 0.3906 | 0.0734 | 0.2812 |
| 0.3392 | 0.4558 | 0.0948 | 0.1224 | 0.3056 | 0.4276 | 0.0494 | 0.0692 |
| 0.0244 | 0.0088 | 0.0042 | 0.004 | 0.0454 | 0.1764 | 0.7698 | 0.0036 |
| 0.0028 | 0.0122 | 0.007 | 0.0062 | 0.0142 | 0.1916 | 0.781 | 0.0062 |
| 0.003 | 0.0574 | 0.0094 | 0.009 | 0.006 | 0.2318 | 0.7438 | 0.011 |
| 0.5808 | 0.0034 | 0.0716 | 0.3622 | 0.5552 | 0.0034 | 0.0078 | 0.0952 |
| 0.382 | 0.0138 | 0.016 | 0.6012 | 0.3646 | 0.0128 | 0.0052 | 0.0428 |
| 0.0022 | 0.0028 | 0.024 | 0.9634 | 0.002 | 0.0028 | 0.008 | 0.089 |
| 0.0092 | 0.0366 | 0.0258 | 0.034 | 0.0214 | 0.1892 | 0.7294 | 0.0224 |
| 0.0092 | 0.1284 | 0.118 | 0.0046 | 0.0132 | 0.1744 | 0.6898 | 0.0506 |
| 0.0112 | 0.0186 | 0.0102 | 0.0064 | 0.0272 | 0.1866 | 0.7698 | 0.0108 |
| 0.003 | 0.0038 | 0.0052 | 0.0078 | 0.0184 | 0.177 | 0.7918 | 0.0054 |
| 0.0504 | 0.0088 | 0.1922 | 0.7156 | 0.0218 | 0.0044 | 0.0656 | 0.2174 |
| 0.0046 | 0.0196 | 0.0064 | 0.0086 | 0.019 | 0.193 | 0.7732 | 0.0078 |
| 0.0156 | 0.0142 | 0.0172 | 0.0108 | 0.0294 | 0.1756 | 0.767 | 0.0178 |
| 0.0158 | 0.0238 | 0.0592 | 0.0692 | 0.0226 | 0.166 | 0.6828 | 0.038 |
| 0.0062 | 0.0122 | 0.0236 | 0.0542 | 0.013 | 0.1754 | 0.7342 | 0.0248 |
| 0.2058 | 0.1412 | 0.148 | 0.501 | 0.1822 | 0.1322 | 0.0364 | 0.1838 |
| 0.004 | 0.0268 | 0.059 | 0.884 | 0.0032 | 0.0252 | 0.0286 | 0.1802 |

| X6.Cluster2 | X6.Cluster3 | X6.Cluster4 | X6.Cluster5 | X6.Cluster6 |
|---|---|---|---|---|
| 0.256 | 0.0226 | 0.0088 | 0.019 | 0.4664 |
| 0.0314 | 0.0716 | 0.0072 | 0.1086 | 0.6762 |
| 0.278 | 0.0722 | 0.0306 | 0.0268 | 0.4366 |
| 0.243 | 0.0172 | 0.0258 | 0.046 | 0.443 |
| 0.0376 | 0.0142 | 0.0214 | 0.0188 | 0.8822 |
| 0.3876 | 0.1496 | 0.0258 | 0.045 | 0.2292 |
| 0.314 | 0.4406 | 0.052 | 0.0114 | 0.1664 |
| 0.3276 | 0.0578 | 0.0528 | 0.0152 | 0.5136 |
| 0.1388 | 0.0216 | 0.0248 | 0.012 | 0.7282 |
| 0.2518 | 0.0564 | 0.0054 | 0.0746 | 0.5452 |
| 0.2814 | 0.3388 | 0.0064 | 0.1004 | 0.1996 |
| 0.0038 | 0.0258 | 0.007 | 0.0498 | 0.8672 |
| 0.0814 | 0.103 | 0.0064 | 0.0234 | 0.6936 |
| 0.0136 | 0.0238 | 0.0456 | 0.0214 | 0.8776 |
| 0.0048 | 0.0594 | 0.0136 | 0.1068 | 0.699 |
| 0.0362 | 0.0412 | 0.1408 | 0.0552 | 0.6034 |
| 0.0136 | 0.0288 | 0.058 | 0.0234 | 0.8692 |
| 0.0178 | 0.025 | 0.0396 | 0.0134 | 0.8846 |
| 0.0188 | 0.08 | 0.0056 | 0.0114 | 0.8684 |
| 0.0064 | 0.0666 | 0.0238 | 0.0084 | 0.8836 |
| 0.071 | 0.0068 | 0.0514 | 0.049 | 0.7944 |
| 0.3692 | 0.3702 | 0.0164 | 0.0296 | 0.1664 |
| 0.105 | 0.0174 | 0.0442 | 0.0414 | 0.712 |
| 0.028 | 0.1314 | 0.1012 | 0.1094 | 0.5966 |
| 0.0942 | 0.1296 | 0.0288 | 0.0526 | 0.643 |
| 0.1666 | 0.1112 | 0.0226 | 0.0234 | 0.6424 |
| 0.0096 | 0.007 | 0.0158 | 0.0086 | 0.9444 |
| 0.0418 | 0.0404 | 0.0864 | 0.0256 | 0.7926 |
| 0.0068 | 0.0438 | 0.0026 | 0.0766 | 0.7898 |
| 0.0708 | 0.0166 | 0.2414 | 0.0142 | 0.6002 |
| 0.0104 | 0.0144 | 0.0254 | 0.0278 | 0.905 |
| 0.0376 | 0.0314 | 0.2094 | 0.0188 | 0.678 |
| 0.0052 | 0.0076 | 0.0236 | 0.0056 | 0.9492 |
| 0.0614 | 0.0458 | 0.009 | 0.0832 | 0.7158 |
| 0.005 | 0.0626 | 0.0128 | 0.0628 | 0.8024 |
| 0.0058 | 0.0056 | 0.0424 | 0.033 | 0.9062 |
| 0.004 | 0.0322 | 0.0272 | 0.0828 | 0.7764 |
| 0.0718 | 0.2508 | 0.0062 | 0.0094 | 0.6522 |
| 0.047 | 0.1118 | 0.0042 | 0.01 | 0.8182 |
| 0.0042 | 0.04 | 0.0086 | 0.079 | 0.7792 |
| 0.0048 | 0.0064 | 0.0482 | 0.0092 | 0.9196 |

| | | | | |
|---|---|---|---|---|
| 0.0028 | 0.0434 | 0.0866 | 0.0384 | 0.8234 |
| 0.0966 | 0.0216 | 0.0058 | 0.0078 | 0.8086 |
| 0.0752 | 0.0354 | 0.0334 | 0.0432 | 0.704 |
| 0.0818 | 0.0618 | 0.004 | 0.087 | 0.6716 |
| 0.003 | 0.0214 | 0.0074 | 0.041 | 0.8902 |
| 0.2404 | 0.031 | 0.0166 | 0.0424 | 0.6256 |
| 0.0076 | 0.0138 | 0.091 | 0.026 | 0.8342 |
| 0.0184 | 0.0126 | 0.0052 | 0.008 | 0.9428 |
| 0.015 | 0.0322 | 0.0476 | 0.04 | 0.7846 |
| 0.024 | 0.021 | 0.1194 | 0.0068 | 0.8072 |
| 0.342 | 0.0296 | 0.005 | 0.0132 | 0.5228 |
| 0.027 | 0.0418 | 0.0478 | 0.0172 | 0.7954 |
| 0.2928 | 0.124 | 0.0372 | 0.0634 | 0.4094 |
| 0.2626 | 0.0214 | 0.1256 | 0.0214 | 0.509 |
| 0.0832 | 0.0644 | 0.038 | 0.0966 | 0.603 |
| 0.0206 | 0.0202 | 0.0138 | 0.0122 | 0.9008 |
| 0.1566 | 0.066 | 0.0094 | 0.0168 | 0.5952 |
| 0.004 | 0.0508 | 0.0078 | 0.104 | 0.7162 |
| 0.1022 | 0.0102 | 0.0148 | 0.0188 | 0.778 |
| 0.1776 | 0.0206 | 0.0118 | 0.013 | 0.6334 |
| 0.0542 | 0.061 | 0.0106 | 0.0322 | 0.8064 |
| 0.1188 | 0.0732 | 0.0592 | 0.02 | 0.6844 |
| 0.0148 | 0.0162 | 0.0142 | 0.0092 | 0.937 |
| 0.019 | 0.0104 | 0.036 | 0.0118 | 0.8112 |
| 0.263 | 0.0112 | 0.004 | 0.0082 | 0.6484 |
| 0.2522 | 0.172 | 0.0036 | 0.0132 | 0.5392 |
| 0.0138 | 0.0208 | 0.0144 | 0.025 | 0.8914 |
| 0.052 | 0.042 | 0.0086 | 0.0238 | 0.831 |
| 0.0122 | 0.0112 | 0.1546 | 0.0234 | 0.7746 |
| 0.1514 | 0.2176 | 0.0486 | 0.013 | 0.5458 |
| 0.3836 | 0.0476 | 0.0044 | 0.03 | 0.0332 |
| 0.3916 | 0.0096 | 0.0134 | 0.0142 | 0.1038 |
| 0.5912 | 0.1176 | 0.0094 | 0.0228 | 0.0266 |
| 0.639 | 0.1034 | 0.0034 | 0.0196 | 0.1996 |
| 0.5076 | 0.2692 | 0.0198 | 0.0496 | 0.0036 |
| 0.5464 | 0.4138 | 0.0074 | 0.011 | 0.0026 |
| 0.3866 | 0.0136 | 0.0134 | 0.015 | 0.0096 |
| 0.3536 | 0.245 | 0.0048 | 0.1226 | 0.0102 |
| 0.266 | 0.2346 | 0.0608 | 0.0282 | 0.312 |
| 0.6668 | 0.0364 | 0.006 | 0.011 | 0.2064 |
| 0.5288 | 0.1808 | 0.0038 | 0.0144 | 0.2456 |
| 0.4472 | 0.2248 | 0.0104 | 0.0142 | 0.0012 |
| 0.419 | 0.49 | 0.024 | 0.0072 | 0.0418 |
| 0.6444 | 0.0372 | 0.0414 | 0.0224 | 0.0026 |

| | | | | |
|---|---|---|---|---|
| 0.4462 | 0.1494 | 0.0128 | 0.007 | 0.0076 |
| 0.7306 | 0.0672 | 0.0144 | 0.0112 | 0.0036 |
| 0.4082 | 0.021 | 0.006 | 0.0292 | 0.0102 |
| 0.2836 | 0.0742 | 0.017 | 0.1384 | 0.1396 |
| 0.6872 | 0.0472 | 0.0118 | 0.0052 | 0.0732 |
| 0.4902 | 0.4048 | 0.0044 | 0.0088 | 0.016 |
| 0.5976 | 0.0804 | 0.0082 | 0.0076 | 0.1046 |
| 0.6802 | 0.0208 | 0.0206 | 0.0214 | 0.0102 |
| 0.7918 | 0.049 | 0.009 | 0.0176 | 0.0718 |
| 0.4216 | 0.0808 | 0.0202 | 0.1666 | 0.0292 |
| 0.4052 | 0.191 | 0.0236 | 0.021 | 0.002 |
| 0.3804 | 0.32 | 0.0246 | 0.1554 | 0.002 |
| 0.6482 | 0.1378 | 0.0106 | 0.0984 | 0.0178 |
| 0.752 | 0.176 | 0.0086 | 0.025 | 0.0082 |
| 0.4168 | 0.0912 | 0.0208 | 0.0302 | 0.004 |
| 0.4684 | 0.0208 | 0.0044 | 0.0088 | 0.079 |
| 0.4322 | 0.0666 | 0.0548 | 0.029 | 0.1698 |
| 0.533 | 0.0816 | 0.0206 | 0.1408 | 0.0144 |
| 0.4928 | 0.3624 | 0.0072 | 0.043 | 0.07 |
| 0.3606 | 0.0158 | 0.0024 | 0.006 | 0.1038 |
| 0.5948 | 0.1978 | 0.0078 | 0.01 | 0.1248 |
| 0.2006 | 0.1146 | 0.0158 | 0.2228 | 0.1868 |
| 0.3258 | 0.013 | 0.042 | 0.0934 | 0.101 |
| 0.1982 | 0.1026 | 0.144 | 0.1756 | 0.0178 |
| 0.3774 | 0.3402 | 0.0128 | 0.0158 | 0.0114 |
| 0.5142 | 0.259 | 0.011 | 0.0692 | 0.014 |
| 0.6566 | 0.0136 | 0.0048 | 0.009 | 0.1794 |
| 0.5664 | 0.0314 | 0.007 | 0.0322 | 0.141 |
| 0.37 | 0.358 | 0.0742 | 0.1164 | 0.061 |
| 0.1844 | 0.0278 | 0.0402 | 0.0246 | 0.3714 |
| 0.6892 | 0.1484 | 0.02 | 0.0296 | 0.0484 |
| 0.4526 | 0.238 | 0.0096 | 0.011 | 0.0804 |
| 0.384 | 0.4236 | 0.0808 | 0.0446 | 0.0066 |
| 0.5176 | 0.0264 | 0.0074 | 0.0576 | 0.1406 |
| 0.4016 | 0.0124 | 0.0086 | 0.0078 | 0.0148 |
| 0.369 | 0.0636 | 0.0162 | 0.0088 | 0.1612 |
| 0.8022 | 0.0554 | 0.007 | 0.065 | 0.012 |
| 0.4468 | 0.0322 | 0.0534 | 0.0536 | 0.008 |
| 0.7602 | 0.0808 | 0.0138 | 0.0354 | 0.0032 |
| 0.6392 | 0.0748 | 0.0986 | 0.051 | 0.0288 |
| 0.7386 | 0.0354 | 0.0472 | 0.03 | 0.0054 |
| 0.605 | 0.039 | 0.0034 | 0.0074 | 0.0874 |
| 0.521 | 0.1496 | 0.003 | 0.0166 | 0.0046 |
| 0.3696 | 0.0772 | 0.017 | 0.018 | 0.002 |

| | | | | |
|---|---|---|---|---|
| 0.3732 | 0.3498 | 0.0494 | 0.0512 | 0.0752 |
| 0.394 | 0.0332 | 0.0248 | 0.0662 | 0.0212 |
| 0.216 | 0.1712 | 0.036 | 0.2156 | 0.0558 |
| 0.5494 | 0.1458 | 0.0054 | 0.0136 | 0.0064 |
| 0.4046 | 0.041 | 0.0072 | 0.0172 | 0.1154 |
| 0.4986 | 0.152 | 0.0116 | 0.0842 | 0.0434 |
| 0.697 | 0.214 | 0.0046 | 0.0074 | 0.0014 |
| 0.6926 | 0.0194 | 0.0022 | 0.0106 | 0.0018 |
| 0.7244 | 0.2104 | 0.002 | 0.0036 | 0.0138 |
| 0.3012 | 0.015 | 0.0072 | 0.0096 | 0.2622 |
| 0.574 | 0.0314 | 0.031 | 0.0292 | 0.3102 |
| 0.2816 | 0.2788 | 0.007 | 0.0238 | 0.3012 |
| 0.4632 | 0.097 | 0.01 | 0.1226 | 0.121 |
| 0.5042 | 0.3598 | 0.0036 | 0.0056 | 0.0056 |
| 0.2324 | 0.2614 | 0.063 | 0.186 | 0.058 |
| 0.2276 | 0.0148 | 0.038 | 0.0232 | 0.3324 |
| 0.652 | 0.3158 | 0.0054 | 0.0108 | 0.004 |
| 0.547 | 0.0942 | 0.0074 | 0.1528 | 0.0022 |
| 0.51 | 0.2678 | 0.0588 | 0.037 | 0.0082 |
| 0.4466 | 0.1348 | 0.004 | 0.0132 | 0.0162 |
| 0.6068 | 0.2346 | 0.0068 | 0.0088 | 0.0364 |
| 0.5662 | 0.0444 | 0.0416 | 0.0628 | 0.0666 |
| 0.7298 | 0.0368 | 0.1108 | 0.0678 | 0.037 |
| 0.4082 | 0.2552 | 0.0096 | 0.1388 | 0.0484 |
| 0.4424 | 0.0302 | 0.0024 | 0.009 | 0.0608 |
| 0.6808 | 0.1466 | 0.0068 | 0.046 | 0.013 |
| 0.6888 | 0.0348 | 0.0196 | 0.0232 | 0.0066 |
| 0.45 | 0.3014 | 0.0116 | 0.0666 | 0.0026 |
| 0.465 | 0.035 | 0.0178 | 0.029 | 0.0312 |
| 0.4334 | 0.156 | 0.0154 | 0.012 | 0.0036 |
| 0.6156 | 0.1714 | 0.0308 | 0.0534 | 0.0042 |
| 0.748 | 0.0782 | 0.0052 | 0.0114 | 0.0264 |
| 0.4512 | 0.1312 | 0.0028 | 0.116 | 0.0016 |
| 0.5438 | 0.0908 | 0.0154 | 0.0136 | 0.0042 |
| 0.7682 | 0.0948 | 0.0212 | 0.0394 | 0.003 |
| 0.772 | 0.0438 | 0.0072 | 0.0112 | 0.1418 |
| 0.753 | 0.0204 | 0.0254 | 0.0382 | 0.0128 |
| 0.4348 | 0.0226 | 0.009 | 0.0376 | 0.0116 |
| 0.3914 | 0.0132 | 0.0066 | 0.017 | 0.0378 |
| 0.395 | 0.1564 | 0.0346 | 0.153 | 0.0532 |
| 0.5704 | 0.0936 | 0.0636 | 0.0422 | 0.0172 |
| 0.713 | 0.0262 | 0.0238 | 0.033 | 0.0252 |
| 0.4592 | 0.4 | 0.0116 | 0.033 | 0.0186 |
| 0.4062 | 0.0924 | 0.0186 | 0.0132 | 0.0112 |

| | | | | |
|---|---|---|---|---|
| 0.391 | 0.112 | 0.005 | 0.0366 | 0.1578 |
| 0.415 | 0.4518 | 0.0058 | 0.0712 | 0.004 |
| 0.4232 | 0.1604 | 0.004 | 0.1694 | 0.006 |
| 0.572 | 0.1848 | 0.0024 | 0.0076 | 0.006 |
| 0.6234 | 0.0906 | 0.0058 | 0.1418 | 0.0022 |
| 0.491 | 0.4584 | 0.006 | 0.0208 | 0.0088 |
| 0.2522 | 0.1268 | 0.0116 | 0.218 | 0.05 |
| 0.5262 | 0.0244 | 0.04 | 0.0218 | 0.0112 |
| 0.3602 | 0.0332 | 0.0728 | 0.0566 | 0.247 |
| 0.2334 | 0.0128 | 0.0152 | 0.0116 | 0.3742 |
| 0.5596 | 0.0556 | 0.0086 | 0.009 | 0.0028 |
| 0.3832 | 0.1794 | 0.0204 | 0.0162 | 0.007 |
| 0.5328 | 0.0114 | 0.0126 | 0.0102 | 0.0026 |
| 0.41 | 0.0124 | 0.1724 | 0.0684 | 0.0472 |
| 0.5054 | 0.1662 | 0.0222 | 0.011 | 0.0996 |
| 0.479 | 0.1978 | 0.0058 | 0.0152 | 0.0044 |
| 0.7336 | 0.0372 | 0.0214 | 0.0396 | 0.0036 |
| 0.4074 | 0.2668 | 0.006 | 0.0278 | 0.0096 |
| 0.4762 | 0.0374 | 0.0076 | 0.0084 | 0.2 |
| 0.4086 | 0.2122 | 0.0576 | 0.077 | 0.2272 |
| 0.6518 | 0.1168 | 0.0448 | 0.0318 | 0.0076 |
| 0.3692 | 0.4088 | 0.0628 | 0.016 | 0.0164 |
| 0.3786 | 0.0174 | 0.0138 | 0.008 | 0.1012 |
| 0.3842 | 0.1896 | 0.0172 | 0.0932 | 0.146 |
| 0.4624 | 0.3668 | 0.1034 | 0.0332 | 0.003 |
| 0.5434 | 0.4066 | 0.006 | 0.0214 | 0.0036 |
| 0.383 | 0.0538 | 0.019 | 0.0102 | 0.0036 |
| 0.6226 | 0.023 | 0.0154 | 0.0172 | 0.0768 |
| 0.3712 | 0.0256 | 0.034 | 0.0632 | 0.026 |
| 0.2376 | 0.0128 | 0.202 | 0.0314 | 0.0518 |
| 0.3488 | 0.0462 | 0.0188 | 0.092 | 0.0178 |
| 0.3866 | 0.0156 | 0.013 | 0.0112 | 0.0366 |
| 0.2228 | 0.0494 | 0.4532 | 0.0262 | 0.0056 |
| 0.4528 | 0.5044 | 0.0058 | 0.0114 | 0.0028 |
| 0.281 | 0.1538 | 0.2564 | 0.0696 | 0.0022 |
| 0.4076 | 0.233 | 0.0142 | 0.0208 | 0.0296 |
| 0.3844 | 0.093 | 0.0106 | 0.192 | 0.0032 |
| 0.3578 | 0.0254 | 0.0544 | 0.0156 | 0.0168 |
| 0.4446 | 0.3076 | 0.0358 | 0.0116 | 0.0042 |
| 0.3818 | 0.1048 | 0.0072 | 0.0212 | 0.0066 |
| 0.0702 | 0.0196 | 0.0066 | 0.009 | 0.8758 |
| 0.8404 | 0.0682 | 0.0046 | 0.007 | 0.0064 |
| 0.3672 | 0.2322 | 0.0062 | 0.2138 | 0.013 |
| 0.4142 | 0.1768 | 0.0152 | 0.1872 | 0.0056 |

| | | | | |
|---|---|---|---|---|
| 0.354 | 0.049 | 0.01 | 0.0162 | 0.225 |
| 0.5012 | 0.44 | 0.0082 | 0.0112 | 0.0036 |
| 0.3454 | 0.211 | 0.0168 | 0.1986 | 0.0686 |
| 0.7436 | 0.0604 | 0.0068 | 0.025 | 0.023 |
| 0.6866 | 0.101 | 0.013 | 0.0256 | 0.0044 |
| 0.6606 | 0.1358 | 0.0728 | 0.01 | 0.003 |
| 0.5026 | 0.4622 | 0.0084 | 0.0074 | 0.005 |
| 0.2406 | 0.0074 | 0.0294 | 0.0314 | 0.3268 |
| 0.4644 | 0.446 | 0.0108 | 0.0164 | 0.0064 |
| 0.5262 | 0.2814 | 0.1306 | 0.0176 | 0.0072 |
| 0.3636 | 0.0202 | 0.029 | 0.0584 | 0.006 |
| 0.292 | 0.0682 | 0.0874 | 0.155 | 0.005 |
| 0.4994 | 0.4168 | 0.0054 | 0.0092 | 0.002 |
| 0.2952 | 0.1366 | 0.2964 | 0.0208 | 0.1384 |
| 0.581 | 0.061 | 0.0254 | 0.018 | 0.0176 |
| 0.432 | 0.342 | 0.0044 | 0.115 | 0.0084 |
| 0.6852 | 0.0872 | 0.1174 | 0.079 | 0.0072 |
| 0.6652 | 0.075 | 0.04 | 0.0348 | 0.005 |
| 0.5934 | 0.2918 | 0.0356 | 0.0314 | 0.0056 |
| 0.4714 | 0.4066 | 0.0276 | 0.0444 | 0.0034 |
| 0.2864 | 0.252 | 0.0646 | 0.0158 | 0.2196 |
| 0.3116 | 0.016 | 0.008 | 0.0104 | 0.2836 |
| 0.6238 | 0.164 | 0.0216 | 0.0388 | 0.1112 |
| 0.3368 | 0.2662 | 0.0106 | 0.1534 | 0.0036 |
| 0.3834 | 0.0238 | 0.0152 | 0.018 | 0.3986 |
| 0.3938 | 0.0468 | 0.0172 | 0.0236 | 0.0082 |
| 0.504 | 0.1598 | 0.031 | 0.1148 | 0.0144 |
| 0.6028 | 0.061 | 0.0216 | 0.0184 | 0.0888 |
| 0.6568 | 0.0206 | 0.0046 | 0.0086 | 0.004 |
| 0.5546 | 0.2382 | 0.0032 | 0.0082 | 0.0018 |
| 0.3386 | 0.0774 | 0.0048 | 0.0968 | 0.0058 |
| 0.1592 | 0.3026 | 0.1398 | 0.018 | 0.3676 |
| 0.1826 | 0.1034 | 0.3736 | 0.1866 | 0.004 |
| 0.569 | 0.0126 | 0.191 | 0.0324 | 0.0216 |
| 0.27 | 0.2796 | 0.023 | 0.0104 | 0.3312 |
| 0.5774 | 0.1404 | 0.0082 | 0.0278 | 0.1496 |
| 0.887 | 0.0214 | 0.0048 | 0.005 | 0.013 |
| 0.6086 | 0.145 | 0.017 | 0.0134 | 0.003 |
| 0.2116 | 0.3584 | 0.0262 | 0.0382 | 0.2782 |
| 0.4388 | 0.3856 | 0.0116 | 0.0162 | 0.097 |
| 0.6542 | 0.1362 | 0.0154 | 0.0398 | 0.0218 |
| 0.5778 | 0.1668 | 0.0132 | 0.0186 | 0.0022 |
| 0.9022 | 0.029 | 0.0042 | 0.0058 | 0.0034 |
| 0.483 | 0.0202 | 0.0352 | 0.0094 | 0.0722 |

| | | | | |
|---|---|---|---|---|
| 0.334 | 0.0376 | 0.0044 | 0.0402 | 0.0742 |
| 0.353 | 0.0656 | 0.0086 | 0.112 | 0.002 |
| 0.317 | 0.1446 | 0.0772 | 0.0306 | 0.1254 |
| 0.2664 | 0.1702 | 0.0106 | 0.0176 | 0.2716 |
| 0.1688 | 0.0958 | 0.1526 | 0.1784 | 0.1284 |
| 0.4938 | 0.4252 | 0.01 | 0.0236 | 0.0052 |
| 0.4758 | 0.0376 | 0.0062 | 0.0152 | 0.2102 |
| 0.2506 | 0.0142 | 0.0394 | 0.015 | 0.4544 |
| 0.2874 | 0.119 | 0.0376 | 0.046 | 0.2732 |
| 0.3594 | 0.1506 | 0.0492 | 0.028 | 0.3884 |
| 0.328 | 0.0322 | 0.0058 | 0.0144 | 0.259 |
| 0.4172 | 0.3678 | 0.0032 | 0.0066 | 0.1858 |
| 0.1316 | 0.015 | 0.5152 | 0.0086 | 0.2574 |
| 0.6022 | 0.0898 | 0.009 | 0.0166 | 0.0044 |
| 0.3616 | 0.2444 | 0.1456 | 0.012 | 0.0024 |
| 0.4956 | 0.0432 | 0.0086 | 0.0136 | 0.0056 |
| 0.0266 | 0.0098 | 0.8114 | 0.124 | 0.0022 |
| 0.0182 | 0.014 | 0.8428 | 0.0136 | 0.0938 |
| 0.0084 | 0.0544 | 0.8902 | 0.014 | 0.0264 |
| 0.6974 | 0.112 | 0.0034 | 0.0946 | 0.0218 |
| 0.0078 | 0.0048 | 0.9424 | 0.0172 | 0.019 |
| 0.061 | 0.0142 | 0.8454 | 0.015 | 0.0028 |
| 0.0238 | 0.019 | 0.8598 | 0.016 | 0.0522 |
| 0.0242 | 0.0414 | 0.8338 | 0.0354 | 0.033 |
| 0.0346 | 0.0604 | 0.8004 | 0.034 | 0.0068 |
| 0.0228 | 0.0296 | 0.8906 | 0.0146 | 0.01 |
| 0.01 | 0.0108 | 0.8926 | 0.077 | 0.0018 |
| 0.0064 | 0.0114 | 0.803 | 0.1342 | 0.0396 |
| 0.0088 | 0.0134 | 0.9298 | 0.0204 | 0.0012 |
| 0.4208 | 0.265 | 0.023 | 0.0298 | 0.002 |
| 0.0354 | 0.0272 | 0.8788 | 0.037 | 0.0026 |
| 0.0186 | 0.0304 | 0.8744 | 0.022 | 0.0262 |
| 0.036 | 0.092 | 0.8218 | 0.018 | 0.0024 |
| 0.0164 | 0.0418 | 0.8476 | 0.062 | 0.0084 |
| 0.0296 | 0.0184 | 0.8936 | 0.032 | 0.0066 |
| 0.2194 | 0.2724 | 0.3302 | 0.1398 | 0.008 |
| 0.008 | 0.0498 | 0.9154 | 0.0078 | 0.0088 |
| 0.0114 | 0.0124 | 0.897 | 0.0372 | 0.024 |
| 0.0108 | 0.025 | 0.8458 | 0.102 | 0.0056 |
| 0.0036 | 0.0036 | 0.9552 | 0.015 | 0.0162 |
| 0.0034 | 0.0044 | 0.8884 | 0.0188 | 0.0788 |
| 0.0164 | 0.0272 | 0.918 | 0.008 | 0.0196 |
| 0.0214 | 0.0138 | 0.9196 | 0.0246 | 0.0018 |
| 0.0326 | 0.1296 | 0.8006 | 0.0156 | 0.0086 |

| | | | | |
|---|---|---|---|---|
| 0.0072 | 0.0332 | 0.8014 | 0.0682 | 0.0176 |
| 0.0084 | 0.0082 | 0.9078 | 0.0098 | 0.0534 |
| 0.0034 | 0.0232 | 0.8418 | 0.0774 | 0.0022 |
| 0.0036 | 0.0034 | 0.8344 | 0.152 | 0.0032 |
| 0.0316 | 0.0252 | 0.8542 | 0.0566 | 0.0204 |
| 0.0352 | 0.023 | 0.832 | 0.0374 | 0.0332 |
| 0.0074 | 0.0462 | 0.8946 | 0.0396 | 0.0028 |
| 0.0026 | 0.0326 | 0.817 | 0.0696 | 0.0046 |
| 0.0802 | 0.0228 | 0.8034 | 0.046 | 0.0072 |
| 0.0622 | 0.0774 | 0.7986 | 0.0186 | 0.0028 |
| 0.011 | 0.0422 | 0.7674 | 0.0898 | 0.0038 |
| 0.0038 | 0.0494 | 0.7282 | 0.1002 | 0.003 |
| 0.0204 | 0.0796 | 0.8442 | 0.0218 | 0.0082 |
| 0.0148 | 0.0478 | 0.8902 | 0.0278 | 0.005 |
| 0.0096 | 0.0122 | 0.8404 | 0.1268 | 0.004 |
| 0.86 | 0.0354 | 0.0034 | 0.0076 | 0.0084 |
| 0.009 | 0.0068 | 0.9486 | 0.0152 | 0.006 |
| 0.3912 | 0.1286 | 0.0092 | 0.0134 | 0.0096 |
| 0.0184 | 0.0612 | 0.704 | 0.1094 | 0.0038 |
| 0.0116 | 0.0074 | 0.8192 | 0.1454 | 0.0032 |
| 0.007 | 0.0472 | 0.8646 | 0.009 | 0.0646 |
| 0.0028 | 0.0038 | 0.9716 | 0.0084 | 0.0094 |
| 0.0056 | 0.069 | 0.7 | 0.0838 | 0.0732 |
| 0.003 | 0.0312 | 0.8274 | 0.0666 | 0.004 |
| 0.0028 | 0.003 | 0.9522 | 0.0294 | 0.009 |
| 0.0338 | 0.0204 | 0.8736 | 0.0298 | 0.006 |
| 0.0298 | 0.032 | 0.8356 | 0.0186 | 0.0656 |
| 0.0584 | 0.0262 | 0.8404 | 0.0354 | 0.0036 |
| 0.0482 | 0.039 | 0.788 | 0.0462 | 0.0424 |
| 0.0304 | 0.0246 | 0.8936 | 0.021 | 0.0038 |
| 0.013 | 0.0256 | 0.8808 | 0.0316 | 0.0028 |
| 0.8412 | 0.093 | 0.0044 | 0.0138 | 0.0108 |
| 0.0068 | 0.0068 | 0.9626 | 0.0078 | 0.008 |
| 0.0156 | 0.0206 | 0.8806 | 0.0384 | 0.023 |
| 0.0084 | 0.0544 | 0.8958 | 0.0332 | 0.0024 |
| 0.0036 | 0.045 | 0.862 | 0.0464 | 0.0054 |
| 0.043 | 0.0154 | 0.8868 | 0.0222 | 0.0042 |
| 0.0278 | 0.0356 | 0.8746 | 0.0132 | 0.0316 |
| 0.0064 | 0.004 | 0.9226 | 0.0582 | 0.0026 |
| 0.0072 | 0.008 | 0.885 | 0.061 | 0.0236 |
| 0.012 | 0.0062 | 0.8986 | 0.019 | 0.0524 |
| 0.0088 | 0.0082 | 0.9474 | 0.0182 | 0.0074 |
| 0.018 | 0.0992 | 0.7376 | 0.0554 | 0.0074 |
| 0.0056 | 0.0042 | 0.9448 | 0.0114 | 0.0274 |

| | | | | |
|---|---|---|---|---|
| 0.0076 | 0.0134 | 0.9562 | 0.0092 | 0.0068 |
| 0.025 | 0.0638 | 0.7878 | 0.0416 | 0.0064 |
| 0.0158 | 0.0366 | 0.8642 | 0.056 | 0.0198 |
| 0.0182 | 0.017 | 0.9016 | 0.034 | 0.0132 |
| 0.0056 | 0.0538 | 0.852 | 0.0152 | 0.0658 |
| 0.0054 | 0.0374 | 0.7876 | 0.0842 | 0.0018 |
| 0.0492 | 0.0132 | 0.8504 | 0.0128 | 0.0258 |
| 0.0424 | 0.0346 | 0.8444 | 0.0398 | 0.0048 |
| 0.0326 | 0.0178 | 0.9178 | 0.0128 | 0.0022 |
| 0.409 | 0.368 | 0.081 | 0.0124 | 0.0796 |
| 0.009 | 0.0666 | 0.8916 | 0.015 | 0.011 |
| 0.0164 | 0.069 | 0.8582 | 0.032 | 0.0044 |
| 0.0376 | 0.0886 | 0.7858 | 0.016 | 0.0458 |
| 0.0206 | 0.0256 | 0.844 | 0.0332 | 0.0568 |
| 0.0188 | 0.039 | 0.8822 | 0.0404 | 0.0028 |
| 0.1716 | 0.0178 | 0.4652 | 0.0684 | 0.003 |
| 0.0064 | 0.0052 | 0.883 | 0.0262 | 0.0722 |
| 0.05 | 0.0324 | 0.8672 | 0.019 | 0.0022 |
| 0.0208 | 0.0448 | 0.8732 | 0.0276 | 0.0036 |
| 0.029 | 0.0276 | 0.8212 | 0.0982 | 0.0028 |
| 0.007 | 0.0108 | 0.7906 | 0.18 | 0.0042 |
| 0.029 | 0.0192 | 0.8726 | 0.0594 | 0.0026 |
| 0.0078 | 0.0364 | 0.8322 | 0.0422 | 0.0744 |
| 0.016 | 0.0146 | 0.8898 | 0.0474 | 0.0106 |
| 0.0094 | 0.0132 | 0.9456 | 0.0148 | 0.0038 |
| 0.0334 | 0.0368 | 0.841 | 0.0284 | 0.0084 |
| 0.0508 | 0.0572 | 0.7878 | 0.0162 | 0.0694 |
| 0.0358 | 0.0932 | 0.82 | 0.026 | 0.0058 |
| 0.0132 | 0.0754 | 0.8802 | 0.012 | 0.0026 |
| 0.0172 | 0.0172 | 0.847 | 0.0716 | 0.034 |
| 0.0052 | 0.0302 | 0.8292 | 0.0626 | 0.01 |
| 0.0072 | 0.0414 | 0.8988 | 0.02 | 0.0142 |
| 0.0042 | 0.0038 | 0.9488 | 0.0332 | 0.0046 |
| 0.1606 | 0.1204 | 0.632 | 0.029 | 0.0098 |
| 0.0086 | 0.0818 | 0.7778 | 0.0664 | 0.0074 |
| 0.0344 | 0.0302 | 0.8504 | 0.0396 | 0.005 |
| 0.052 | 0.0296 | 0.5982 | 0.0972 | 0.1806 |
| 0.006 | 0.0658 | 0.889 | 0.0176 | 0.018 |
| 0.0212 | 0.065 | 0.885 | 0.0108 | 0.004 |
| 0.0142 | 0.0646 | 0.89 | 0.0142 | 0.0028 |
| 0.0342 | 0.0394 | 0.8144 | 0.0822 | 0.0074 |
| 0.2392 | 0.0322 | 0.4616 | 0.0112 | 0.0052 |
| 0.1022 | 0.0644 | 0.5662 | 0.127 | 0.0018 |
| 0.0552 | 0.047 | 0.8486 | 0.0126 | 0.0068 |

| | | | | |
|---|---|---|---|---|
| 0.0058 | 0.0272 | 0.9234 | 0.018 | 0.019 |
| 0.0692 | 0.0148 | 0.808 | 0.0158 | 0.024 |
| 0.0124 | 0.045 | 0.8534 | 0.0538 | 0.0244 |
| 0.0984 | 0.0696 | 0.5992 | 0.145 | 0.0128 |
| 0.0658 | 0.0316 | 0.6312 | 0.114 | 0.1356 |
| 0.0152 | 0.0358 | 0.8726 | 0.0428 | 0.0136 |
| 0.3384 | 0.0968 | 0.013 | 0.1632 | 0.0184 |
| 0.0124 | 0.0148 | 0.8708 | 0.0626 | 0.003 |
| 0.0376 | 0.0336 | 0.8672 | 0.0128 | 0.025 |
| 0.003 | 0.0038 | 0.973 | 0.0154 | 0.0018 |
| 0.0374 | 0.0212 | 0.7476 | 0.1092 | 0.0498 |
| 0.022 | 0.0304 | 0.75 | 0.1562 | 0.0068 |
| 0.0614 | 0.0364 | 0.83 | 0.0236 | 0.004 |
| 0.0076 | 0.0072 | 0.9096 | 0.0664 | 0.0024 |
| 0.0098 | 0.0096 | 0.9364 | 0.0214 | 0.005 |
| 0.1048 | 0.0114 | 0.7778 | 0.0474 | 0.0028 |
| 0.0724 | 0.0792 | 0.8252 | 0.0062 | 0.0022 |
| 0.0258 | 0.0434 | 0.8414 | 0.0176 | 0.0616 |
| 0.7138 | 0.058 | 0.0274 | 0.0264 | 0.0042 |
| 0.004 | 0.0066 | 0.5806 | 0.0084 | 0.397 |
| 0.734 | 0.0446 | 0.1536 | 0.025 | 0.0124 |
| 0.204 | 0.0724 | 0.473 | 0.0312 | 0.0118 |
| 0.1938 | 0.1358 | 0.4008 | 0.043 | 0.0166 |
| 0.7294 | 0.088 | 0.013 | 0.0098 | 0.08 |
| 0.4758 | 0.445 | 0.0084 | 0.0172 | 0.0024 |
| 0.6284 | 0.0456 | 0.009 | 0.0092 | 0.0184 |
| 0.4254 | 0.05 | 0.01 | 0.0174 | 0.0026 |
| 0.4054 | 0.012 | 0.0872 | 0.0356 | 0.2054 |
| 0.471 | 0.1298 | 0.0034 | 0.1046 | 0.0068 |
| 0.3084 | 0.2512 | 0.013 | 0.2104 | 0.004 |
| 0.6068 | 0.0266 | 0.022 | 0.0198 | 0.148 |
| 0.8066 | 0.0414 | 0.0152 | 0.024 | 0.006 |
| 0.0288 | 0.0152 | 0.9098 | 0.0138 | 0.0044 |
| 0.0558 | 0.0228 | 0.8626 | 0.0224 | 0.01 |
| 0.3622 | 0.0128 | 0.3204 | 0.0096 | 0.0036 |
| 0.1564 | 0.0166 | 0.3774 | 0.0118 | 0.1988 |
| 0.5256 | 0.336 | 0.0152 | 0.0218 | 0.0366 |
| 0.4628 | 0.3816 | 0.0284 | 0.0192 | 0.0248 |
| 0.4784 | 0.1178 | 0.0054 | 0.1158 | 0.0064 |
| 0.4014 | 0.0714 | 0.0144 | 0.0412 | 0.0052 |
| 0.3534 | 0.0916 | 0.0896 | 0.0498 | 0.2718 |
| 0.5072 | 0.135 | 0.0738 | 0.0122 | 0.026 |
| 0.4082 | 0.0202 | 0.0152 | 0.0176 | 0.0938 |
| 0.0918 | 0.0316 | 0.2914 | 0.5138 | 0.0094 |

| | | | | |
|---|---|---|---|---|
| 0.0602 | 0.0686 | 0.4298 | 0.0256 | 0.2972 |
| 0.1166 | 0.145 | 0.0114 | 0.1522 | 0.4106 |
| 0.1894 | 0.0188 | 0.0626 | 0.064 | 0.5818 |
| 0.03 | 0.0834 | 0.5932 | 0.1252 | 0.016 |
| 0.3992 | 0.0202 | 0.1058 | 0.0212 | 0.0918 |
| 0.0468 | 0.149 | 0.4238 | 0.101 | 0.196 |
| 0.2632 | 0.0128 | 0.0596 | 0.0618 | 0.245 |
| 0.2974 | 0.0876 | 0.0062 | 0.1454 | 0.2294 |
| 0.2156 | 0.0702 | 0.0344 | 0.1822 | 0.092 |
| 0.0072 | 0.054 | 0.8756 | 0.014 | 0.0362 |
| 0.1672 | 0.022 | 0.5184 | 0.0468 | 0.0584 |
| 0.0492 | 0.123 | 0.2564 | 0.3752 | 0.0142 |
| 0.539 | 0.351 | 0.016 | 0.0138 | 0.01 |
| 0.4298 | 0.1636 | 0.0072 | 0.0392 | 0.2008 |
| 0.319 | 0.0218 | 0.0044 | 0.0064 | 0.1862 |
| 0.2494 | 0.1716 | 0.1336 | 0.0634 | 0.0732 |
| 0.3536 | 0.0666 | 0.073 | 0.0692 | 0.0488 |
| 0.3 | 0.0654 | 0.0142 | 0.034 | 0.4594 |
| 0.3004 | 0.2594 | 0.0192 | 0.027 | 0.3588 |
| 0.2414 | 0.0214 | 0.0166 | 0.036 | 0.315 |
| 0.3542 | 0.0512 | 0.005 | 0.0144 | 0.1702 |
| 0.8208 | 0.0364 | 0.0104 | 0.0132 | 0.071 |
| 0.5876 | 0.202 | 0.0024 | 0.0084 | 0.1512 |
| 0.004 | 0.0052 | 0.3824 | 0.5866 | 0.018 |
| 0.0072 | 0.0122 | 0.3944 | 0.5214 | 0.0586 |
| 0.0494 | 0.0284 | 0.3374 | 0.5468 | 0.0128 |
| 0.0134 | 0.0086 | 0.3796 | 0.5776 | 0.0106 |
| 0.0064 | 0.0052 | 0.3714 | 0.572 | 0.04 |
| 0.0032 | 0.0048 | 0.3702 | 0.5894 | 0.0294 |
| 0.0104 | 0.013 | 0.3118 | 0.571 | 0.0878 |
| 0.014 | 0.0256 | 0.3902 | 0.5572 | 0.0038 |
| 0.0052 | 0.0086 | 0.3922 | 0.5622 | 0.026 |
| 0.0516 | 0.033 | 0.2988 | 0.5314 | 0.0628 |
| 0.0254 | 0.0692 | 0.3022 | 0.525 | 0.059 |
| 0.0058 | 0.0076 | 0.3692 | 0.558 | 0.054 |
| 0.036 | 0.0858 | 0.2612 | 0.5152 | 0.0882 |
| 0.0024 | 0.0236 | 0.324 | 0.5726 | 0.0254 |
| 0.0028 | 0.0046 | 0.319 | 0.5816 | 0.0896 |
| 0.0042 | 0.0054 | 0.4004 | 0.5816 | 0.0044 |
| 0.0164 | 0.0178 | 0.3648 | 0.582 | 0.0058 |
| 0.0056 | 0.0066 | 0.335 | 0.547 | 0.1012 |
| 0.0144 | 0.013 | 0.47 | 0.4916 | 0.0036 |
| 0.053 | 0.0196 | 0.3098 | 0.554 | 0.0382 |
| 0.0054 | 0.0082 | 0.3398 | 0.5576 | 0.0846 |

| | | | | |
|---|---|---|---|---|
| 0.0054 | 0.006 | 0.333 | 0.5766 | 0.0726 |
| 0.0108 | 0.0076 | 0.468 | 0.4932 | 0.0122 |
| 0.0042 | 0.012 | 0.3734 | 0.5532 | 0.0518 |
| 0.4762 | 0.1882 | 0.0126 | 0.085 | 0.1034 |
| 0.5296 | 0.0762 | 0.0046 | 0.0104 | 0.3574 |
| 0.4002 | 0.0176 | 0.0048 | 0.0068 | 0.005 |
| 0.5422 | 0.0842 | 0.0066 | 0.0106 | 0.048 |
| 0.2274 | 0.36 | 0.0356 | 0.0302 | 0.3018 |
| 0.1628 | 0.2542 | 0.0076 | 0.2082 | 0.1944 |
| 0.3998 | 0.0816 | 0.2002 | 0.1634 | 0.0042 |
| 0.25 | 0.263 | 0.0348 | 0.118 | 0.0684 |
| 0.5674 | 0.0926 | 0.0118 | 0.0236 | 0.0162 |
| 0.628 | 0.178 | 0.0072 | 0.0176 | 0.1156 |
| 0.2686 | 0.149 | 0.0186 | 0.172 | 0.098 |
| 0.4104 | 0.0286 | 0.0054 | 0.02 | 0.0032 |
| 0.335 | 0.06 | 0.0146 | 0.1088 | 0.0026 |
| 0.6294 | 0.0388 | 0.004 | 0.0096 | 0.0078 |
| 0.6694 | 0.0754 | 0.0042 | 0.008 | 0.0162 |
| 0.5716 | 0.0364 | 0.0142 | 0.0516 | 0.0076 |
| 0.68 | 0.1522 | 0.0024 | 0.006 | 0.0032 |
| 0.469 | 0.0514 | 0.0068 | 0.0128 | 0.2168 |
| 0.408 | 0.0262 | 0.0044 | 0.009 | 0.2386 |
| 0.418 | 0.3472 | 0.0256 | 0.0284 | 0.0212 |
| 0.3204 | 0.0946 | 0.0226 | 0.1176 | 0.0048 |
| 0.5804 | 0.129 | 0.0072 | 0.0412 | 0.005 |
| 0.0154 | 0.0306 | 0.9286 | 0.0116 | 0.003 |
| 0.326 | 0.1786 | 0.0218 | 0.018 | 0.0426 |
| 0.3784 | 0.1386 | 0.0218 | 0.0564 | 0.0356 |
| 0.613 | 0.1654 | 0.0022 | 0.0046 | 0.0186 |
| 0.5822 | 0.1302 | 0.0128 | 0.0206 | 0.0078 |
| 0.223 | 0.2784 | 0.1986 | 0.2666 | 0.0188 |
| 0.465 | 0.4202 | 0.0252 | 0.0118 | 0.009 |
| 0.4822 | 0.3672 | 0.0454 | 0.0162 | 0.0058 |
| 0.443 | 0.0468 | 0.129 | 0.2008 | 0.005 |
| 0.1708 | 0.1102 | 0.141 | 0.3762 | 0.0026 |
| 0.512 | 0.2436 | 0.0994 | 0.0408 | 0.0592 |
| 0.5146 | 0.3166 | 0.0482 | 0.0684 | 0.002 |
| 0.2984 | 0.1332 | 0.1188 | 0.0498 | 0.1304 |
| 0.4402 | 0.0636 | 0.1578 | 0.0526 | 0.0858 |
| 0.7916 | 0.0666 | 0.0052 | 0.016 | 0.081 |
| 0.489 | 0.0704 | 0.022 | 0.0122 | 0.1536 |
| 0.031 | 0.0448 | 0.3504 | 0.5246 | 0.022 |
| 0.1426 | 0.0128 | 0.5714 | 0.094 | 0.0918 |
| 0.3032 | 0.1884 | 0.2122 | 0.0178 | 0.252 |

| | | | | |
|---|---|---|---|---|
| 0.414 | 0.3114 | 0.0322 | 0.1334 | 0.004 |
| 0.5298 | 0.2 | 0.0376 | 0.0192 | 0.1238 |
| 0.3588 | 0.0724 | 0.1776 | 0.3222 | 0.0186 |
| 0.0024 | 0.002 | 0.4904 | 0.4974 | 0.0052 |
| 0.067 | 0.0378 | 0.401 | 0.1898 | 0.1752 |
| 0.0546 | 0.0054 | 0.3378 | 0.5322 | 0.0348 |
| 0.0038 | 0.0064 | 0.3816 | 0.5596 | 0.0422 |
| 0.004 | 0.0122 | 0.37 | 0.5548 | 0.055 |
| 0.0802 | 0.0058 | 0.722 | 0.0478 | 0.0132 |
| 0.1226 | 0.053 | 0.3174 | 0.4176 | 0.0536 |
| 0.0086 | 0.0064 | 0.3712 | 0.5912 | 0.0134 |
| 0.003 | 0.002 | 0.3572 | 0.5828 | 0.0512 |
| 0.0072 | 0.006 | 0.3804 | 0.584 | 0.0152 |
| 0.2694 | 0.3302 | 0.1622 | 0.2106 | 0.0188 |
| 0.3822 | 0.0352 | 0.1488 | 0.2788 | 0.0234 |
| 0.0414 | 0.023 | 0.3914 | 0.513 | 0.014 |
| 0.0036 | 0.0042 | 0.4086 | 0.564 | 0.0156 |
| 0.318 | 0.1772 | 0.2948 | 0.1662 | 0.0218 |
| 0.1764 | 0.0736 | 0.4324 | 0.132 | 0.0644 |
| 0.2794 | 0.19 | 0.0572 | 0.081 | 0.2972 |
| 0.5056 | 0.0284 | 0.0132 | 0.0082 | 0.097 |
| 0.9068 | 0.049 | 0.007 | 0.007 | 0.0026 |
| 0.3744 | 0.0594 | 0.0128 | 0.0106 | 0.0906 |
| 0.3536 | 0.1754 | 0.0406 | 0.0848 | 0.0112 |
| 0.49 | 0.444 | 0.0228 | 0.0068 | 0.0022 |
| 0.5068 | 0.025 | 0.0158 | 0.0234 | 0.0028 |
| 0.514 | 0.2006 | 0.1978 | 0.0284 | 0.0108 |
| 0.327 | 0.2502 | 0.0144 | 0.027 | 0.2 |
| 0.4214 | 0.0476 | 0.0208 | 0.0596 | 0.0166 |
| 0.6144 | 0.0536 | 0.0376 | 0.066 | 0.0734 |
| 0.413 | 0.1796 | 0.0042 | 0.0252 | 0.1724 |
| 0.5818 | 0.2096 | 0.037 | 0.0182 | 0.0186 |
| 0.1152 | 0.0106 | 0.0532 | 0.0886 | 0.538 |
| 0.5364 | 0.1436 | 0.009 | 0.0152 | 0.2714 |
| 0.4214 | 0.0284 | 0.0454 | 0.0192 | 0.0096 |
| 0.4128 | 0.0892 | 0.0304 | 0.032 | 0.0322 |
| 0.4544 | 0.0156 | 0.0064 | 0.0252 | 0.3098 |
| 0.759 | 0.06 | 0.0118 | 0.0214 | 0.0142 |
| 0.2868 | 0.3718 | 0.1866 | 0.011 | 0.02 |
| 0.3732 | 0.0112 | 0.0242 | 0.0776 | 0.0094 |
| 0.5132 | 0.3734 | 0.0214 | 0.0152 | 0.003 |
| 0.6746 | 0.0384 | 0.0218 | 0.0356 | 0.0382 |
| 0.5166 | 0.1054 | 0.0444 | 0.0082 | 0.003 |
| 0.6634 | 0.0586 | 0.118 | 0.023 | 0.0934 |

| | | | | |
|---|---|---|---|---|
| 0.4098 | 0.4282 | 0.0386 | 0.0448 | 0.0148 |
| 0.6658 | 0.0648 | 0.0038 | 0.0062 | 0.0218 |
| 0.325 | 0.0154 | 0.2128 | 0.0216 | 0.0022 |
| 0.5772 | 0.0772 | 0.025 | 0.105 | 0.0968 |
| 0.3712 | 0.0148 | 0.1052 | 0.0156 | 0.0056 |
| 0.663 | 0.1134 | 0.1088 | 0.027 | 0.0098 |
| 0.2814 | 0.069 | 0.0518 | 0.0498 | 0.0648 |
| 0.4262 | 0.0376 | 0.0062 | 0.0148 | 0.0022 |
| 0.7452 | 0.0148 | 0.0104 | 0.0154 | 0.0346 |
| 0.699 | 0.0522 | 0.02 | 0.0276 | 0.049 |
| 0.2352 | 0.2948 | 0.043 | 0.0082 | 0.3754 |
| 0.3362 | 0.0622 | 0.0282 | 0.0426 | 0.3382 |
| 0.0082 | 0.0078 | 0.4088 | 0.5242 | 0.0426 |
| 0.1322 | 0.0086 | 0.3084 | 0.4806 | 0.0358 |
| 0.218 | 0.0276 | 0.3952 | 0.0726 | 0.0056 |
| 0.0852 | 0.046 | 0.4228 | 0.0762 | 0.3002 |
| 0.004 | 0.0042 | 0.3474 | 0.5684 | 0.072 |
| 0.0064 | 0.0086 | 0.3736 | 0.581 | 0.0246 |
| 0.0122 | 0.0124 | 0.4108 | 0.545 | 0.0084 |
| 0.343 | 0.009 | 0.0036 | 0.0112 | 0.5376 |
| 0.5026 | 0.1018 | 0.0104 | 0.0064 | 0.336 |
| 0.8814 | 0.0148 | 0.003 | 0.0098 | 0.002 |
| 0.0432 | 0.0116 | 0.3536 | 0.537 | 0.0318 |
| 0.0044 | 0.058 | 0.333 | 0.531 | 0.023 |
| 0.0068 | 0.008 | 0.3604 | 0.5706 | 0.043 |
| 0.0116 | 0.0092 | 0.3514 | 0.5922 | 0.0306 |
| 0.3886 | 0.233 | 0.0052 | 0.124 | 0.0312 |
| 0.0102 | 0.0092 | 0.3696 | 0.572 | 0.031 |
| 0.009 | 0.0326 | 0.3342 | 0.5658 | 0.0408 |
| 0.0464 | 0.059 | 0.3148 | 0.5054 | 0.0362 |
| 0.0358 | 0.0398 | 0.3362 | 0.5466 | 0.017 |
| 0.2628 | 0.2198 | 0.1234 | 0.0356 | 0.1744 |
| 0.616 | 0.141 | 0.0288 | 0.03 | 0.0032 |
